# Multilevel feedback architecture for adaptive regulation of learning in the insect brain

**DOI:** 10.1101/649731

**Authors:** Claire Eschbach, Akira Fushiki, Michael Winding, Casey M. Schneider-Mizell, Mei Shao, Rebecca Arruda, Katharina Eichler, Javier Valdes-Aleman, Tomoko Ohyama, Andreas S. Thum, Bertram Gerber, Richard D. Fetter, James W. Truman, Ashok Litwin-Kumar, Albert Cardona, Marta Zlatic

**Author notes:** co-first authors (Claire Eschbach, Akira Fushiki). co-second authors (Michael Winding, Casey M. Schneider-Mizell, Mei Shao).

## Abstract

Modulatory (*e.g*. dopaminergic) neurons provide “teaching signals” that drive associative learning across the animal kingdom, but the circuits that regulate their activity and compute teaching signals are still poorly understood. We provide the first synaptic-resolution connectome of the circuitry upstream of all modulatory neurons in a brain center for associative learning, the mushroom body (MB) of the *Drosophila* larva. We discovered afferent pathways from sensory neurons, as well as an unexpected large population of 61 feedback neuron pairs that provide one- and two-step feedback from MB output neurons. The majority of these feedback pathways link distinct memory systems (*e.g*. aversive and appetitive). We functionally confirmed some of the structural pathways and found that some modulatory neurons compare inhibitory input from their own compartment and excitatory input from compartments of opposite valence, enabling them to compute integrated common-currency predicted values across aversive and appetitive memory systems. This architecture suggests that the MB functions as an interconnected ensemble during learning and that distinct types of previously formed memories can regulate future learning about a stimulus. We developed a model of the circuit constrained by the connectome and by the functional data which revealed that the newly discovered architectural motifs, namely the multilevel feedback architecture and the extensive cross-compartment connections, increase the computational performance and flexibility on learning tasks. Together our study provides the most detailed view to date of a recurrent brain circuit that computes teaching signals and provides insights into the architectural motifs that support reinforcement learning in a biological system.

## Introduction

To behave adaptively in an ever-changing environment, animals must not only be able to learn new associations between conditioned stimuli (CS) and rewards or punishments (in Pavlovian terms, aversive and appetitive unconditioned stimuli, US), but also continuously update previous memories, depending on their relevance and reliability^1–14^. For example, memories can be consolidated into a persistent form and maintained^5,6,8,15–17^, extinguished^5,7,9,11,14,18^, or expanded and combined into chains of associations (as in higher-order conditioning^7,19,20^). Furthermore, learning can itself be flexible and depends both on present context and past history^14,21,22^. These are fundamental brain functions across the animal kingdom, but the learning algorithms used by brains and their circuit implementations are still unclear.

Modulatory neurons (*e.g*. dopaminergic, DANs) convey information about rewards and punishments and provide the so-called teaching signals for updating the valence associated with CS in learning circuits across the animal kingdom (*e.g*. in the vertebrate basal ganglia, or in the insect mushroom body, MB)^12,14,21,23–25^. In the simplest models of associative learning, learning is driven by correlations between CS and US, and modulatory neuron activity represents just the received US^26,27^. To account for more complex behavioral phenomena, theories have been developed in which learning can be regulated by previously formed associations and modulatory neuron responses to CS are adaptively modified by prior learning^4,12,21,28–36^. For example, in reinforcement learning, learning is driven by errors between predicted and actual US (so-called prediction errors)^21,28–35^, which are represented by the activity of modulatory neurons. Indeed, responses of many modulatory neurons have been shown to be adaptive, in monkeys^21,31,32,34,37^, rodents^12,25,36,38–41^, and insects^13,23,42–44^, although the extent to which this is the case in insects has been less extensively investigated. Despite recent progress^12–14,41,45–49^, the basic principles by which modulatory neuron activity is adaptively regulated, and what teaching signals they compute and encode, are not well understood.

A prerequisite for the adaptive regulation of modulatory neuron activity is convergence of afferent pathways that convey information about received rewards and punishments^12,21^ with feedback pathways that convey information about previous experience. In the rodent basal ganglia, DANs have indeed been shown to receive feedback from striatum and striatal neurons have been implicated in prediction error computations^12,39,50–52^. In the *Drosophila* mushroom body (MB), some DANs have also been shown to receive direct feedback input from MB output neurons and specific output neurons have been implicated in memory updating^13,14,46–49,53–55^. However, we still know very little about the nature of these feedback circuits and the way in which they compute features such as predictions and prediction errors. How much input do modulatory neurons receive from afferent vs. feedback pathways? Do these pathways converge on modulatory neurons themselves, or at multiple levels upstream? Distinct types of memories (*e.g*. aversive and appetitive, short-term and long-term) are often formed in distinct locations, in both flies^14,24,56–58^ and rodents^59–61^, but are there feedback pathways that enable memories of one type to influence the formation of memories of a different type? How are integrated common-currency predictions across aversive and appetitive memory systems computed? How many distinct types of feedback motifs are there and what computational advantages do they offer? Addressing these questions is essential for understanding how learning algorithms are implemented in neural circuits. However, such a comprehensive characterization of feedback pathways requires a synaptic-resolution connectivity map of the complete set of modulatory neurons, their target output neurons, and of all of their pre- and post-synaptic partners, which has previously been out of reach.

Insects, especially their larval stages, have small and compact brains that have recently become amenable to large-scale electron microscopy (EM) circuit mapping^47,62^. Both adult^8,14,27,48,63–67^ and larval^47,57,58,68–70^ insect stages possess a brain center essential for associative learning, the MB. The MB contains parallel fiber neurons called Kenyon Cells (KCs) that sparsely encode CS^66,71–74^; MB modulatory neurons (collectively called MBINs) that provide the teaching signals for updat_ing the valence associated with CS_^13,14,42,46,47,55–58,74–84^; and MB output neurons (MBONs) whose activity represents learnt valences of stimuli^14,58,66,85–93^. Most modulatory neurons are dopaminergic (we called this subset DANs, adding a letter that indicates their target compartment in the MB, *e.g*. DAN-g1), some are octopaminergic (OANs, *e.g*. OAN-g1), and some have unidentified neurotransmitters (so we refer to this subset by their generic name, *e.g*. MBIN-e1). Modulatory neurons and MBONs project axon terminals and dendrites, respectively, onto the KC axons in a tiled manner, defining MB compartments, in both adult^14,45,48,66,75,94^ and larval^47,58,95^ *Drosophila*. In adult *Drosophila*, it has been shown that co-activation of KCs and DANs reduces the strength of the KC-MBON synapse in that compartment^14,55,66,85,90,91,96,97^. Different compartments have been implicated in the formation of distinct types of memories, for example aversive and appetitive, or short- and long-term^14,24,56–58,66,74,76,81,83,84,89–91,98–100^. However, the extent to which learning in a specific compartment is regulated by the output from its own compartment or from other compartments is still unclear. A few direct anatomical connections from MBONs to modulatory neurons have been identified^13,14,45–49,53,54^, but possible indirect connections via intermediate feedback neurons have not been investigated. In total, despite a good understanding of the structure and function of the core components of the MB^14,47,48,58^, the circuits presynaptic to modulatory neurons that regulate their activity have remained largely uncharacterized, in both adult and larval *Drosophila*.

We therefore reconstructed all neurons upstream of all modulatory neurons in an EM volume that spans the entire nervous system of a 1st instar *Drosophila* larva, in which we had previously reconstructed all the core components of the MB (including 145 KCs, 48 MBONs and 28 modulatory neurons)^47^. Working in the same EM volume enabled us to not only identify all the neuron types upstream of the modulatory neurons, but also to precisely determine which MBONs they receive input from. The present EM reconstruction effort was even larger than reconstructing the core components, because of the large number of neuron types presynaptic to modulatory neurons. We reconstructed a total of 431 previously unknown neurons, of which 102 pairs of homologous neurons (*i.e*. present in each brain hemisphere) make at least 3 synaptic connections onto the modulatory neurons. We also determined which individual modulatory neurons are activated by punishments and reconstructed their afferent US pathways from nociceptive and mechanosensory neurons. We characterized the neurotransmitter profiles of some of the neurons in the network and functionally confirmed some of the identified structural connections. Finally, we developed a model of the circuit constrained by the connectome, the neurotransmitter data, and the functional data and used it to explore the computational advantages offered by the newly discovered architectural motifs for performing distinct learning tasks.

Surprisingly, we found that the majority of neuron types (61 out of 102) presynaptic to modulatory neurons provide one-step or two-step feedback from MBONs. Furthermore, modulatory neurons received extensive input not only from MBONs in their own compartment, but also from many other compartments. In our model the multilevel and cross-compartment feedback architecture improves computational performance on learning tasks that rely on the adaptation of modulatory neuron responses. These pathways may therefore form the neural architecture that permits previously formed associations to instruct future learning, a critical computation in brain circuits that implement reinforcement learning algorithms.

## Results

### Larval MB modulatory neurons for aversive and appetitive memory formation

To more easily interpret the circuitry for regulating modulatory neuron activity and to better constrain models of the circuit, we explored the functional diversity of larval modulatory neurons and identified individual compartments of the larval MB whose modulatory neuron activation paired with odor can evoke aversive or appetitive memory. Previous studies have already shown that pairing of an odor with activation of all, or individual DANs, which target the MB medial lobe (ML), induces appetitive memory^47,57,58^. Another study has shown that pairing an odor with the activation of all DANs that target the vertical lobe (VL), the lateral appendix (LA), and the peduncle (P) (with a broadly-expressing TH-GAL4 driver line) induces aversive memory^56^. To disentangle the role of individual modulatory neuron types in aversive learning, we generated Split-GAL4 lines^101,102^ that drive expression selectively in one or two modulatory neurons per hemisphere (Fig. 1a, Extended Data Fig. 1, Supplementary Table 1). We then paired an odor (CS) with Chrimson-mediated optogenetic activation^103^ of these modulatory neurons in a three-trial, one-odor, associative memory paradigm (Fig. 1b). Because we tested larvae immediately after the last training trial, and less than 20 min after the first training trial, we assume the test reveals mainly short-term memory^70^.

**Figure 1.**
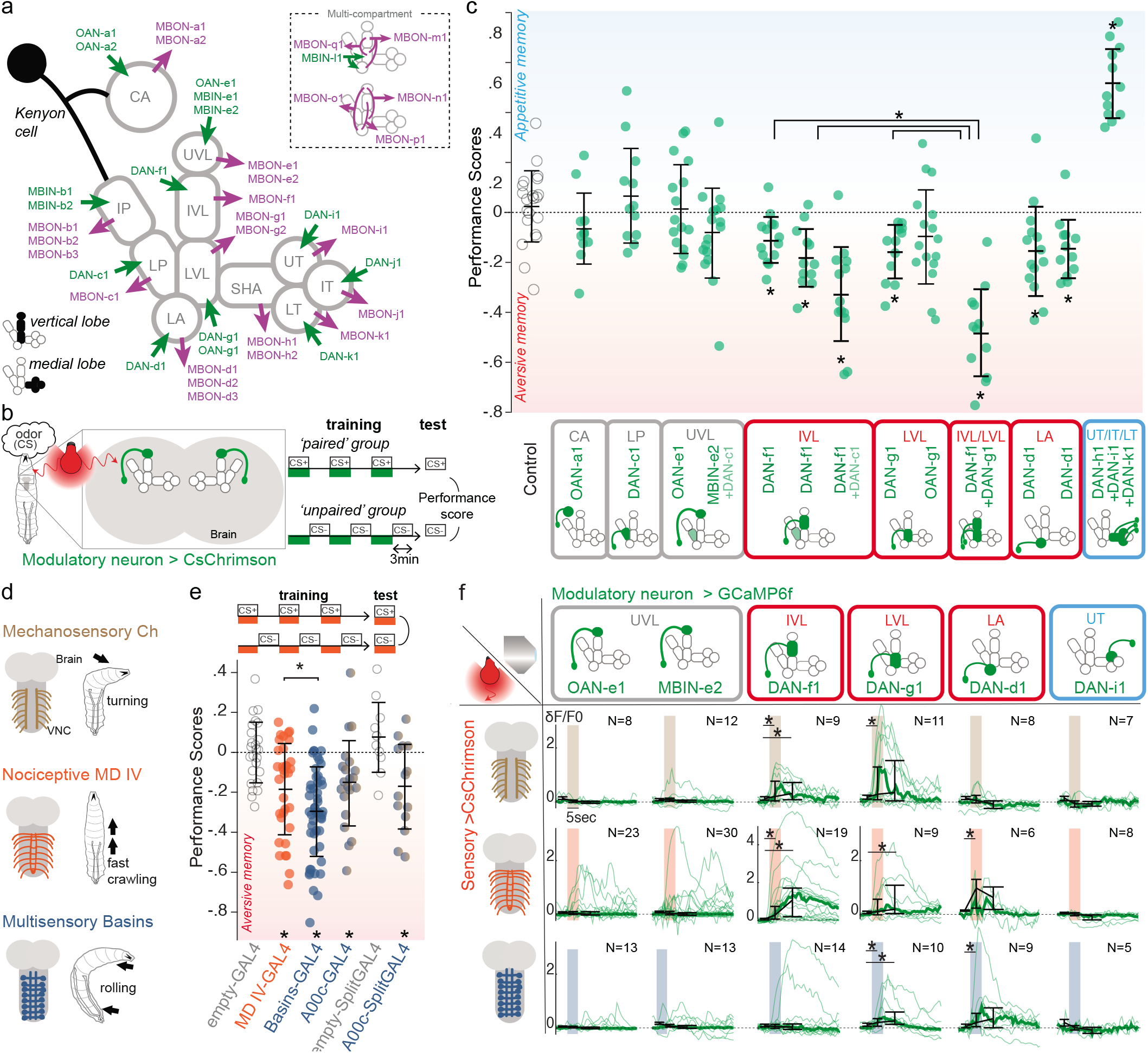
Individual vertical lobe DANs can induce aversive memory and represent different kinds of punishments in *Drosophila* larva. **a** Schematic diagram of the larval mushroom body (MB) compartments with an example Kenyon cell (*black*), all Mushroom Body Output Neurons (MBONs, *purple*) and modulatory neurons (*green*), named based on neurotransmitter expression and compartmental localisation: dopaminergic neurons (DANs), octopaminergic (OANs), or unknown (MBINs). CA: calyx; LP, IP: lower and intermediate peduncle; LA: lateral appendix; VL: vertical lobe; UVL, IVL, LVL: upper, intermediate and lower vertical lobe; SHA, UT, IT, LT: shaft, upper, intermediate and lower toe of the medial lobe. **b** Schematic diagram of the one-odor associative memory optogenetic training protocol consisting of three training trails (3 min each). Larvae from “paired” groups are trained to associate an odor (ethyl acetate) with optogenetic activation (red light) of a pair of modulatory neurons expressing CsChrimson. Larvae from “unpaired” groups undergo bouts of odor presentation intercalated with bouts of modulatory neuron activation. A preference index (*PI*) was computed for each group: *PI = [N(larvae on the odor side) - N(larvae on the no-odor side)] /N(total*). The learning performance score (*LPS*) was then computed: *LPS = [PI(paired) - PI(unpaired)] /2*. Positive learning performance scores result from a higher fraction of larvae on the odor side in the paired group than in the unpaired group, indicating appetitive memory; negative learning performance scores indicate aversive memory. **c** Activation of vertical or medial lobe DANs induces memories of opposite valence. Figure shows learning performance scores obtained with CsChrimson-mediated activation of one or a few pair(s) of modulatory neurons using Split-GAL4 in a one-odor associative memory protocol: OAN-a1 (*SS24765*, n=11), DAN-c1 (*SS02160*, n=12), OAN-e1 (*SS01958*, n=16), combined MBIN-e2 and DAN-c1 (*SS01702*, n=18), DAN-f1 (*MB145B*, n=14 and *SS02180*, n=12), combined DAN-f1 and DAN-c1 (*MB065B*, n=14), DAN-g1 (*SS01716*, n=12), OAN-g1 (*SS04268*, n=16), combined DAN-f1 and DAN-g1 (*MB054B*, n=12), DAN-d1 (*MB143B*, n=14 and *MB328B*, n=12), combined DAN-h1, DAN-i1, DAN-k1 (*SS01948*, n=12), control (*w;attP40;attP2*, n=21). Pairing of odor with optogenetic activation of individual DAN-f1, -g1 and -d1 in VL or LA induces aversive memory. In contrast, pairing of odor with the combined optogenetic activation of DAN-h1, -i1 and -k1 in the ML induces appetitive memory. MB compartments for aversive memory (negative performance scores) are outlined in *red*; compartments for appetitive memory (positive performance scores) are outlined in *blue*. Each data point represents the learning performance score for one reciprocal experiment involving one “paired” and one “unpaired” group. Mean and standard deviations are plotted. *: significant difference from the scores obtained in the control group (*open circles*, Mann-Whitney U test P-values compared to threshold of 0.05 adjusted with a Holm-Bonferroni correction for multiple comparisons). **d** Different types of somatosensory neurons induce distinct types of innate escape responses. Optogenetic or thermogenetic activation of mechanosensory neurons (chordotonal, Ch, *brown*), nociceptive multidendritic class IV neurons (MD IV, *orange*), or multisensory interneurons (Basins, *blue*) induces turning, fast forward crawling or rolling, respectively (Ohyama *et al*. 2013, 2015, Jovanic et al. 2016). **e** Optogenetic activation of nociceptive MD IV (*ppk1.9-GAL4*, n=33), Basins (*GMR72F11-GAL4*, n=52), or the ascending neuron A00c (postsynaptic to Basins, *GMR71A10-GAL4,ppk-GAL80,repo-GAL80*, n=21 or *SS00883-Split-GAL4*, n=14) induces aversive memory when paired with odor (ethyl acetate). Learning performance scores were computed as described in **b**. The larvae were tested under constant optogenetic activation, as depicted in the schematic diagram, to ensure the expression of aversive memory (See *Extended Data Fig. 2* for details). Mean and standard deviations are plotted, as well as significant differences from the scores of the respective control group (*w;;attP2*, n=25 or *w;attP40;attP2*, n=10, according to the driver line, *open circles*), in the same way as in **c**. **f** DANs whose activation paired with odor induces aversive memory respond consistently to optogenetic activation of somatosensory neurons. DAN-f1 preferentially responds to Ch and MD IV stimulations, DAN-d1 to MD IV and Basins stimulations, and DAN-g1 responds to all three types of stimulation. Plots show calcium transients in selected modulatory neurons evoked by optogenetic activation of Ch (*iav-LexA>LexAop-CsChrimson, top* row), MD IV (*ppk1.9-LexA> LexAop-CsChrimson, middle row*) and Basin (*GMR72F11-LexA>LexAop-CsChrimson, bottom row*) neurons. GCaMP6f was expressed in OAN-e1 (SS01958), MBIN-e1 (*SS01702*), DAN-f1 (*MB145b*), DAN-g1 (SS01716), DAN-d1 (*MB143b*), or DAN-i1 (SS00864) using *Split-GAL4*. In each plot, *thin lines* are the averaged responses for one brain, from 3 repeats (triplet trial); *thick lines* are the median across all animals tested. Black plots indicate the median peak *δ*F/F0 of the individual curves in the following time windows: 1 sec before, 1 sec during, and 2 sec following the stimulation. Errorbars show the 25th and 75th percentile of peak *δ*F/F0 of the individual curves. *, P<0.05 Mann-Whitney U-test. Outline color corresponds to the type of memory induced by the neuron, as described in **c**.

We found that pairing an odor with the activation of DAN-f1 (projecting to the intermediate vertical lobe, IVL), DAN-g1 (projecting to lower vertical lobe, LVL), or DAN-d1 (projecting to LA) established aversive memory (Fig. 1c and Extended Data Fig. 2a). In contrast, and as previously reported^47,57,58^, pairing an odor with the activation of DANs that project to the ML (DAN-h1, DAN-i1, and DAN-k1) led to the formation of an appetitive memory (Fig. 1c and Extended Data Fig. 2a). Thus, similar to findings in the adult fly^14,24,66,74,76–78,84,104,105^, larval DANs that innervate distinct lobes are functionally distinct from each other, in that their activation signals opposite valences. Activation of larval PAM-cluster DANs that innervate the ML signals positive valence, whereas activation of larval DL-cluster^95^ DANs that innervate the VL and the LA signals negative valence. Our results also suggest, in accordance with other studies^7,8,13,106^, that presenting an odor unpaired with the activation of some of these DANs induces memory of opposite valence to the paired presentation (Extended Data Fig. 2b).

We found that pairing of an odor with the activation of DAN-c1 that projects to lower peduncle (LP) induced neither appetitive nor aversive memory (Fig. 1c and Extended Data Fig. 2a). Similarly, no memory was induced by pairing an odor with activation of any OANs, or of MBIN-e2 (which was immunonegative for dopamine, octopamine, acetlylcholine, GABA, and glutamate^47^). Thus, DAN-c1, OANs, and MBIN-e2 appear to be functionally distinct from the lobe DANs (with a possible caveat that the GAL4 lines for these neurons may be weaker than the ones for lobe DANs). What role, if any, these neurons play in learning remains to be uncovered in the larva (for some roles of the OAN system see^42,56,79,107^). In any case our analysis has revealed at least three functionally distinct classes of compartments in the larval MB: ML compartments whose DANs can induce appetitive memory when their activation is paired with odor; LA, LVL and IVL compartments whose DANs can induce aversive memory when their activation is paired with odor; and others whose modulatory neurons were not sufficient to induce memory (Fig. 1c).

### Punishment encoding across larval MB modulatory neurons

Next, we asked whether there is any functional diversity within the population of VL/LA DANs whose activation signals punishment. In principle, multiple DANs that project to distinct VL/LA compartments could represent a functionally uniform population and redundantly signal any type of aversive US, as proposed for DANs that signal reward in the mammalian ventral tegmental area^12,40^. Alternatively, distinct DANs could signal distinct types of aversive US, as proposed for DANs that signal rewards in *Drosophila* larva and adult^58,74,78,79,82,83,108,109^_. In the_ adult, the same DANs can convey the teaching signals for different aversive stimuli^77,81,104,110–112^, and respond to multiple aversive stimuli^75,110–113^, but some DANs also appear to be preferentially tuned to some aversive stimuli, but not others^75,111,112^. The extent to which individual DANs that signal aversive stimuli are functionally diverse and whether and how punishment quality or punishment salience may be encoded by DANs in *Drosophila* are therefore open questions.

Larvae sense multiple types of innately aversive somatosensory stimuli that evoke distinct types of innate responses^62,114–124^ (Fig. 1d). Vibration or optogenetic activation of the vibration-sensing mechanosensory neurons evokes hunching (startle) and turning (avoidance)^62,114,119,121,123,124^. Optogenetic activation of nociceptive neurons evokes a more vigorous escape response: fast crawling^62,116,119,123^. Wasp attack that stimulates both nociceptive and mechanosensory neurons, or optogenetic activation of Basin interneurons that integrate mechanosensory and nociceptive inputs, evokes the most vigorous and fastest escape response: rolling^62,115,119,122,125^. Already the mildest of these punishments, vibration, induces aversive associative memory in an olfactory learning paradigm^126^. Fittingly, we found that the stronger forms of punishment, namely, optogenetic activation of the nociceptive sensory neurons, Basins, or the A00c neurons that are directly downstream of Basins, also induce aversive associative memory when paired with odor (Fig. 1e and Extended Data Fig. 2a). We therefore asked how individual modulatory neurons respond to each type of punishment administered by optogenetic activation of specific somatosensory-related neurons, by monitoring their calcium transients (using GCaMP6f^127^) (Fig. 1f). We performed these experiments in isolated nervous systems to avoid any movement-related responses^55^.

In each of the VL/LA-DANs whose activation paired with odor induced aversive memory, we found reliable responses to at least two of the three fictive punishment types. Each punishment type evoked reliable and statistically significant responses in at least two out of three VL/LA-DANs (Fig. 1f). Nevertheless, individual VL/LA-DANs differed in terms of which punishment types evoked reliable and statistically significant responses (Fig. 1f). Specifically, DAN-g1 responded non-selectively to all three punishment types, DAN-f1 responded preferentially to mechanosensory and nociceptive neuron activation, and DAN-d1 to nociceptive and Basin neuron activation. Therefore, considering one of these DANs alone would not allow decoding punishment type, but considering all three would. Thus, the three VL/LA DANs could combinatorially encode punishment type or punishment salience.

For comparison, we also tested the response of a few modulatory neurons whose activation paired with odor did not induce aversive memory. As expected, we found that DAN-i1 (projecting to ML) whose activation paired with odor induced appetitive memory was not activated by the fictive punishments (Fig. 1f). Modulatory neurons that project to UVL (OAN-e1 and MBIN-e2) and whose activation paired with odor induced neither aversive nor appetitive memory were not significantly activated by the fictive punishments, although we observed occasional responses to nociceptive neuron activation (Fig. 1f).

### Comprehensive EM reconstruction of all input neurons to larval MB modulatory neurons

To provide a basis for understanding how the activity and function of modulatory neurons is regulated we sought to comprehensively identify all the neurons presynaptic to them. We have previously reconstructed all of the KCs, olfactory projection neurons (PNs), MBONs, and modulatory neurons in an EM volume of a 1st instar larval nervous system^47^. Here we systematically reconstructed all neurons presynaptic to all modulatory neurons in the same EM volume (Fig. 2a-e). We identified 213 left-right homologous pairs and 5 unpaired neurons presynaptic to modulatory neurons (that we collectively called pre-modulatory neurons). Out of these, we consider 102 homologous pairs to be reliably connected, since both the left and right homologous connections have at least 3 synapses and their sum is at least 10 (thresholds were chosen to make the likelihood of a false positive connection extremely small^62,128^, Fig. 2a-b and e, Supplementary Adjacency Matrix, Supplementary Atlas, Materials and Methods). We refer to the remaining pre-modulatory neurons as “other weakly connected partners”. While they could also influence modulatory neuron activity, especially in combination with each other, we focus our study mainly on the reliably and numerically more strongly connected 102 pre-modulatory neuron pairs.

**Figure 2.**
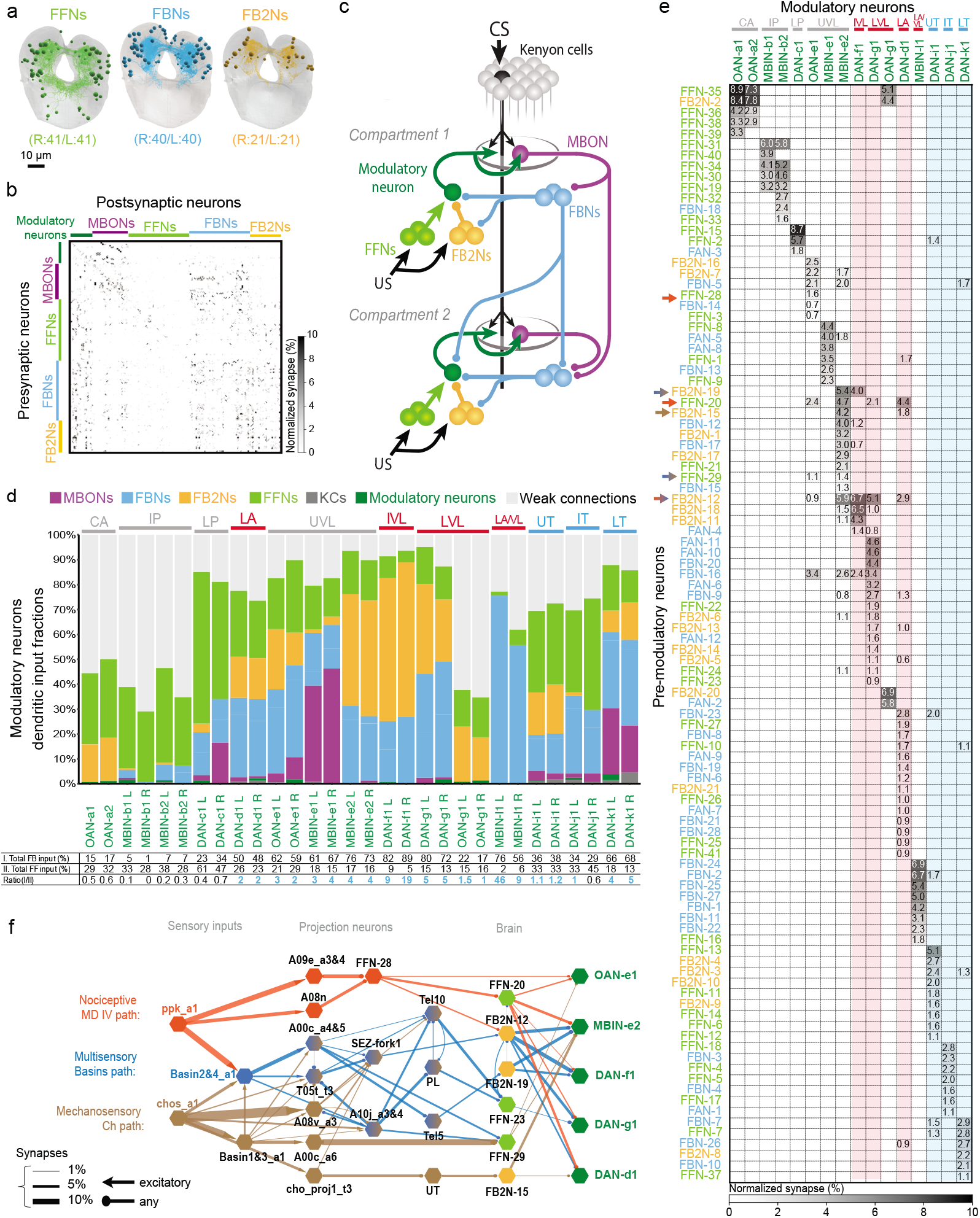
Comprehensive EM reconstruction of pre-modulatory neurons reveals a multilayered recurrent architecture for regulating learning. **a-c** Comprehensive EM reconstruction of all neurons presynaptic to all modulatory neurons reveals a large previoulsy unknown population of feedback neurons. **a**, The newly discovered 102 neuron pairs presynaptic to the modulatory neurons. Of these, the majority (61) relays inputs from mushroom body output neurons (MBONs): 40 first-order feedback neuron pairs (FBNs, *light blue*) receive direct input from MBONs and provide one-step feedback to modulatory neurons; and 21 second-order feedback neuron pairs (FB2Ns, *yellow*) receive one-step MBON input via FBNs and provide two-step feedback to modulatory neurons. The remaining 41 are tentatively classified as feedforward neurons (FFNs, *light green*) that may represent unconditioned stimuli (US). Projections of EM reconstructions of all neurons within each category are shown. The complete set of neurons presynaptic to modulatory neurons amounts to 211 left-right pairs and 5 bilaterally projecting unpaired neurons (not shown). Of these, only 204 neurons (102 pairs) make at least 3 synapses onto any particular modulatory neuron and together as left-right pairs make at least 10 synapses onto any particular left-right pair of homologous modulatory neurons, and are therefore considered reliably and strongly connected. **b**, Connectivity matrix showing normalized synaptic input (expressed as % input) each postsynaptic (*columns*) neuron receives from each presynaptic (*rows*) neuron in the extended MB circuit, comprising modulatory neurons (*green*), MBONs (*purple*), and the different types of pre-modulatory neurons. The normalization is done by computing the % of total input a postsynaptic neuron receives from a presynaptic neuron *i.e*. by dividing the number of synapses from a presynaptic neuron to a postsynaptic neuron by the total number of input synapses of that postsynaptic neuron and multiplying by 100. The average for left and right homologs is shown. Only reliable connections for which both the left and right homologous connections have at least 3 synapses and their sum is at least 10 are shown. Note that among the pre-modulatory neurons, FBNs (*blue*) and FB2Ns (*yellow*) are more interconnected to each other than to FFNs (*light green*). **c**, Layers of feedback neurons reveal the highly recurrent architecture of the circuits. Figure shows a schematic wiring diagram of the core components of the MB together with the newly discovered components upstream of modulatory neurons and downstream of MBONs. Most modulatory neurons receive one-step and two-step within and cross-compartment feedback. **d** Fraction of total dendritic input each modulatory neuron receives from MBONs, FBNs, FB2Ns, FFNs, and from other weakly connected partners (i.e. those that make less than 3 synapses onto a modulatory neuron and less than 10 synapses onto any left-right pair of homologous modulatory neurons). Many modulatory neurons, including most of the DANs, receive more than half of their total dendritic input from the feedback pathways (i.e. from MBONs, FBNs, and FB2Ns), indicating the likely importance of these pathways for governing modulatory neuron activity and thereby learning. The OANs in the calyx (CA), intermediate peduncle (IP) and low vertical lobe (LVL), are drastically different from other modulatory neurons in that they receive a large fraction of input from many weakly connected partners (i.e. neurons that connect to modulatory neurons with very few synapses as explained in a), and relatively little feedback from MBONs. These OANs may therefore encode very different features from the rest of the modulatory neurons. Some DANs extend their dendritic arbors to the KCs which accounts for a small proportion of KC dendritic input. We also note that KCs can modulate the output of modulatory neurons through axo-axonic connections (Eichler *et al*. 2017, Cervantes-Sandoval *et al*. 2017, Takemura *et al*. 2017), but the presynaptic modulatory input likely has a very distinct functional role from the dendritic input and we do not consider it further in this study. *Bottom:* Table shows percent of inputs onto dendrites of modulatory neurons from (I.) all known feedback types (direct MBON feedback, one-step FBN feedback, and two-step FB2N feedback), (II.) FFNs (likely conveying feedforward input from US pathways), and their ratio. Note that for most modulatory neurons this ratio is greater than 1 (color coded in blue), suggesting feedback input may be at least as important as feedforward input for regulating modulatory neuron activity. **e** Connectivity matrix showing normalized synaptic input (expressed as % input, computed as in **b**) each modulatory neuron (*columns*) receives from each pre-modulatory neuron (*rows*). Only reliable connections are shown for which both the left and right homologous connections have at least 3 synapses, and their sum is at least 10. Modulatory neurons are color-coded as in c according to the type of memory they can induce, *red* for aversive and *blue* for appetitive). Many pre-modulatory neurons (*rows*) synapse onto a single modulatory neuron, or onto modulatory neurons of similar function (*columns*). Functionally distinct modulatory neurons (*e.g*. DANs whose activation signals positive and negative valence, **Fig. 1b**) that innervate distinct MB lobes receive inputs from distinct subsets of pre-modulatory neurons (See also *Extended Data Fig. 3*). Interestingly, two UVL MBINs (OAN-e1 and MBIN-e2), whose activation paired with odor did not induce memory in our paradigm and that were not significantly activated by fictive punishments share a higher fraction of their input with the VL/LA DANs than with other modulatory neurons. This raises the possibility that the UVL modulatory neurons map be recruited by similar stimuli to the VL/LA DANs, but only in specific circumstances. **f** US pathways converge with feedback pathways from MBONs at multiple levels: at the modulatory neurons themselves, and at FB2Ns. The EM connectivity map shows the shortest identified pathways from distinct types of US pathways from somatosensory neurons to VL modulatory neurons. US pathways target both FFNs and FB2Ns. The hexagonal shape denotes a group of left-right homolog neurons. Connections with less than 10 synapses are not included (see *Supplemental Adjacency Matrix* for complete connections). *Orange*, Nociceptive MD IV specific pathway; *brown*, Mechanosensory Ch specific pathway; *blue*, multimodal Basins 2&4 pathway. Thickness of the arrow is proportional to normalized synaptic input that a postsynaptic neuron type receives from a presynaptic neuron type at the source of the arrow, defined as in **b**.

### Relationship between functional diversity and input diversity of MB modulatory neurons

We wondered how the functional diversity of modulatory neurons relates to their input diversity. As expected, functionally distinct DANs (whose activation leads to aversive or appetitive memory for paired odors; Fig. 1c) that innervate distinct lobes receive inputs from distinct subsets of pre-modulatory neurons (Fig. 2e and Extended Data Fig. 3). We found that even some functionally distinct modulatory neurons that innervate the same compartment receive inputs from drastically different subsets of pre-modulatory neurons: for example DAN-g1 and OAN-g1 that express different neuromodulators and differ in their ability to induce memory (*e.g*. Fig. 1c, 2e and Extended Data Fig. 3). Such difference in input structure suggests that some modulatory neurons that innervate the same compartment may be differentially recruited during learning.

In contrast, functionally similar VL/LA-DANs (whose activation leads to aversive memory for paired odors) share a higher fraction of presynaptic partners with each other than with other DANs (Fig. 2e and Extended Data Fig. 3). Nevertheless, even these functionally similar VL/LA DANs receive input from distinct combinations of input neurons (Fig. 2e), potentially explaining why they have similar, but not identical tuning properties to different punishment types (Fig. 1f).

In summary, we found that each modulatory neuron type that is distinguishable based on the compartment it innervates, or based on neurotransmitter expression, receives input from a unique combination of neurons and thus potentially encodes a unique set of features.

### Feedback neurons reveal a highly recurrent architecture for computing teaching signals

We aimed to characterize the pre-modulatory neurons based on the inputs they receive. Specifically, we asked whether they can convey feedback information about previously formed memories (via input originating from MBONs), or about received US (via afferent input from sensory neurons), or both. Surprisingly, we found that the majority (61/102) of pre-modulatory neurons receive feedback input from MBONs (Fig. 2a-c, Extended Data Fig. 4a-c). 40 left-right homologous neuron pairs receive reliable (as defined above) direct input from MBONs, providing one-step feedback from MBONs to modulatory neurons (we call these one-step feedback neurons, FBNs, Fig. 2a-c, Extended Data Fig. 4a). Another 21 pre-modulatory neuron pairs receive reliable direct input from FBNs (but not MBONs) and provide two-step feedback from MBONs to modulatory neurons (we call these two-step feedback neurons, FB2Ns, Fig. 2a-c, Extended Data Fig. 4b). The majority of FBNs also receive input from other FBNs, providing two-step, as well as one-step feedback (Fig. 2b-c and Extended Data Fig. 4a). The remaining (41/102) pre-modulatory input neuron pairs do not receive reliable direct MBON, FBN, nor FB2N input, so we classified them tentatively as “feedforward neurons” (FFNs, Fig. 2a-c).

To determine the likelihood that MBONs could influence modulatory neuron activity, via the feedback pathways we also analyzed the fraction of total input that FBNs and FB2Ns receive from MBONs, and that modulatory neurons receive from feedback pathways. In previous studies we have demonstrated functional connections even when neurons received 2% of their input from another neuron^62,124^. We found that individual FBNs receive on average 12% of their total synaptic input from MBONs and 26% from MBONs and FBNs combined (Extended Data Fig. 4a and c). Similarly, individual FB2Ns receive on average 17% of their total synaptic input from FBNs and 28% from FBNs and FB2Ns combined (Extended Data Fig. 4b and c). Based on these input fractions we expect that MBONs can significantly influence FBN and FB2N activity. Strikingly, we found that many modulatory neurons receive more than 50% of their total dendritic input from feedback pathways, including directly from MBONs, one-step, and two-step feedback (Fig. 2d). This suggests that modulatory neuron activity could be strongly modulated by MBON activity, via the newly discovered one-step and two-step feedback neurons.

### Multilevel convergence of afferent US pathways with feedback pathways

We investigated how the feedback pathways from MBONs converge with afferent pathways from US sensory neurons. We focused on the VL/LA-DANs that we identified as responding to nociceptive and/or mechanosensory neuron activation (Fig. 1f) and asked whether they receive the somatosensory and MBON input via distinct or overlapping pre-modulatory neurons. In the larva, the early portions of the somatosensory circuits that process aversive cues are well characterized^62,124,129,130^. We had previously reconstructed all 1st order PNs downstream of nociceptive and mechanosensory sensory neurons, a subset of 2nd order PNs, and a few 3rd order PNs^62,124^. This enabled us to search for shortest pathways from the nociceptive and mechanosensory sensory neurons to the VL/LA-modulatory neurons. We note that the pathways identified in this way represent only a subset of existing pathways, because not all of the 2nd, 3rd and 4th order somatosensory PNs have been reconstructed. Nevertheless, we were able to identify two-, three-, and four-step pathways from the nociceptive and mechanosensory sensory neurons to six different pre-modulatory neurons that target the VL/LA-modulatory neurons: three FFNs and three FB2Ns (Fig. 2f, Extended Data Fig. 5, Supplementary Adjacency Matrix, Supplementary Atlas). Thus, the afferent US pathways converge with feedback pathways from MBONs at multiple levels: both onto the modulatory neurons themselves (via FFNs) and onto the pre-modulatory FB2Ns.

### Modulatory neurons receive convergent one-step feedback from multiple MBONs from functionally distinct compartments

Next, we analyzed in more detail the types of one-step feedback motifs formed by FBNs (Fig. 2a-c, 3a-c, Extended Data Fig. 6a-b, Supplementary Adjacency Matrix, and Supplementary Atlas). Specifically, we asked whether FBNs mostly provide input to their own compartment, or whether they link multiple compartments for forming distinct types of memories. We observed a surprising diversity of one-step FBNs that linked unique combinations of MBONs with unique combinations of modulatory neurons (Fig. 3a and Extended Data Fig. 6a-b). Some (7/40) FBNs provide exclusively within-compartment feedback (Fig. 3a). Some (13/40) provide exclusively cross-compartment feedback (we named individual neurons of this subset of FBNs, FANs, for feed-across neurons, Fig. 3a). Some (8/40) FBNs synapse onto multiple modulatory neurons from multiple compartments (Fig. 3a and Extended Data Fig. 6b). Interestingly, the largest class of FBNs (17/40, Fig. 3a, 3d, and Extended Data Fig. 6a) receives input from multiple MBONs and appears to be well suited for comparing odor drive to functionally distinct compartments of the MB (Fig. 1c). Thus, almost all of these FBNs (at least 13/17, and potentially more, but the neurotransmitters of all MBONs are not known) receive GABAergic (inhibitory) or glutamatergic (potentially also inhibitory^131,132^ in insects) input from MBONs from compartments implicated in memory formation, and cholinergic (excitatory) inputs from MBONs from other compartments (Fig. 3a, 3d, and Extended Data Fig. 6a). The comparison of inhibitory and excitatory input may enable these FBNs to more accurately read out the results of learning-induced plasticity^55,85,133^ in memory compartments, relative to other compartments.

**Figure 3.**
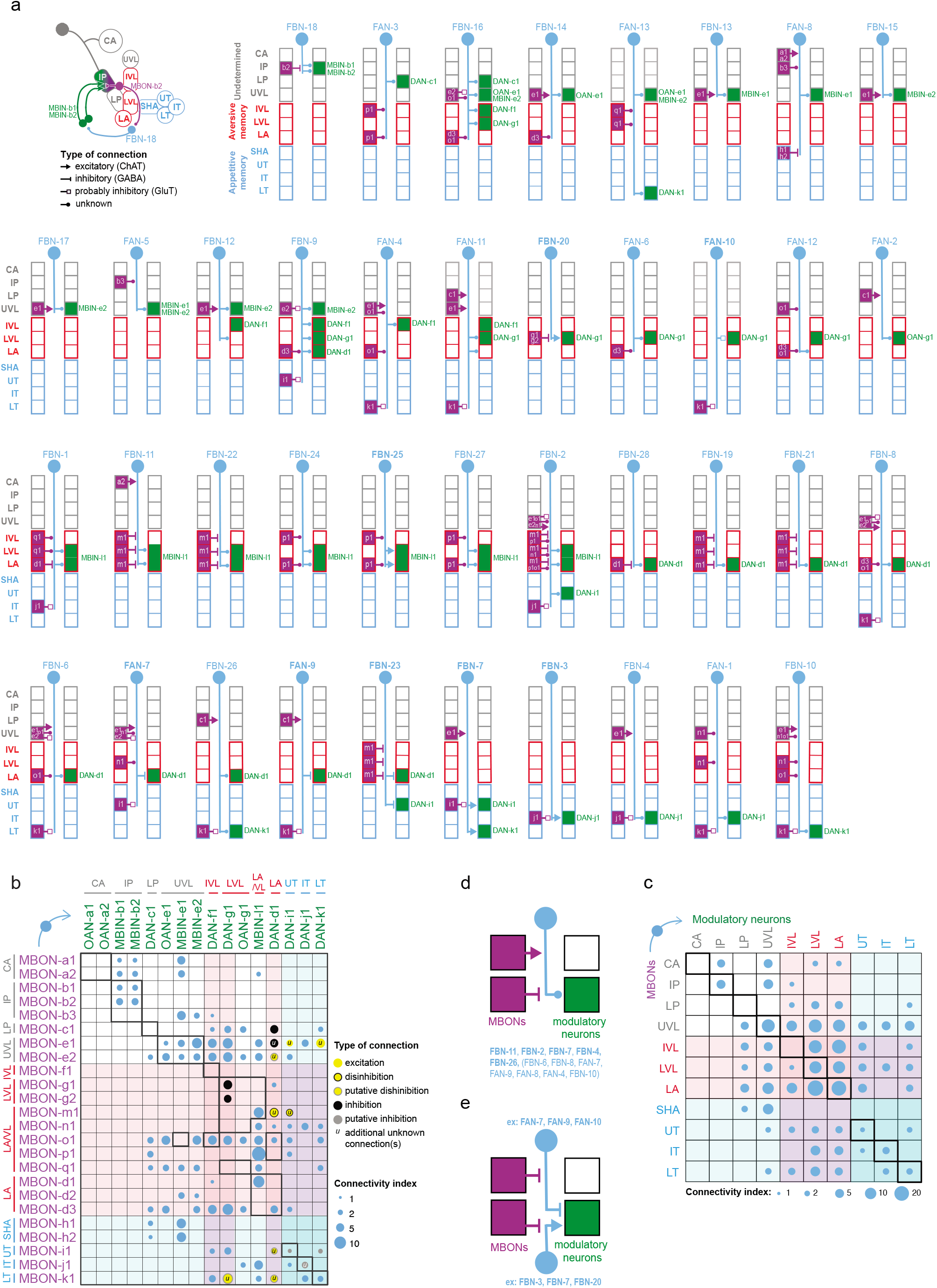
Modulatory neurons receive convergent one-step feedback from multiple MBONs from functionally distinct compartments. **a** Connectivity of each of the 40 feedback neuron (FBN) pairs that provide one-step feedback from MBONs to DANs. Each diagram represents the connectivity of a single left-right pair of homologous FBNs. Each box indicates a separate compartment. *Purple*, compartment(s) of the presynaptic MBON(s). *Green*, compartment(s) of the postsynaptic modulatory neuron(s). FBNs are ordered according to the modulatory neuron they innervate, starting with peduncle modulatory neurons and ending with the medial lobe ones. Classical neurotransmitter profiles of the MBONs and FBNs are indicated by the arrow (cholinergic, excitatory connection), vertical line (GABAergic, inhibitory connection) or square (glutamatergic, probably also inhibitory connection) for the neurons for which they are known from immunostaining (For images, see *Extended Data Fig. 10* for FBNs and Eichler *et al*. 2017 *Extended Data Fig. 2b* for MBONs), or by a circle when they are unknown. 7 FBNs provide exclusively within-compartment feedback. 13 FBNs provide exclusively cross-compartment feedback (named FANs, for feed-across). 8 FBNs synapse onto multiple modulatory neurons from multiple compartments. The largest class of FBNs (17) receives input from multiple MBONs, with the majority (at least 13 and maybe more) receiving input of potentially opposite sign from MBONs from functionally distinct compartments. **b** Modulatory neurons receive input from multiple MBONs from functionally distinct compartments via the FBNs. Connectivity matrix shows connections between MBONs and modulatory neurons via the indirect one-step feedback pathways obtained by multiplying the MBON→FBN, and FBN→modulatory neuron connectivity matrices (normalized as in **Fig. 2b** and including all connections where the presynaptic neuron accounts for at least 1% of input onto the postsynaptic neuron). A connectivity index was computed by taking the square root of the numbers in the resulting matrix product. A connectivity index of 1, 10, and 100 means that for both connections comprising that indirect feedback pathway the presynaptic neuron accounts for 1%, 10%, and 100% of input onto that postsynaptic neuron, respectively. When the neurotransmitters of both the MBON and the FBN(s) that comprise a connection are known, the circle is color-coded to represent types of connection: excitatory (ChAT and ChAT), disinhibitory (GABA and GABA), probably disinhibitory (GluT and GABA/GluT), inhibitory (GABA and ChAT), probably inhibitory (GluT and ChAT). Color shades represent the valence of the memory formed in a given compartment (*red*: aversive memory, *blue*: appetitive memory). True within-compartment feedback connections from an MBON that receives direct synaptic input from that modulatory neuron are boxed in bold (*e.g*. from MBON-g1 and MBON-g2 onto DAN-g1). Some multicompartment MBONs provide feedback to modulatory neurons from the same compartment that do not synapse onto them directly, and we do not consider these to be true within-compartment feedback connections (*e.g* from MBON-m1 onto DAN-d1). Note that all four true within-compartment feedback connections with known neurotransmitters are potentially inhibitory: MBON-g1 to DAN-g1, MBON-g2 to DAN g2, MBON-i1 to DAN-i1 and MBON-j1 to DAN-j1. In contrast, many cross-compartment connections are potentially excitatory or disinhibitory. **c** All compartments except CA and IP receive one-step feedback from multiple compartments and from each of the three functionally distinct regions of the MB. Matrix shows the relative one-step connection strength index from a row compartment to a column compartment via FBNs. Strength is calculated as in b, but pooled (summed and normalized) for all MBONs from a compartment and all modulatory neurons from a compartment, and then multiplied by 100. Note the compartments of the VL (vertical lobe: UVL, IVL, LVL) and LA (lateral appendix) are strongly interconnected. The color shades are as in **b**. **d-e** Summary diagram of commonly observed convergence motifs. Feedback connections of opposite sign from functionally distinct compartments converge onto FBNs and DANs. In each diagram, the FBN (*blue*) receives direct input from one or more MBON(s) (*purple*), and connects with the postsynaptic modulatory neuron(s) (*green*). Each box denotes a separate compartment. The type of connection (GABAergic, glutamatergic, cholinergic, or unknown) is represented by different arrowheads as described in the legend in **a. d** More than a quarter of FBNs (at least 12, and potentially more) receive direct GABAergic (inhibitory) or glutamatergic (also potentially inhibitory) input from MBONs from one compartment and direct cholinergic (excitatory) input from MBONs from a functionally distinct compartments enabling them to compare the odor drive to these MBONs. **e** Many DANs (DAN-f1, d1, i1, j1, and k1) receive potentially inhibitory (excitatory FBN downstream of an inhibitory MBON) one-step feedback from MBONs from one compartment and potentially disinhibitory (inhibitory FBN downstream of an inhibitory MBON) or excitatory (excitatory FBN downstream of an excitatory MBON) one-step feedback from MBONs from a functionally distinct compartment. A common pattern for the lobe DANs implicated in memory formation may be a likely inhibitory connection from an MBON from their own compartment and a likely disinhibitory connection from an MBON from a compartment of opposite valence (observed for both DAN-g1 and i1), that could enable these DANs to compare the odor drive to MBONs from compartments of opposite valence.

We also found that most modulatory neurons receive input from multiple FBNs. For example, all ML-DANs and VL-DANs capable of evoking olfactory memory (Fig. 1c) received input from at least three different FBNs (Fig. 2e, 3a, and Extended Data Fig. 6b). Clustering FBNs based on the pattern of output to modulatory neurons revealed that some UVL, LVL and LA modulatory neurons stand out as prominent targets of feedback input, receiving significant input from clusters of 7 or more different FBNs (*e.g*. neurons forming aversive memory DAN-d1, DAN-g1, and neurons of unknown function MBIN-l1, MBIN-e1 and -e2, Extended Data Fig. 6b and 7). Similarly, clustering FBNs based on the input from MBONs revealed that specific VL-MBONs stood out as prominent sources that provide output to clusters of FBNs (for example, the cholinergic UVL MBON-e1 strongly targets a cluster of 13 different FBNs, Extended Data Fig. 6a and 8).

Since most FBNs receive input from multiple MBONs, and most modulatory neurons receive input from multiple FBNs, we analyzed the connections from all MBONs to all modulatory neurons via all possible one-step feedback pathways (by multiplying normalized MBON-to-FBN and FBN-to-modulatory neuron connectivity matrices from Supplementary Adjacency Matrix). We found that most modulatory neurons received indirect one-step feedback from many MBONs (Fig. 3b). Most compartments therefore received one-step feedback from many other compartments (Fig. 3c). All modulatory neurons except those that innervate CA and IP received one-step feedback from each of the three functionally distinct regions of the MB: UVL (unknown function), VL aversive memory compartments, and ML appetitive memory compartments (Fig. 3b-c). This is in stark contrast to the direct connectivity from MBONs to modulatory neurons, which is very sparse and connects very few compartments (Extended data fig. 9a-b). Thus, the newly discovered FBNs greatly increase the connectivity between MBONs and modulatory neurons, enabling the output from multiple functionally distinct regions of the MB to influence the activity of a single modulatory neuron during memory formation.

### A modulatory neuron receives inhibitory and excitatory feedback from compartments of opposite valence

To gain a better understanding of how feedback motifs could influence modulatory neuron activity we wanted to determine which feedback neurons were excitatory and which inhibitory. We were able to identify GAL4 lines^134^ that drive expression in eight different FBNs and three FB2Ns. We could therefore label these feedback neurons with GFP^135^ and test whether they express GABA^136^, vesicular glutamate transporter (vGlut)^137^, or choline acetyl transferase (ChAT)^138^ using immunohistochemistry. Unfortunately, while these GAL4 lines were selective enough to allow visualization of the relevant neurons, most lines were not selective enough to enable targeted manipulation for functional connectivity and behavioral analysis. We found that four of the tested FBNs were cholinergic (*i.e*. excitatory), three were GABAergic (*i.e*. inhibitory), and one was glutamatergic (possibly also inhibitory^131,132^, Fig. 3a and Extended Data Fig. 10). Two FB2Ns were glutamatergic and one was cholinergic (Extended Data Fig. 10).

For a few cases where we could identify the neurotransmitter profiles of both the MBON^47^ and the FBN in a one-step feedback connection, we attempted to predict the signs of these connections (Fig. 3b). All of the true within-compartment feedback connections with known neurotransmitters were potentially inhibitory (4/4), comprising a GABAergic or glutamatergic MBON and an excitatory FBN (Fig. 3b and 3e, MBON-g1 to DAN-g1, MBON-g2 to DAN-g1, MBON-i1 to DAN-i1 and MBON-j1 to DAN-j1). In contrast, most of the (8/11) cross-compartment connections with known neurotransmitters were potentially functionally excitatory, either disinhibitory (comprising an inhibitory MBON and an inhibitory FBN), or excitatory (comprising an excitatory MBON and an excitatory FBN, Fig. 3b and 3e). Out of those, all of the connections between compartments of opposite valence (4/4) were potentially disinhibitory (Fig. 3b and 3e). Furthermore, we observed that some modulatory neurons (*e.g*. DAN-g1 and DAN-i1) received both potentially inhibitory feedback from their own compartment and potentially excitatory feedback from compartments of opposite valence (Fig. 3b and 3e).

We designed experiments to functionally confirm the two types of predicted feedback connections onto the same DAN (Fig. 3e, 4a-g). We were able to identify a strong LexA line for DAN-i1^78^. DAN-i1 receives potentially inhibitory one-step feedback from the glutamatergic MBON-i1 in its own compartment (via the excitatory FBN-7, Fig. 4a-b), and potentially disinhibitory one-step feedback from the GABAergic MBON-m1 from compartments of opposite valence (via the GABAergic FBN-23, Fig. 4a and 4e). Neither of these MBONs synapses directly onto DAN-i1. We also generated Split-GAL4 lines to selectively express Chrimson in MBON-i1 or in MBON–m1. We activated these MBONs optogenetically while recording intracellularly from DAN-i1 (labelled with GFP using the LexA line).

**Figure 4.**
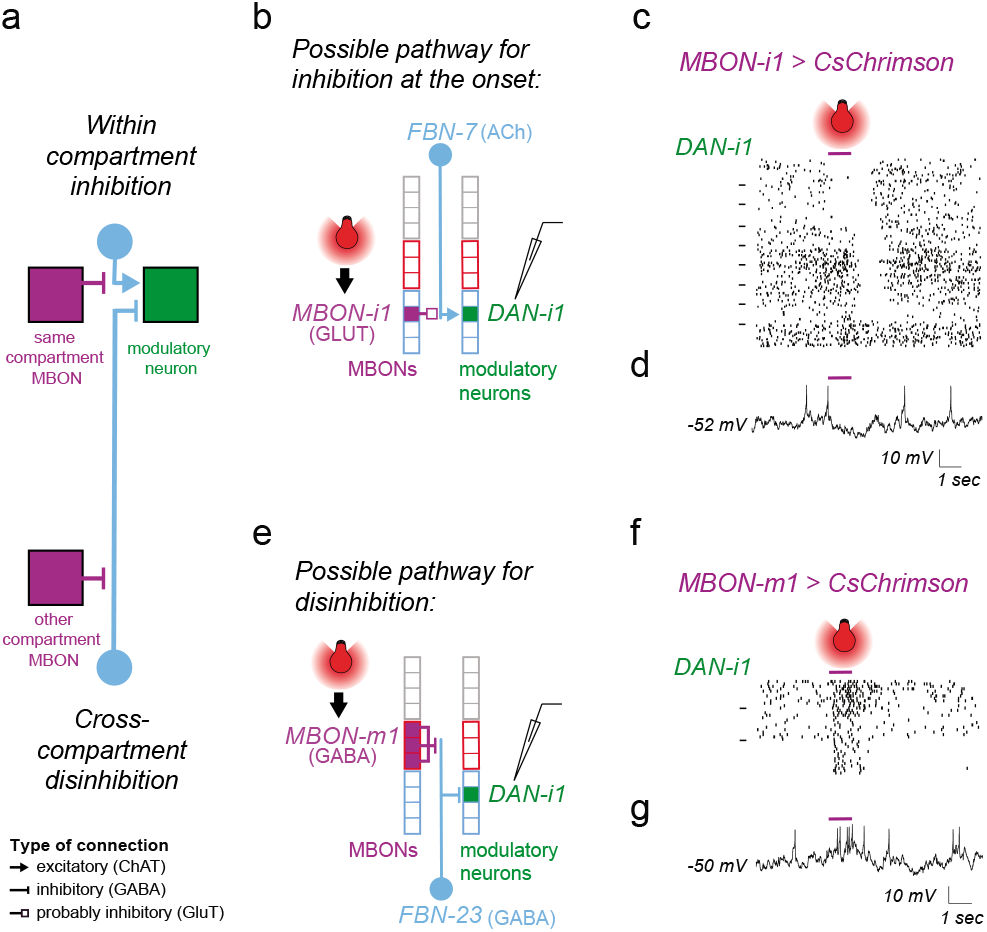
Functional inhibitory and excitatory feedback connections from compartments of opposite valence converge onto a DAN. **a** Schematic diagram showing indirect inhibitory within-compartment feedback and disinhibitory feedback from a compartment of opposite valence converging onto the same DAN, as predicted based on connectivity and neurotransmitter profiles. These two types of predicted connections are tested separately using optogenetic activation of the MBONs and patch-clamp recording of a DAN (**b-d** and **e-g**). *Boxes* denote compartments. *Blue* and *red* outlines, appetitive and aversive memory compartments, respectively. *Purple*, *light blue*, and *green*, MBONs, FBNs, and DANs, respectively. **b** The cholinergic FBN-7 downstream of the glutamatergic medial lobe MBON-i1 could mediate inhibitory one-step within-compartment feedback onto DAN-i1. **c** Whole-cell patch-clamp recording of DAN-i1 (medial lobe) during optogenetic activation of the medial lobe MBON-i1 of the same compartment reveals inhibitory feedback at the onset or offset of MBON-i1 activation. The action potentials of all 180 electrophysiologically recorded traces from 9 animals in response to MBON-i1 activation (*purple bar*) are shown as raster plots. Dashes on the left separate rasters belonging to distinct animals. We observed two types of inhibitory responses in DAN-i1 to MBON-i1 activation: a long-latency inhibitory response to the onset of MBON-i1 activation in 3/9 animals (shown at the top, 55.3 ± 17.3 ms, n=60 traces from 3 animals), and an even longer latency inhibitory response to the offset of MBON-i1 activation in 4/9 animals (shown in the middle, 95.3 ± 43.5 ms, n=80 trials from 4 animals). In two animals (shown at the bottom) DAN-i1 showed no response to MBON-i1 activation (n=40 trials). Inhibition at the onset in some animals but not others might result from distinct baseline states of FBN, as shown in the model in *Extended Data Fig. 11b*. Inhibition at the offset might be result of post-inhibitory rebound within a two-step feedback pathway as shown in *Extended Data Fig. 11c*. **d** An example individual trace from c. More examples are shown in *Extended Data Fig. 11a*. **e** The GABAergic FBN-23 downstream of the GABAergic vertical lobe MBON-m1 could mediate a disinhibitory one-step connection onto the medial lobe DAN-i1. The vertical lobe MBON-m1 receives input from DAN-g1, whose activation can induce aversive memory. In contrast, the medial lobe DAN-i1 activation induces appetitive memory. **f** Whole-cell patch-clamp recording of DAN-i1 during optogenetic activation of the medial lobe MBON-m1 from a compartment of opposite valence reveals an excitatory cross-compartment feedback connection. The action potentials of all the 45 electrophysiologically recorded traces from the 3 animals in response to MBON-m1 activation (*purple bar*) are shown as raster plots. In response to optogenetic activation of MBON-m1, we observed a long-latency excitatory response in DAN-i1 in 3/3 animals tested (51.3 ± 7.7 ms, n=45 trials from 3 animals). **g** An example individual trace from **f** (more examples in *Extended Data Fig. 11d*).

Activating the glutamatergic MBON-i1 evoked long-latency (55ms +/− 17) inhibitory responses in DAN-i1 on 9/27 trials (on all 3 trials in 3/9 animals, Fig. 4c-d, Extended Data Fig. 11a). The inter-animal variability could potentially be due to different baseline activity levels of FBNs mediating this connection: activating an inhibitory MBON can only lead to detectable inhibition of the DAN if the excitatory FBN targeted by the MBON has baseline activity, as illustrated by the simple rate-model in Extended data Fig. 11b. On 12/27 trials (on all 3 trials in 4/9 animals, Fig. 4c and Extended Data Fig. 11a) we observed very long latency inhibitory responses to the offset of MBON-i1 activation only. These inhibitory responses to the offset had a longer latency (95 ms +/− 44) than the inhibitory responses to the onset (55 ms +/− 17) of MBON-i1 activation and could therefore be mediated by a longer two-step feedback pathway (as proposed in the Extended Data Fig. 11c).

In contrast, we found that activating the GABAergic MBON-m1 evoked excitatory responses in DAN-i1 on 9/9 trials (on all 3 trials in 3/3 animals) with a similar latency (47 ms +/− 9) to the inhibitory responses evoked by MBON-i1 activation (Fig. 4e-g and Extended Data Fig. 11d). In summary, we confirmed with physiological recording an inference we had made from structural connectivity and neurotransmitter information: that functionally inhibitory and excitatory MBON connections from compartments of opposite valence converge onto the same DAN (Fig. 4a-g). DANs that receive this pattern of feedback could compare the odor-evoked excitation of MBONs in compartments of opposite valence and thereby compute the integrated predicted value of an odor across aversive and appetitive memory systems.

### Two-step feedback from most MBONs to most modulatory neurons further increases intercompartment connectivity

Next, we investigated in more detail the two-step feedback motifs (Fig. 2a-c, 5a-e, Extended Data Extended Data Fig. 12, 13a-d, and 14, Supplementary Adjacency Matrix and Supplementary Atlas). We found 21 FB2N pairs, which do not themselves receive direct MBON input, but do receive input from FBNs, thus providing two-step feedback from MBONs to modulatory neurons (Fig. 2a-c, 5a-b, Extended Data Fig. 4b-c, Extended Data Fig 12 and 13b-c, Supplementary Adjacency Matrix and Supplementary Atlas). Many FBNs also receive input from other FBNs (Fig. 5c-d, Extended Data Fig. 4a, 4c, 13a and 14) and provide both two-step and one-step feedback. We therefore also analyzed the connections between all MBONs to all modulatory neurons via the two-step pathways (by multiplying the MBON-FBN, FBN-FB2N/FBN and FB2N/FBN-modulatory neuron normalized connectivity matrices from Supplementary Adjacency Matrix). We found two-step feedback from most MBONs to most modulatory neurons further increases inter-compartment connectivity (Fig. 5e and Extended Data Fig. 9b). We were able to determine neurotransmitter profiles for seven neurons that provide two-step feedback: three (FBN-25, FBN-7, and FB2N-19) were cholinergic, two (FBN-23 and FAN-7) were GABAergic, and two (FAN-10 and FB2N-18) were glutamatergic (Fig. 5b and d, Extended Data Fig. 10). In summary we found a diverse set of two-step feedback motifs that could support within- and crosscompartment computations.

**Figure 5.**
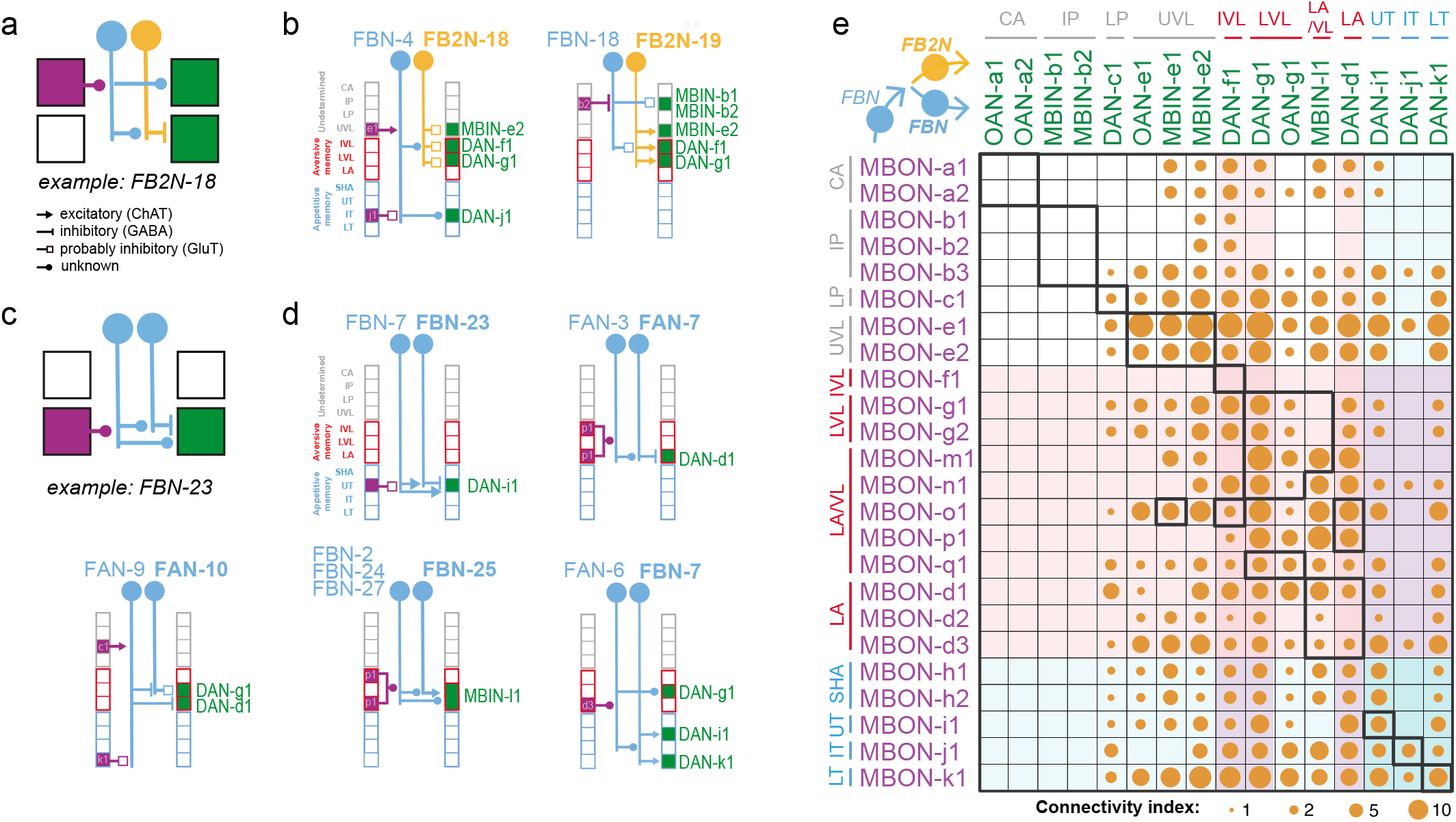
Two-step feedback from most MBONs to most modulatory neurons further increases inter-compartment connectivity. **a** Schematic diagram of a two-step feedback motif involving an FBN (*blue*) and an inhibitory FB2N (*yellow*). The FBN provides one-step feedback to some compartments and two-step feedback to others via the FB2N. **b** Two example two-step within-compartment feedback motifs involving FB2Ns with identified neurotransmitters. In the first example, FBN-4 integrates opposite drives from MBON-e1 and MBON-j1 and transmits its signal directly to ML DAN-j1 and indirectly to VL modulatory neurons via the glutamatergic FB2N-18. In the second example, FBN-18 distributes a disinhibitory signal from MBON-b2 to many modulatory neurons both via one-step feedback and two-step feedback involving the cholinergic FB2N-19. Arrowheads denote the type of synaptic connection as in **a**. **c** Schematic diagram of a two-step feedback motif involving two FBNs (*blue*) rather than an FBN and an FB2N. The FBN provides one-step feedback to some compartments and two-step feedback to others via another FBN. **d** Five example two-step within-compartment feedback motifs involving FBNs with identified neurotransmitters. In these examples, FBN-23, FAN-7 or FBN-25 provide within-compartment two-step feedbacks onto DAN-i1, DAN-d1 and MBIN-l1 respectively. FAN-10 and FBN-7 distribute the signal of their presynaptic FBN to modulatory neurons in other compartments. The type of synaptic connection is symbolized by different arrowheads as in **a**. **e** Most modulatory neurons receive two-step feedback from most MBONs via the FBNs and FB2Ns. Connectivity matrix shows connections between MBONs and modulatory neurons via two-step feedback pathways, obtained by multiplying the MBON→FBN, FBN→FB2N/FBN and FB2N/FBN→modulatory neuron connectivity matrices (normalized as in **Fig. 2b** and including all connections where the presynaptic neuron accounts for at least 1% of input onto the postsynaptic neuron). The connectivity index was computed by taking the cubic root of the numbers in the resulting matrix product. A connectivity index of 1, 10, and 100 means that for the three connections comprising that indirect feedback pathway the presynaptic neuron accounts for 1%, 10%, and 100% of input onto that postsynaptic neuron, respectively.

### Feedback neurons can drive memory formation

So far, we have shown that at least some of the indirect feedback connections from MBONs to DANs are functional (Fig. 4b-g). However, we also wanted to test whether the feedback neurons can sufficiently influence DAN activity to actually induce learning. We succeeded in generating Split-GAL4 lines^102^ that drive expression in one or very few neuron types for: a cholinergic FB2N, a glutamatergic FB2N, and a GABAergic FBN that project onto DANs whose activation can induce aversive memory (Fig. 6a-c, Extended Data Fig. 1). We asked whether optogenetic activation of these feedback neurons (without directly activating any modulatory neurons) was sufficient to induce memory in our olfactory training paradigm (Fig. 6d-f). We found that pairing of an odor with activation of the excitatory cholinergic FB2N-19 induces aversive memory (Fig. 6d and Extended Data Fig. 2a-b), similar to direct activation of its postsynaptic DAN-f1 (Fig. 1c). In contrast, pairing of an odor with the activation of the GABAergic FAN-7 induces appetitive memory (Fig. 6e and Extended Data Fig. 2a-b), opposite to direct activation of its postsynaptic DAN-d1 (Fig. 1c). The line that drives expression in FAN-7 also drives expression in another neuron pair in the brain (MB2ON-86, Fig. 6c), but pairing an odor with the activation of that neuron alone did not induce memory (Fig. 6e and Extended Data Fig. 2a-b). The line also drives expression in a few somatosensory interneurons in the nerve cord, but somatosensory pathways are expected to induce aversive memory (Fig. 1e). Similarly, pairing of an odor with the activation of the glutamatergic FB2N-18 and FB2N-11 (likely also inhibitory) induces appetitive memory (Fig. 6f, and Extended Data Fig. 2a-b), opposite to direct activation of their postsynaptic DAN-f1 and DAN-g1 (Fig. 1c). Thus, at least some feedback neurons can induce memory formation. Interestingly, activation of inhibitory feedback neurons induces memories of opposite valence to the activation of the DANs that they inhibit.

**Figure 6.**
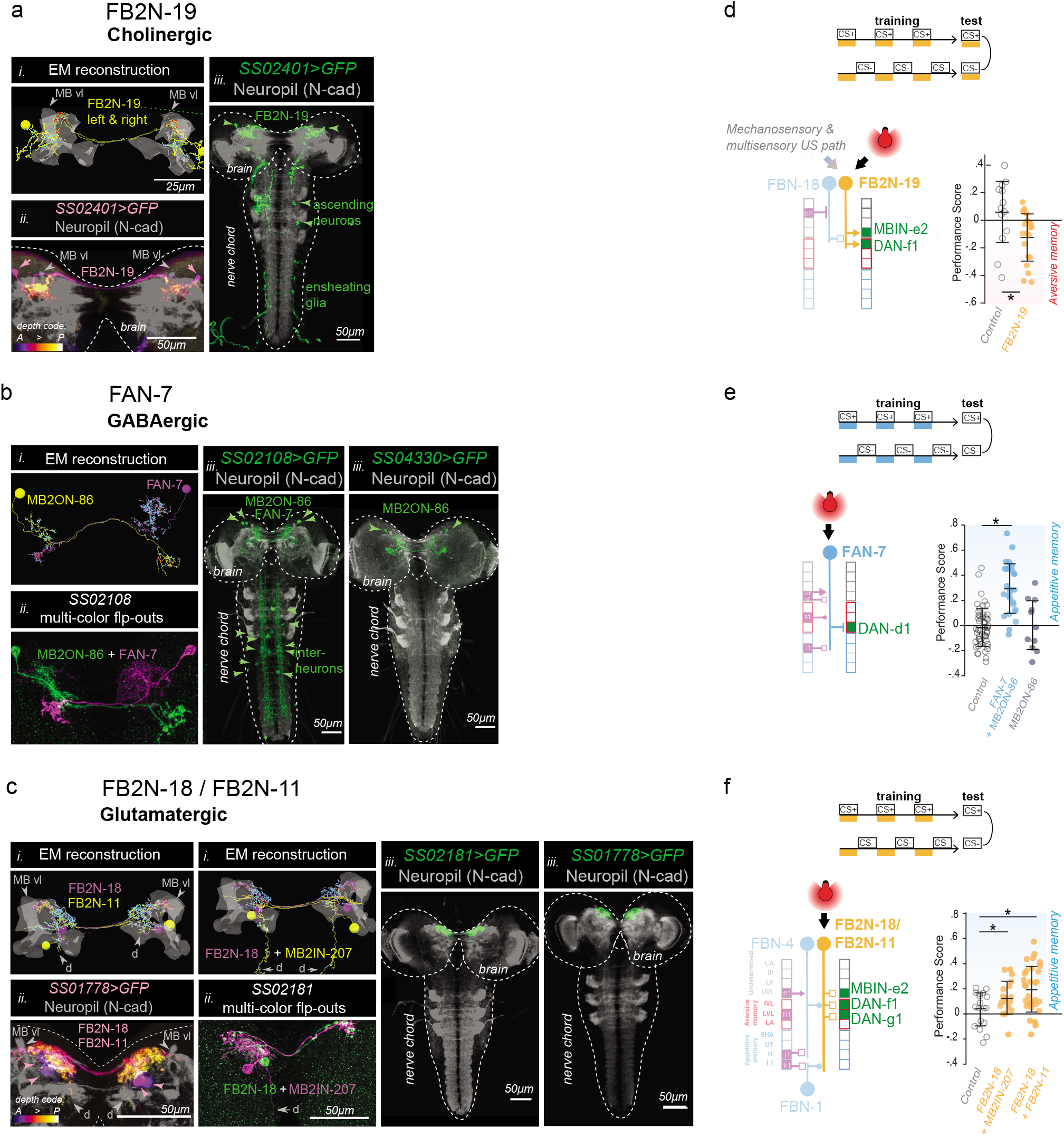
Feedback neurons can drive associative memory formation. We were able to generate Split-GAL4 lines that drive expression in a single pair of neurons, or in very few cell types, for three different pairs of FBNs or FB2Ns that target VL DANs (**a-c**). We used these lines to optogenetically activate these neurons instead of a US during an associative learning paradigm (**d-f**). **a-c** Identification of driver lines for EM-reconstructed neurons. *i*) Skeletons of specific feedback neurons reconstructed in the EM. *Red dots*, presynaptic sites. *Blue dots*, postsynaptic sites. *Grey*, mushroom body vertical lobe (*MB vl*) for reference. d, dendritic arbor. *ii*) Maximum intensity projections of confocal stacks of larval brains showing the same neurons visualized with reporters targeted using specific Split-GAL4 lines. For some lines multicolor FLP-outs were used to visualize each neuron in a different color to facilitate identification and comparison with EM. Grey, neuropil visualized with N-cad. *Dashed line*, brain outline. *iii*) Maximum intensity projections of confocal stacks of the entire nervous system showing the complete expression pattern of each line revealed by driving UAS-myr-GFP. Grey, neuropil visualized with N-cad. *Dashed line*, nervous system outline. **a**, The *SS02401-Split-GAL4* line drives expression in FB2N-19 (*i*) in the brain (*ii*), and very weakly and stochastically (not reproducibly in all samples) in a few ascending neurons and ensheathing glia in the nerve chord that are unlikely to have an ability to evoke associative memory, due to weak and stochastic expression (*iii*). **b**, The *SS02108-Split-GAL4* line drives expression in FAN-7 and MB2ON-86 (*i*) in the brain visualized with multicolor flp-outs in (*ii*). Complete expression pattern of *SS02108-Split-GAL4* visualized with *UAS-myr-GFP* shows additional expression in a few somatosensory interneurons in the nerve cord, called ladders, that mediate avoidance behavior and are hence unlikely to have a positive valence and evoke an appetitive memory. We identified the *SS04330-Split-GAL4* line as driving expression specifically in the MB2ON-86 neuron and used it as an additional control in **e**. **c**, The *SS01778-Split-GAL4* line drives expression in both FB2N-18 and FB2N-11, which have very similar morphology and very similar connectivity (Extended Data Fig. 12 and 13b-d). They both connect strongly to DAN-f1 and weakly (but reliably) to MBIN-e2; FB2N-18 also connects weakly (but reliably) to DAN-g1 (Extended Data Fig. 13c). The *SS02181-Split-GAL4* line (ii shows multi-color flpouts) drives expression in FB2N-18 and in MB2IN-207, one of the weakly connected pre-modulatory neurons from lineage DAMv12. Notice the ventrally projecting dendrite (*d*), a distinctive feature of MB2IN-207 neuron (*i*). *UAS-myr-GFP* expression patterns of the two lines show that they do not drive expression in any other neurons in the nerve cord (*iii*). **d-f** Pairing the optogenetic activation of the neurons in these lines with the odor ethyl acetate as a CS (as in **Fig. 1b-c**) induces associative memory. Plotted are the learning performance scores (computed as described in **Fig. 1b**) obtained after paired or unpaired optogenetic activation of these neurons with odor, compared to a corresponding empty line *w;attp40;attP2>UAS-CsChrimson (open circle)* as a control. Horizontal lines indicate means and standard deviations of the individual data points; *P < 0.05 Mann-Whitney U-test comparison between groups. **d**, Optogenetic activation of the excitatory cholinergic FB2N-19 that is presynaptic to DAN-f1 and MBIN-e2 (with *SS02401-Split-GAL4* line) induces aversive memory (*yellow*), same as the activation of its presynaptic DAN-f1 (**Fig. 1c**). Note that FB2N-19 has to be activated during the test for memory expression as is the case for many punishing stimuli for *Drosophila* larva (Gerber et al., 2009). **e**, Optogenetic activation of the inhibitory GABAergic FAN-7 (*SS02108-Split-GAL4* line) induces appetitive memory (*dark blue*), opposite to the aversive memory induced by the activation of its presynaptic DAN-d1(**Fig. 1c**). *SS02108-Split-GAL4* line drives expression both in FAN-7, as well as in MB2ON-86, but pairing the activation of MB2ON-86 alone with odor (to disambiguate) (using the *SS04330-Split-GAL4* line) did not evoke any memory (*dark grey*). **f**, Optogenetic activation of the glutamatergic FB2N-18 and FB2N-11 together (with the *SS01778-Split-GAL4* line) induces appetitive memory (*yellow*), opposite to activation of their presynaptic DANs-f1 and -g1 (**Fig. 1c**). Even activation of the glutamatergic FB2N-18 without FB2N-11, but with another weakly connected pre-modulatory neuron which is unlikely to be able to significantly influence modulatory neuron activity (with the *SS02181-Split-GAL4* line) induces appetitive memory.

### Connectivity-constrained model of the circuit reveals feedback neurons improve performance on complex learning tasks

To explore the computational consequences of the feedback neurons, we developed a model of the circuit constrained by i) the connectome, ii) the known neurotransmitter identities of MBONs^47^ and pre-modulatory neurons (Extended Data Fig. 10), and iii) the valences of compartments whose modulatory neuron activation evokes aversive or appetitive memory when paired with odor (Fig. 1c). We modeled modifications of KC to MBON connections using a synaptic plasticity rule that depends on the timing of KC and modulatory neuron activity, consistent with findings in larval^58^ and adult *Drosophila*^84,133,139^. We optimized the model using gradient descent to perform various associative learning tasks^140^ (see Materials and Methods) and assessed the contributions of different feedback types by repeating the optimization procedure for networks lacking such feedback and comparing their performance. Tasks included first-order conditioning, second-order conditioning, extinction, and context-dependent conditioning (Fig. 7a-b). First-order conditioning and extinction have been demonstrated in adult^5,13,14,53^ and larval *Drosophila*^141^. While second-order^7,20^ and context-dependent^22,142,143^ conditioning have so far been investigated only in adult insects, we used these as example tasks to probe the ability of the circuit to support conditioning paradigms requiring additional computations. In the case of second-order conditioning, a reinforcement predicting conditioned stimulus is used to reinforce a second stimulus, while in context-dependent conditioning, the US valence depends on a previous contextual input.

**Figure 7.**
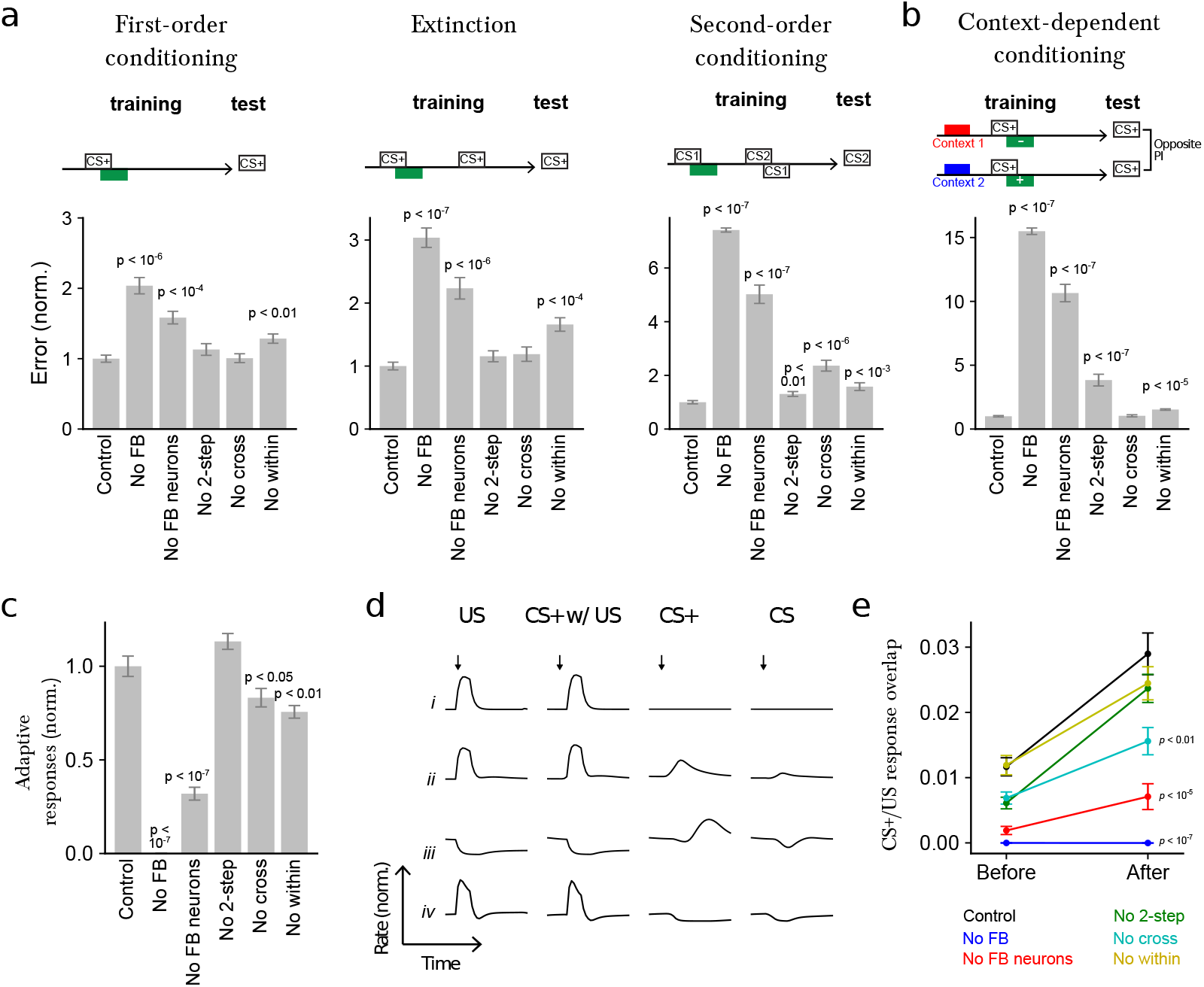
Model of the circuit reveals how each type of feedback connection improves performance on distinct learning tasks. **a** Normalized error (mean-squared difference between actual and and target PI, normalized to the error for control networks) after optimizing models to perform first-order conditioning, extinction, and second-order conditioning. Error is shown for six cases: Control: full network, No FB: networks in which all feedback onto modulatory neurons, including direct MBON connections, is removed, No FB neurons: networks in which indirect FBN/FB2N feedback is removed but direct MBON connections are intact, No 2-step: networks in which only FB2Ns and all FBN-to-FBN connections are removed, No cross: networks in which indirect cross-compartment connections are removed, No within: networks in which indirect within-compartment connections are removed. **b** Similar to **a**, but for networks optimized only to perform a context-dependent conditioning task. **c** Adaptive response index for the networks in **a**, defined as the magnitude of the change in firing rate responses to CS+ presentation before and after conditioning, averaged over modulatory neurons. The results are normalized by the value for control networks. **d** Selected example responses of DANs from networks in a to US alone (US), CS+ paired with US following training (CS+ w/ US), to CS+ alone after training (CS+), and to CS prior to training (CS). *Row i:* A DAN selective only to US that does not show adaptive responses to CS+. *Row ii:* A DAN selective to US that acquires a CS+ response after conditioning. *Row iii and iv:* DANs with “prediction-error” like responses. CS+ responses are opposite in sign to US responses. *Row iii:* a neuron that is inhibited by the US is activated when that US is omitted (*e.g*. when the CS+ is presented alone after training). *Row iv:* a neuron that is excited by the US is inhibited when that US is omitted (*e.g*. when the CS+ is presented alone after training). Note that the negative CS+ response is prolonged compared to the response to CS prior to training. **e** CS+/US response overlap before and after conditioning. The overlap is computed using the dot product of the vectors of firing rate changes across the modulatory neuron population during CS+ and US presentations. *p*-values represent a comparison to control networks, after conditioning.

We found that the performance on all tasks was significantly degraded in the absence of all feedback, including direct MBON feedback, one-step feedback via FBNs, and two-step feedback via FB2Ns and FBNs (Fig. 7a-b). The removal of the indirect feedback alone (with intact direct MBON feedback) also significantly degraded the performance on all tasks, with especially strong effects on the more complex tasks (Fig. 7a-b). Even the removal of two-step feedback alone significantly diminished performance on two of the more complex tasks (second-order conditioning and context-dependent conditioning), with a drastic effect on context-dependent conditioning (Fig. 7a-b). Thus, each additional feedback layer improves the performance of the network when it is tested on challenging associative learning tasks.

We also constructed networks lacking one- and two-step feedback only within or only across compartments. Removal of within-compartment feedback diminished performance on all tasks, while removal of cross-compartment communication substantially reduced performance for second-order conditioning (Fig. 7a-b). In total, each of the feedback pathways identified by the reconstruction may be important for associative learning paradigms that require computations such as prediction, prediction error, or context dependence.

### Feedback neurons enable adaptive responses of modulatory neurons in the model

The high fraction of feedback input originating from MBONs onto modulatory neurons suggests that their activity could be adaptively regulated by prior learning. To test this idea, we computed an index that quantifies the mean change in modulatory neuron firing rates in response to CS+ (*i.e*. the CS that was paired with the US) presentations before and after conditioning in the model and found that it is indeed substantially enhanced by the presence of feedback neurons (Fig. 7c). The optimized networks exhibit a diversity of adaptive modulatory neuron responses (some examples are shown in Fig. 7d).

After a CS/US pairing, many modulatory neurons acquired responses to CS+ that resemble their responses to the US that had been paired with that CS+ (Fig. 7e and 7d-ii). These responses were significantly attenuated in networks that lacked feedback, including those that lacked just indirect feedback, and just cross-compartment feedback (Fig. 7e). Such responses have been observed in modulatory neurons of species across the animal kingdom^12,23,32^, including adult and larval *Drosophila*^43,44,144^. They are consistent with a computation of the valence of the US that is predicted by the CS+ (*i.e*. a predicted value associated with the CS+) and could drive the formation of an association between a novel CS and a CS+ during higher-order conditioning.

Additionally, some modulatory neurons acquired CS+ responses that were opposite in sign to their responses to that US (Fig. 7d-iii and 7d-iv). Some of those appear to be activated by the omission of a predicted US whose valence is opposite that of neuron’s preferred US (Fig. 7d-iii). Such responses have been proposed to support extinction by inducing a parallel memory of opposite valence following US omission^9,13,14,53^. Consistent with this idea, in adult flies, DANs of opposite valence and direct cross-compartment MBON-to-DAN connections have been implicated in extinction^13,14,53^, but the role of indirect feedback pathways has not been investigated. In our model we find that removing indirect feedback significantly reduces performance of networks optimized to extinguish a previous association (Fig. 7a). Some modulatory neurons also showed prolonged inhibition in response to the omission of a predicted US whose valence is the same as the neuron’s preferred US (Fig. 7d-iv). Such responses have been proposed to support extinction in mammals by erasing the memory formed by the activation of that modulatory neuron^9,28,145^. Thus, our model raises the possibility that extinction could be implemented via multiple mechanisms in this circuit^9^.

## Discussion

Modulatory neurons (*e.g*. dopaminergic, DAN) are key components of higher-order circuits for adaptive behavioral control, such as the vertebrate basal ganglia or the insect mushroom body (MB), and they provide teaching signals that drive memory formation and updating^12,14,21,24,25,49,58,66^. Here, we provide the first synaptic-resolution connectivity map of a recurrent neural network that regulates the activity of all modulatory neurons in a higher-order learning center, the *Drosophila* larval MB (Fig. 2a-f). We discovered an unexpected component of the insect MB: a large population of 61 feedback neuron pairs that provide one- and/or two-step feedback from the MB output neurons (MBONs) to modulatory neurons (Fig. 2a-d, 3a-b and 5a-e). The majority of these one- and two-step feedback pathways link distinct memory systems, suggesting that the entire MB functions as an interconnected ensemble during learning (Fig. 3b and 5e). We also systematically determined which modulatory neurons evoke aversive and appetitive memories and functionally tested some of the newly identified structural pathways (Fig. 1c-f, 4b-g, and 6a-f). We developed a model of the circuit constrained by the connectome and by our functional data and explored the roles of the newly discovered architectural motifs in different learning tasks (Fig. 7a-e). Our study provides a basis for understanding the circuit implementation of learning algorithms in the tractable insect nervous system.

### Feedback pathways enable adaptive regulation of learning by prior learning

Adaptive regulation of modulatory neuron activity has been proposed to underlie aspects of learning and memory in both vertebrates^31,32,146,147^ and insects^23,42–44^, including long-term memory consolidation^46,49,100,148^, memory reconsolidation^11,13,14^, extinction^13,14,53,149^, and higher-order conditioning^21,150^. However, the architecture and functional principles of the circuits that support this regulation are not well understood. Furthermore, the extent to which modulatory neuron activity is regulated by previous memories in insects is still unclear. Strikingly, we found that many modulatory neurons, including the ones that provide teaching signals for appetitive and aversive olfactory memory formation (Fig. 1c), receive more than 50% of their total dendritic input from feedback pathways that relay MBON signals (Fig. 2d). We confirmed that some of the identified indirect feedback pathways are functional and that feedback neurons can induce memory formation (Fig. 4b-g and 6a-f). These results suggest that prior memories as represented by the pattern of MBON activity can strongly influence modulatory neuron activity in *Drosophila* larva. Indeed, in our connectivity-constrained model, modulatory neuron responses to CS are modified after pairing with US, and this modulation is reduced in the absence of feedback neurons (Fig. 7c-e). While further model constraints are likely required to directly compare model neurons to recordings, the model demonstrates that feedback supports such adaptive responses.

Learning and memory systems in vertebrates^151,152^ and insects^45,47,48,55,84,94,153^ are often organized into distinct compartments implicated in forming distinct types of memories (*e.g*. aversive and appetitive or short- and long-term). However, the extent and nature of interactions between distinct memory systems during memory formation is still an open question. Here, we provide the first comprehensive view of an extensive set of anatomical feedback pathways that can mediate interactions between distinct memory systems (Fig. 3b-c, 5e, and Extended Data Fig. 9b). The cross-compartment feedback pathways identified here suggest that prior memories formed about an odor in one compartment can influence the formation and updating of future memories about that odor in many other functionally distinct compartments.

Interestingly, we also found that the feedback neurons receive input from brain areas other than MB (Extended Data Fig. 4a-c). They could therefore play a role in encoding variables determined by these areas, such as context or internal state. Consistent with this, our model revealed that the performance on a context-dependent conditioning task was significantly reduced in the absence of these neurons (Fig. 7b). The feedback neurons could therefore provide a substrate for flexible and adaptive regulation of learning, based on both previous experience and context or internal state.

### Circuit motifs for computing integrated predicted value signals across aversive and appetitive memory systems

The use of internal predictions to inform future learning can dramatically increase the flexibility of a learning system^28,29^. Indeed, modulatory neurons in vertebrates^12,21,32^ and insects^23^, including adult and larval *Drosophila*^43,44,144^, show adaptive responses consistent with the idea that they encode predictions. However, the circuit properties that permit the computation of predictions are not well understood. In particular, the way in which integrated common-currency predicted value signals across appetitive and aversive memory systems are computed is unclear. Our study reveals candidate circuit motifs that could implement this computation and mediate more complex tasks that require it, such as second-order conditioning. While second-order conditioning has not yet been investigated in the larva, adult *Drosophila* has been shown to be capable of it^20^.

A prominent motif revealed by our analysis of connectivity and neurotransmitter expression is convergence of potentially excitatory and inhibitory connections from MBONs from compartments of opposite valence onto some DANs (Fig. 3b-e). We confirmed that DAN-i1 (whose activation can induce appetitive memory for paired odor) receives functionally inhibitory input from its own compartment and functionally excitatory (likely disinhibitory) input from compartments of opposite valence profile (Fig. 4a-g). By integrating these inputs, such DANs may compute a comparison between the odor-evoked excitation of MBONs in compartments of opposite valence. In naïve animals, odor-evoked MBON excitation in all compartments is thought to be similar. However, associative learning selectively depresses conditioned odor MBON excitation in compartments whose modulatory neuron activation has been paired with the odor^14,66,85,90,133^. The valence predicted by the conditioned odor is therefore thought to be encoded as a skew in the relative excitation of MBONs in compartments of opposite valence. We propose that by comparing the conditioned odor-evoked MBON excitation in compartments of opposite valence via the cross-compartment feedback connections, modulatory neurons could compute an integrated common-currency predicted value signal across appetitive and aversive domains. Our model results are consistent with this idea. Some modulatory neurons in the model acquire responses to conditioned odors that resemble their responses to US (Fig. 7d-ii and 7e), consistent with a value prediction, and these responses, as well as performance on second-order conditioning, were degraded in the absence of cross-compartment feedback (Fig. 7a and 7e).

### Convergence of feedback and US pathways could allow the computation of prediction errors

An important aspect of reinforcement learning theories is the idea that modulatory neurons compare predicted and actual US (compute the so-called prediction errors) and drive memory formation or extinction depending on the sign of the prediction error. Computing prediction errors requires structural convergence between feedback pathways, which carry information about predicted US valence, and afferent US pathways. However, the site of this convergence is still an open question. Feedback and afferent US pathways could converge at the modulatory neurons themselves, upstream of modulatory neurons, or both.

While *Drosophila* modulatory neurons have not yet been directly shown to represent prediction errors, adult and larval *Drosophila* are capable of extinction^5,13,53,141,154^, and our study reveals candidate motifs that could support the comparison of expected and actual US. We found that modulatory neurons receive convergent input from feedback pathways from MBONs and from US pathways (Fig. 2d-f). Modulatory neurons could therefore potentially compute prediction errors by comparing inhibitory drive from the feedback pathways to the excitatory drive from the US pathways, or vice versa, excitatory drive from the feedback pathways and inhibitory drive from the US pathways. Consistent with this idea, we observed in our model some DANs that are inhibited by US alone and activated by CS+ alone, or vice versa (Fig. 7d-iii and 7d-iv).

Our study also revealed that US pathways and feedback pathways converge at two levels: not only at the modulatory neurons themselves, but also at the two-step feedback neurons (FB2Ns, Fig. 2f). Actual and expected outcomes could therefore also be compared by FB2Ns. A recent study in the mouse VTA used retrograde labelling and electrophysiology to characterize the response properties of some of the neurons presynaptic to the DANs^12,41^ and found that a fraction of the analyzed pre-DAN neurons encoded only actual, or only expected reward, while the remainder encoded both variables^41^. Thus, both in vertebrates and in insects, comparing predicted and actual outcomes may be a complex computation involving multiple levels of integration that eventually converges onto an ensemble of modulatory neurons^41^. The connectome of such a network presented here provides a basis for understanding the circuit implementation of this computation.

### The multilevel and cross–compartment feedback increase performance and flexibility

Our connectivity and modeling studies revealed two architectural features of the circuit that provides input to the modulatory neurons that increase its computational performance and flexibility on learning tasks (Fig. 7a-b). The first is the multilevel feedback architecture that includes not only the previously known direct MBON feedback, but also multiple levels of indirect feedback. The second is the extensive set of cross-compartment connections. Our results also reveal modulatory neurons receive a diverse set of feedback inputs (Fig. 2e) that could enable each modulatory neuron to compute a unique set of features. Consistent with this, we observed a diversity of adaptive response types in the modulatory neurons in our model. This suggests that instead of computing a single global reward prediction error that is distributed to all modulatory neurons^21^, the network uses a range of distinct compartmentalized and distributed teaching signals.

In adult *Drosophila*, functional connections between some MBONs and DANs^13,46,49,53–55,153^, as well as between KCs and DANs^48,155^ have been reported, and some of these have been shown to play a role in short-term memory formation^49,153^, long-term memory consolidation^46,54^, re-consolidation^13^, extinction^53^, or in synchronizing DAN ensemble activity in a context-dependent manner^55^. For some of these cases, direct within- or cross-compartment MBON-to-DAN connections have been demonstrated^13,46,49^. While direct axodendritic connections from several MBONs onto DANs exist in the larva^47^ (Extended Data Fig. 9a), we find that indirect connections via the feedback layer account for a much larger fraction of a modulatory neuron’s dendritic input than direct MBON synapses (Fig. 2d), suggesting that adaptive DAN responses may be largely driven by such indirect feedback. Additionally, direct axo-axonic connections from KCs could modulate modulatory neuron output in both larva^47^ and adult^48,155^, but they cannot convey learning-related changes in the strengths of KC-to-MBON connections, so they likely play a very different role to the feedback from MBONs. Since many aspects of connectivity between the core components of the MB (modulatory neurons, KCs and MBONs) are shared between larval and adult *Drosophila* stages and other insects^47,48,67^, we expect that the indirect feedback motifs discovered here are also shared across insects.

Some of the within-compartment feedback motifs we found are reminiscent of the feedback motifs so far described for the DANs in the vertebrate midbrain. Rabies tracing studies have shown that DANs receive input from their direct targets in the striatum (analogous to direct MBON to DAN connections), as well as from the direct targets of striatal neurons (analogous to the one-step feedback described here)^12,39,41,50^. While the diversity and the inputs of striatal feedback neurons have not yet been fully explored, in the future it will be interesting to determine whether many of the striatal feedback neurons also link distinct memory systems.

In summary, we present the first complete circuit diagram of a recurrent network that computes teaching signals in a biological system, providing insights into the architectural motifs that increase the computational power and flexibility of the learning center. Our connectome-constrained model provides numerous predictions that can be tested in the future in a tractable model organism, for which genetic tools can be generated to monitor and manipulate individual neurons^102,134,156,157^. The connectome, together with the functional and modelling studies therefore provides exciting opportunities for elucidating the biological implementation of reinforcement learning algorithms.

## Acknowledgements

We thank Fly Light at HHMI Janelia Research Campus (JRC) for generating confocal images of the GAL4 lines, J. Simpson for sharing a 2-photon microscope, Y. Aso and G. Rubin, for sharing unpublished driver lines, V. Jayaraman for sharing unpublished versions of CsChrimson and GCamp6f, L. Feng and I. Andrade for help with circuit mapping from EM, K. Hibbard and JRC FLY Core, for generating some of the fly stocks, Fly EM at JRC for generating the EM volume, T. Saumweber for discussions, D. Bonnery for help with analyses, L. Abbott for helpful comments on the manuscript, Z. Zavala-Ruiz and the JRC Visiting Scientist Program and HHMI JRC for funding. A.L.-K. was supported by the Burroughs Wellcome Foundation, the Gatsby Charitable Foundation, the Simons Collaboration on the Global Brain, and NSF award DBI-1707398. B.G. received grant support from the Deutsche Forschungsgemeinschaft Ge1091/4-1 and FOR 2705-TP2.

## Materials and Methods

### Fly lines

In the main text and figures, short names are used to describe genotypes for clarity. We used GAL4, Split-GAL4 lines to direct the expression of the red-shifted channel-rhodopsin *20XUAS-CsChrimson-mVenus*^103^ (Bloomington *Drosophila* Stock Center BDSC 55134, gift of V. Jayaraman) or the Calcium indicator *20xUAS-IVS-GCaMP6f*^127^ in pairs of neurons or subsets of neurons. Split-GAL4 lines restrict expression of the effector to a few cells, under the double control of two enhancers (inserted in the attP2 and attP40 docking sites), selected by us or others in Janelia Research Campus (HHMI, VA, USA) based on their GAL4 expression pattern^101,102,158^.

### Modulatory MBINs

We used *SS24765-Split-GAL4* to optogenetically activate OAN-a1 in the calyx. We generated *SS02160-Split-GAL4* to activate DAN-c1 in the lower peduncle. For the vertical lobe, we generated *SS01702-Split-GAL4* to activate or image calcium transients in MBIN-e2 (DAN-c1 was also covered by this line) and *SS01958-Split-GAL4* to activate or image calcium transients in OAN-e1 in the UVL. We used *SS02180-Split-GAL4, MB145B-Split-GAL4* (used for activation and calcium imaging, gift of G. Rubin and Y. Aso) and *MB065B-Split-GAL4*^45^, (which also covered DAN-c1) to target DAN-f1 in the IVL. We used *SS01716-Split-GAL4*^58^ to induce or image DAN-g1 activity in the LVL, and we generated *SS04268-Split-GAL4* to activate OAN-g1, also in the LVL. *MB054B-Split-GAL4* (gift of G. Rubin and Y. Aso) was also used to co-activate DAN-g1 and DAN-f1. We used two lines to target DAN-d1 in the lateral appendix: *MB143B-Split-GAL4* (used for activation and calcium imaging) and *MB328B-Split-GAL4* (both gifts of G. Rubin and Y. Aso). In the medial lobe, we generated a broad line *SS01948-GAL4* which allows coactivation of DAN-h1, DAN-i1, DAN-k1, and sometimes DAN-j1. We also imaged calcium transients in DAN-i17 using the more specific GAL4 *SS00864-Split-GAL4*.

### Neurons presynaptic to the modulatory MBINs

We optogenetically activated multidendritic Class IV neurons (MD IV) with the driver line *ppk-1.9-GAL48* (gift of D. Tracey); Basin interneurons with *GMR72F11-GAL4*^62^; the ascending neuron A00c with *GMR71A10-GAL4*^62,123^ crossed to *ppk-GAL80*^160^, *repo-GAL80*^161^ (to prevent expression in MD IV and glial cells, respectively). We also activated A00c with the more specific GAL4 line *SS00883-Split-GAL4*. We generated *SS01778-Split-GAL4* and *SS02181-Split-GAL4*, which target FB2N-11 and/or FB2N-18. *SS02108-Split-GAL4* targets FAN-7; *SS02401-Split-GAL4* targets FB2N-19.

### Control lines

As a control for the GAL4 lines inserted at the *attP2* site, we used the empty control stock *y w;;attP2*^102,158^ crossed to the effector line. As a control for Split-GAL4 lines with AD at *attP40* and DBD at *attP2*, we used the empty stock *y w;attP40;attP2*^102,158^ crossed to the effector line.

### Lines for recording neuronal activity

Calcium transients in modulatory neurons were imaged using the following constructs to verify functional input of mechano-ch neurons: *w; iav-LexA*^62^ in *attP40; 20xUAS-IVS-GCaMP6f 15.693*^127^ in *attP2, 13XLexAop2-CsChrimson-tdTomato*^103^ in *VK00005*. For Basins multi-sensory interneurons: *w; GMR72F11-LexA*^158^ in *JK22C*; *20xUAS-IVS-GCaMP6f 15.693*^127^ at *attP2, 13XLexAop2-CsChrimson-tdTomato*^103^ at *VK00005*. And for MD class IV nociceptive neurons: *w; 13XLexAop2-CsChrimson-mVenus*^103^ at *attP40* (BDSC 55138); *ppk-1kb-hs43-lexA-GAD10* at *attP2, 20xUAS-IVS-GCaMP6f*^127^ at *VK00005*. All the effectors used in these stocks are a gift from V. Jayaraman. Transvection was tested by bathing some samples in 100 mM mecamylamine and observing the disappearance of responses to optogenetic stimulation (data not shown). If a response remained during mecamylamine application, the experiments were repeated using a spatially defined photo-stimulation using spatial light modulator (SLM) technology (see functional connectivity section for details of the procedure and the lines concerned).

For patch-clamp recording we crossed the genetic driver lines for MBON-m1 (*SS02163-Split-GAL4*) or for MBON-i1 (*SS01726-Split-GAL4*) to *58E02-LexAp65 at attP40*^78^; *13xLexAop2-IVS-GCaMP6f-p10 15.693*^127^ at *VK00005* (BDSC 44276), *20xUAS-CsChrimson-mCherry*^103^ at *su(Hw)attP1* in order to activate MBONs and visualize the medial lobe DANs (ML-DANs) for patch-clamping. Only data for DAN-i1, which was the most frequently hit by the recording pipette, as revealed by post hoc identification, are shown.

The reporter *pJFRC29-10xUAS-IVS-myr::GFP p10*^135^ at *attP2* was used for immunostaining.

### Learning experiments

Learning experiments were performed as previously described^58,47,56^. The larvae were reared in the dark at 25°C in food vials supplemented with 1:200 retinal. The experimenter selected two groups of 30 third-instar larvae and was blind to their specific genotype. The two groups underwent a training procedure involving odor and light exposures, either fully overlapping in time (paired group), or fully non-overlapping (unpaired group). The paired group was placed for 3 minutes on 4% agarose plates and exposed to constant red-light illumination (wavelength: 629 nm, power: 350 *μ*W/cm2; except for *ppk-1.9-GAL4*, which targets neurons at the surface of the body and for which a light power of 35 *μ*W/cm2 was used) paired with the presentation of 12 *μ*l of odor ethyl-acetate (10^-4^ dilution in distilled water) absorbed on two filter papers located on the plate lid. These larvae were then transferred to a new plate with no odor and in the dark for 3 minutes. This paired training cycle was repeated three times in total. The unpaired group of larvae underwent odor presentation in the dark and red light without odor following the same protocol. After a 3-minute test with odor presentation on one side of the plate lid, larvae were counted on the side of the odor, on the opposite side, and in the 1 cm-wide midline of the plate. Preference and performance indices were calculated as in a previous study^57^. Briefly, a preference index (PI) was first computed, for each group as: PI = [N (larvae on the odor side) - N (larvae on the no-odor side)]/N(total), N(total) includes larvae in the middle of the plate. The Learning Performance Score (LPS) was then computed as LPS = [PI (paired) – PI (unpaired)]/2.

### Circuit mapping and electron microscopy

We reconstructed neurons and annotated synapses in a single, complete central nervous system from a 6 hr old female [iso] *Canton S G1 x w1118 [iso] 5905* larva, acquired with serial section transmission EM at a resolution of 3.8 x 3.8 x 50 nm, that was first published along with the detailed sample preparation protocol^62^. Briefly, the CNS was dissected and placed in 2% gluteraldehyde 0.1 M sodium cacodylate buffer (pH 7.4). An equal volume of 2% OsO_4_ was added and the larva was fixed with a Pelco BioWave microwave oven with 350-W, 375-W and 400-W pulses for 30 sec each, separated by 60-sec pauses, and followed by another round of microwaving but with 1% OsO_4_ solution in the same buffer. Next, samples were stained en bloc with 1% uranyl acetate in water and microwaved at 350 W for 3×3 30 sec with 60-sec pauses. Samples were dehydrated in an ethanol series, transferred to propylene oxide, and infiltrated and embedded with Epon resin. After sectioning the volume with a Leica UC6 ultramicrotome, sections were imaged semi-automatically with Leginon^162^ driving an FEI Spirit TEM (Hillsboro, OR), and then assembled with TrakEM2^163^ using the elastic method^164^. The volume is available at https://l1em.catmaid.virtualflybrain.org/?pid=1.

To map the wiring diagram we used the web-based software CATMAID^165^, updated with a novel suite of neuron skeletonization and analysis tools^128^, and applied the iterative reconstruction method^128^. All annotated synapses in this wiring diagram fulfill the four following criteria of mature synapses^62,128^: (1) There is a clearly visible T-bar or ribbon on at least two adjacent z-sections. (2) There are multiple vesicles immediately adjacent to the T-bar or ribbon. (3) There is a cleft between the presynaptic and the postsynaptic neurites, visible as a dark-light-dark parallel line. (4) There are postsynaptic densities, visible as dark staining at the cytoplasmic side of the postsynaptic membrane.

We validated the reconstructions as previously described^62,128^, a method successfully employed in multiple studies^62,124,128,132,166,167^. Briefly, in *Drosophila*, as in other insects, the gross morphology of many neurons is stereotyped and individual neurons are uniquely identifiable based on morphology^167–169^. Furthermore, the nervous system in insects is largely bilaterally symmetric and homologous, with mirror-symmetric neurons reproducibly found on the left and the right side of the animal. We therefore validated neuron reconstructions by independently reconstructing synaptic partners of homologous neurons on the left and right side of the nervous system. With this approach, we have previously estimated the false positive rate of synaptic contact detection to be 0.0167 (1 error per 60 synaptic contacts)^56^. Assuming the false positives are uncorrelated, for an n-synapse connection the probability that all n are wrong (and thus that the entire connection is a false positive) occurs at a rate of 0.0167^*n*^. Thus, the probability that a connection is a false positive reduces dramatically with the number of synaptic contacts contributing to that connection. Even for n = 2 synaptic contacts, the probability that a connection is not true is 0.00028 (once in every 3,586 two-synapse connections). We therefore consider ‘reliable’ connections those for which the connections between the left and right homologous neurons have at least 3 synapses each and their sum is at least 10. See^62,128^ for more details.

### Immunostaining

Dissected brains were fixed in phosphate buffered saline (PBS, NaCl 137 mM, KCl 2.7 mM, Na_2_HPO_4_ 8.1 mM, KH2PO4 1.5 mM, pH7.3) containing 4% paraformaldehyde (Merck) for 30-min at room temperature. After two 15-minute washes with PBT (PBS with 1% or 3% Triton X-100; Sigma-Aldrich), the brains were blocked with 5% normal goat serum (Vector Laboratories) in PBT and incubated for at least 24 hours with primary antibodies at 4°C. Before application of the secondary antibodies for at least 24 hours at 4°C or for 2 hours at room temperature, brains were washed several times with PBT. After that, brains were again washed with PBT, mounted in Vectashield (Vector Laboratories) and stored at 4°C in darkness. Images were taken with a Zeiss LSM 710M confocal microscope. The resulting image stacks were projected and analyzed with the image processing software Fiji^170^. Primary antibodies were used at the following dilutions: rabbit anti-GFP (cat# Af2020, Frontier Institute; 1:1000), chick anti-GFP (ab13970, abcam, 1:1000), rabbit anti-GABA (A2052, Sigma; 1:100), mouse anti-ChAT (ChAT4B1, DSHB Hybridoma Product deposited by P.M. Salvaterra; 1:50). Rabbit anti-DVGlut^137^ was diluted 1:1000. Secondary antibodies were used at the following dilutions: Alexa Fluor 568-conjugated goat anti-rabbit IgG (A-11036, Invitrogen Molecular Probes; 1:300), Alexa Fluor 633-conjugated goat anti-mouse IgG (A-21050, Invitrogen Molecular Probes; 1:300) and Alexa Fluor 488-conjugated goat anti-chicken IgG (A-11039, Invitrogen Molecular Probes; 1:300).

### Identifying GAL4 lines that drive expression in modulatory neurons and their presynaptic partners

To identify GAL4 lines (listed in Supplementary Table 1) that drive expression in specific neurons, we performed single-cell FlpOut experiments (for FlpOut methodology see ^62,171^) of many candidate GAL4 lines^134^. We generated high-resolution confocal image stacks of individual neuron morphology (multiple examples per cell type). Most MBONs and MBINs were uniquely identifiable based on the dendritic and axonal projection patterns (which MB compartment they project to and the shape of input or output arbor outside the MB). These were also compared to previously reported singlecell FlpOuts of dopaminergic and octopaminergic neurons in the larva^57,95,123,172,173^. For the neurons upstream of MBINs (FBNs/FANs/FB2Ns), we used morphology and cell body position to identify the lineage of the neuron. The precise shape and 3D location of dendritic and axonal projections were then examined and compared to all potential candidates in the lineage which have been fully reconstructed from the electron microscopy volume. In some cases, two neurons had very similar morphology at both light and EM level, and in such cases they also had nearly identical connectivity (e.g. FB2N-11 and FB2N-18).

### Functional connectivity assays

Central nervous systems were dissected in a cold buffer containing 103 mM NaCl, 3 mM KCl, 5 mM TES, 26 mM NaHCO_3_, 1 mM NaH_2_PO_4_, 8 mM trehalose, 10 mM glucose, 2 mM CaCl_2_, 4 mM MgCl_2_ and adhered to poly-L-lysine (SIGMA, P1524) coated cover glass in small Sylgard (Dow Corning) plates.

For optogenetic activation, red illumination (617nm High-Power Lightguide Coupled LED Source, Mightex) was positioned above the sample to depolarize the axon terminal parts of the sensory neurons (MD IV or chordotonal) or the second order interneurons (Basins). Light stimulations were performed with 1 or 15 sec duration and in 40 and 600 cycles of laser on/off pulses of 20 msec/5 msec. Each preparation underwent three types of light stimulation of increasing power: ca. 390 μW/mm^2^, 920 μW/mm^2^ and 4.6 mW/mm^2^. Only the data for the highest light power during 1 sec is displayed (Fig.1f). The same stimulus was spaced with 30 sec for a total of three presentations in each scan. Each scan consisted in imaging dopaminergic neurons on a two-photon scanning microscope (Bruker) using a 60x 3 1.1 NA objective (Olympus). A mode-locked Ti:Sapphire laser (Chameleon Ultra II, Coherent) tuned to 925 nm was used for photo-activation of the GCaMP6f. Fluorescence was collected with photomultiplier tubes (Hamamatsu) after band-pass filtering. Images were acquired in line scanning mode (5.15 fps) for a single plane of the CNS.

To overcome transvection observed between the transgenes at the attP40 landing site of the *MB143B-Split-GAL4* line (targeting DAN-d1) crossed to *w; 13XLexAop2-CsChrimson-mVenus*^103^ in attP40; ppk-1kb-hs43-lexA-GAD10 in attP2, 20xUAS-IVS-GCaMP6f2 in VK00005, we used 3-dimension spatially defined photo-stimulation. MD IV neurons expressing CsChrimson were photoactivated by a holographic pattern generated by a two-photon 1040 nm laser (femtoTrain, Spectra-Physics) coupled to a phase-only SLM (Intelligent Imaging Innovations). GCaMP6f signal was imaged by a laser tuned to 925 nm (Insight DS+ Dual, Spectra-Physics). The optogenetic stimulations were 50 cycles of laser on/off pulses of 2 msec/18 msec, ranging from 1 to 1.5 mW/mm^2^. Off-target (equidistant from the Chrimson-expressing DAN-d1 neuron, but not targeting Chrimson-expressing MD IV neurons) and on-target stimulations were alternatively performed and the difference between transvection-only generated calcium signals and transvection + MD IV neuron activation-generated signal was computed and used as the fluorescence signal. DAN-d1 neurons were imaged at a frame rate of ca. 5 fps on a two-photon scanning microscope (Vivo, Intelligent Imaging Innovations) using a 25x 21.1 NA objective (Nikon).

For image analysis, image data were processed by Fiji software^170^ and analyzed using custom code in Matlab (The Mathworks, Inc). Specifically, we manually determine the regions of interest (ROIs) from maximum intensity projection of entire time series images, and measure the mean intensity. In all cases, changes in fluorescence were calculated relative to baseline fluorescence levels (F0) as determined by averaging over a period of at least 2 sec. just before the optogenetic stimulation. The *δ*F/F_0_ values were calculated as *δ*F/F_0_ = (F_*t*_-F_0_)/F_0_, where F_*t*_ is the fluorescent mean value of a ROI in a given frame. Analyses were performed on the mean *δ*F/F_0_ of the consecutive 3 stimulations.

### Whole-cell patch-clamp recordings from DANs on optogenetic activation of MBONs

For recording, the isolated brain attached with VNC were dissected from third instar larvae in Baines external solution^174^, which contained (mM): 135 NaCl, 5 KCl, 2 CaCl_2_.2H_2_O, 4 MgCl_2_.6H_2_O, 5 2-[(2-Hydroxy-1,1-bis(hydroxymethyl)ethyl)amino] ethanesulfonic acid, 5 N-[Tris(hydroxymethyl) methyl]-2-aminoethanesulfonic acid, and 36 sucrose. The pH was adjusted to 7.15 with NaOH, and osmolarity was 310-320 mOsm. The preparation was viewed with a 60x 1 NA water-immersion objective equipped with an Olympus microscopy (BX51WI; Olympus). GCaMP6f–labeled DANs were visualized with a 470-nm wavelength LED. The glial sheath above the targeted DANs was ruptured using 0.1% protease (Protease XIV; Sigma-Aldrich). Recording electrodes were pulled from thick-wall glass pipet (O.D. 1.5mm, I.D. 0.86mm) using P-97 puller (Sutter Instruments) and fire-polished to resistances of 10–15 M*ω*. The Baines intracellular solution^174^ contained (mM): 140 potassium gluconate, 5 KCl, 2 MgCl_2_.6H_2_O, 2 EGTA, 20 HEPES. The pH was adjusted to 7.4 with KOH, and the osmolarity was 280 mOsm. Biocytin was dissolved in intracellular solution at 0.5% for further post hoc morphological identification of recorded DANs. The data were acquired and processed using Digidata 1550, Multiclamp 700B, and Clampex 10.4 software (Molecular Devices). The recording was sampled at 20 kHz and filtered at 6 kHz under current-clamp mode. CsChrimson was activated by 617-nm wavelength LED.

#### DAN identification

After the electrophysiology recording, the preparation containing the VNC and brain was fixed in 4% paraformaldehyde in 0.1 M phosphate buffer saline (PBS) overnight at 4°C, and then transferred to PBS until staining. After rinsing in PBS, the CNS preparations were placed in Streptavidin Alexa Fluor 647 (1:200) in PBS with 10% Triton X (overnight, room temperature). After rinsing, the preparations were dehydrated and mounted with DPX. The confocal images were captured with Zeiss 800 confocal laser microscope. Alexa Fluor 647 was excited with 633 nm-wavelength light, and mCherry-tagged CsChrimson neurons were excited with 567 nm-wavelength light.

### Statistical analysis

As most fluorescence and behavioral data were non-normally distributed (according to a Shapiro-Wilk test), we opted for non-parametric tests for paired comparisons.

For behavioral experiments, the performance scores obtained for each line tested in optogenetic reinforcement were compared to the ones of its corresponding empty line (i.e. *w;;attP2* or *w;attP40;attP2* for GAL4 or Split GAL4, respectively) using a non-parametric Mann-Whitney U test for independent sets of data. For multiple comparisons, the probability values were compared to a threshold of 0.05 adjusted with a Holm-Bonferroni correction to balance for Type I and Type II statistical errors, unless otherwise stated. Across GAL4 lines, comparisons of performance scores were done using the same methodology. Data were plotted using the Matlab script *errorbarjitter*, available at http://www.mathworks.com/matlabcentral/fileexchange/33658-errorbarjitter.

Fluorescence analyses were done using a non-parametric Wilcoxon test for paired comparisons between the maximum *δ*F/F_0_ plus one standard deviation during 1 sec before photostimulation onset and the maximum *δ*F/F_0_ at two time windows: during the 1 sec of the stimulation, and from 1 to 3 seconds after its onset.

For the clustering analysis, we looked for clusters among FBNs/FANs based on the similarity of their synaptic partners separately for input and output. To find clusters based on synaptic inputs, we defined the similarity between a pair of FBN/FANs as the cosine similarity of the vector of inputs they receive from MBONs where the weight of a given connection is measured as the fraction of total input synapses on the postsynaptic neuron. Specifically, for *v_i_* and *v_j_* being the input vectors for FBN/FANs *i* and *j*, the similarity between them is defined 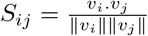. Hierarchical clustering on the similarity matrix was done with Scipy using average linkage. We chose the top five clusters to highlight, which included all clearly differentiated groups of FBN/FANs. Clustering on the output patterns was done identically using the vectors of connectivity from FBN/FANs onto MBINs.

For the input-clustered groups, we assessed the similarity of the patterns of synaptic outputs and vice versa for the synaptic input patterns for output-clustered groups. We measured the overall group similarity as the median of all unique pairwise cosine similarities between neurons within the group. We used a permutation test to assess the significance of the observed similarities by randomizing the relationship between input pattern and output pattern for each FAN/FBN. For example, for each input-clustered group of size n, we randomly chose *n* output patterns and computed their median output similarity in the same way. A one-sided p-value was computed from the distribution of 10,000 random permutations with a Holm-Sidak correction for multiple comparisons across the groups.

### Rate model of the MBON-i1-FBN-7-DAN-i1 one-step feedback motif for Extended Data Fig. 11b

To illustrate the potential effects of different FBN-baselines we modeled the isolated MBON-i1-FBN-7-DAN-i1 feedback motif shown in Fig. 4b with rate equations where the output of neuron type (MBON, FBN, DAN), changed over time according to the equation 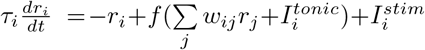 where 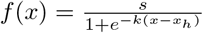, *W* was a matrix with positive and negative values corresponding to the direct interactions between neurons as shown in the circuit schematic of Fig. 4b, 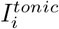 was a nonnegative tonic input into neuron *i*, 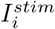 is a stimulus input provided only to MBON, *τ_i_* is a time constant, and parameters *s, k* and *x_h_* set the shape of the sigmoidal response. Equations were solved using *ode45* in Matlab (The Mathworks, Inc).

### Connectivity-constrained model of the entire mushroom body with the feedback neurons

#### Model dynamics

We constructed a recurrent network model of the larval MB containing MBONs, DANs and other feedback neurons. The network receives input from 70 KCs, and external cues, such as US. The normalized firing rate **r**_*i*_ of neurons *i* is modeled as a threshold-linear function of its input:

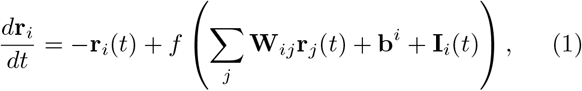

where *f* represents positive rectification. Time is modeled in units of effective time constant (representing combined synaptic and membrane timescales). The connectivity matrix **W**_*ij*_ is constrained using the EM reconstruction. The vector **b**^*i*^ represents the static bias input to each neuron which determines its excitability, while **I**_*i*_(*t*) represents time-varying external input. For MBONs, this includes external input from KCs, 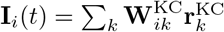.

KCs are initially silent, but during the presentation of an odor CS, the activity of a random fraction *f* of KCs is set to 1, leading to MBON activation. We assume all-to-all KC-to-MBON connectivity. Weights **W**^KC^ are initially set equal to their maximum value of 1/(*N*_KC_*f*), but are modified according to a DAN-dependent synaptic plasticity rule. A weight *W*(*t*) from KC *k* to an MBON in compartment *i* evolves according to:

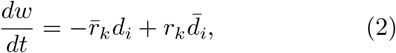

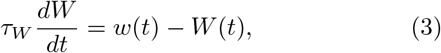

where *d_i_* represents the level of dopamine in the compartment (a weighted sum of DAN inputs according to the DAN-to-MBON connectivity matrix), and *r_k_* represents the firing rate of the KC (note that modifications of weights onto MBONs depend only on KC and DAN activity). The terms 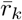 and 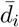 represent the firing rate *r_k_* and dopamine level *d_i_*, respectively, low-pass filtered with time constant *τ*, which leads to an anti-Hebbian timing-dependent synaptic weight update in Equation 2. The second equation results in *W*(*t*) following these updates with a time constant of *τ_W_* (Equation 3). For simplicity, we assume that all modulatory neurons induce plasticity according to this rule.

Weights among DANs, MBONs, and feedback neurons are constrained by the EM reconstruction. Weight matrices are initialized using synapse counts from the EM data, scaled so that the *ℝ*_2_ norm of the inputs received by each neuron 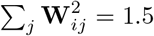. Only reliable connections, as defined previously, are included. Weights from neurons known to communicate using an inhibitory neurotransmitter, are then multiplied by −1. As optimization progresses, weights from neurons of known neurotransmitter identities are constrained to maintain a consistent sign by clipping at 0. At the beginning of a trial, MBON rates are initialized to 0 while DAN and feedback neuron rates are initialized to 0.1. This promotes networks in which MBONs are primarily odor-driven, but some DANs and feedback neurons exhibit baseline levels of activity.

#### Tasks

Neuron *i*’s external input **I**_*i*_(*t*) represents either KC input in the case of MBONs (as described above), or US or contextual signals (depending on the task) in the case of DANs and FB neurons. We assume that 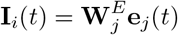, where **W**^*E*^ is initialized as a random standard Gaussian variable and **e**_*j*_(*t*) = 0 or 1 depending on whether signal *j* is active. For most tasks, there are two signals (positive or negative US).

A linear readout of the MBONs determines the preference index via 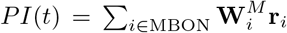, where **W**^*M*^ is initialized as a random Gaussian variable with variance 1/*N*_MBON_. Entries of **W**^*M*^ corresponding to MBONs whose activation is known to produce positive or negative PIs are constrained to be consistent with this sign.

Trials consist of 80 time units. In a first-order conditioning trial, a CS+ is presented for 3 time units starting randomly between *t* = 5 and *t* = 15, followed by a positive or negative US with a delay of 2 time units. A test CS+ presentation occurs between *t* = 65 and *t* = 75, and the system must output the appropriate PI of +1 or −1 depending on the US valence during this second presentation. For extinction, an additional CS+ presentation occurs randomly between *t* = 35 and *t* = 45, and the magnitude of the PI is halved for the final test CS+ presentation. For second-order conditioning, a new CS2 is presented at this time, followed by the original CS+, and the test occurs for CS2. Finally, for context-dependent conditioning, a contextual signal that determines the US valence is presented 3 time units prior to the first CS. At *t* = 30 and *t* = 60 firing rates are reset to their initial conditions to model an arbitrary time delay between CS presentations and preventing networks from using persistent activity, rather than synaptic plasticity, to maintain associations.

For networks trained on first-order conditioning, second-order conditioning, and extinction, training consists of random second-order conditioning and extinction trials (for which first-order conditioning is a subcomponent). On each trial, there is a 50% probability that one of the signals (e.g. the US) will be omitted, or a CS-odor will replace a CS+ odor, and the network report a PI of 0 in these cases, ensuring that only valid CS-US contingencies are learned.

#### Optimization

The network parameters, including all weights except for KC-to-MBON weights, as well as the biases **b**, are optimized using PyTorch using the RMSprop optimizer (www.pytorch.org). Optimization consists of 1500 epochs of 30 trials each. The cost to be minimized is equal to the squared distance between the actual and target PI summed over timesteps, plus a regularization term for DAN activity. The regularization term equals 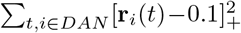, which penalizes DAN activity that exceeds a baseline level of 0.1. We used a timestep of Δ*t* = 0.5, although we verified that our qualitative results hold for smaller timesteps.

| Parameter | Notation | Value |
| --- | --- | --- |
| KC coding level | $f$ | 0.1 |
| Max. KC-to-MBON synaptic weight | $w_{\text{max}}$ | $1/(N_{\text{KC}}f)$ |
| Timing-dependent plasticity window | $\tau$ | 5 |
| Timescale of weight modifications | $\tau_W$ | 5 |
| Initial MBON rate | $m_0$ | 0 |
| Initial DAN rate | $d_0$ | 0.1 |
| Initial FB neuron rate | $x_0$ | 0.1 |
| CS/US presentation length | $T_{\text{stim}}$ | 3 |
| CS-US delay | $\Delta T_{\text{US}}$ | 2 |
| Trial length | $T$ | 80 |
| Timestep | $\Delta t$ | 0.5 |
| RMSprop learning rate | $\eta$ | 0.002 |
| Batch size | $B$ | 30 |
| Number of epochs | $N_{\text{epochs}}$ | 1500 |

---

**Extended Data Figure 1:**
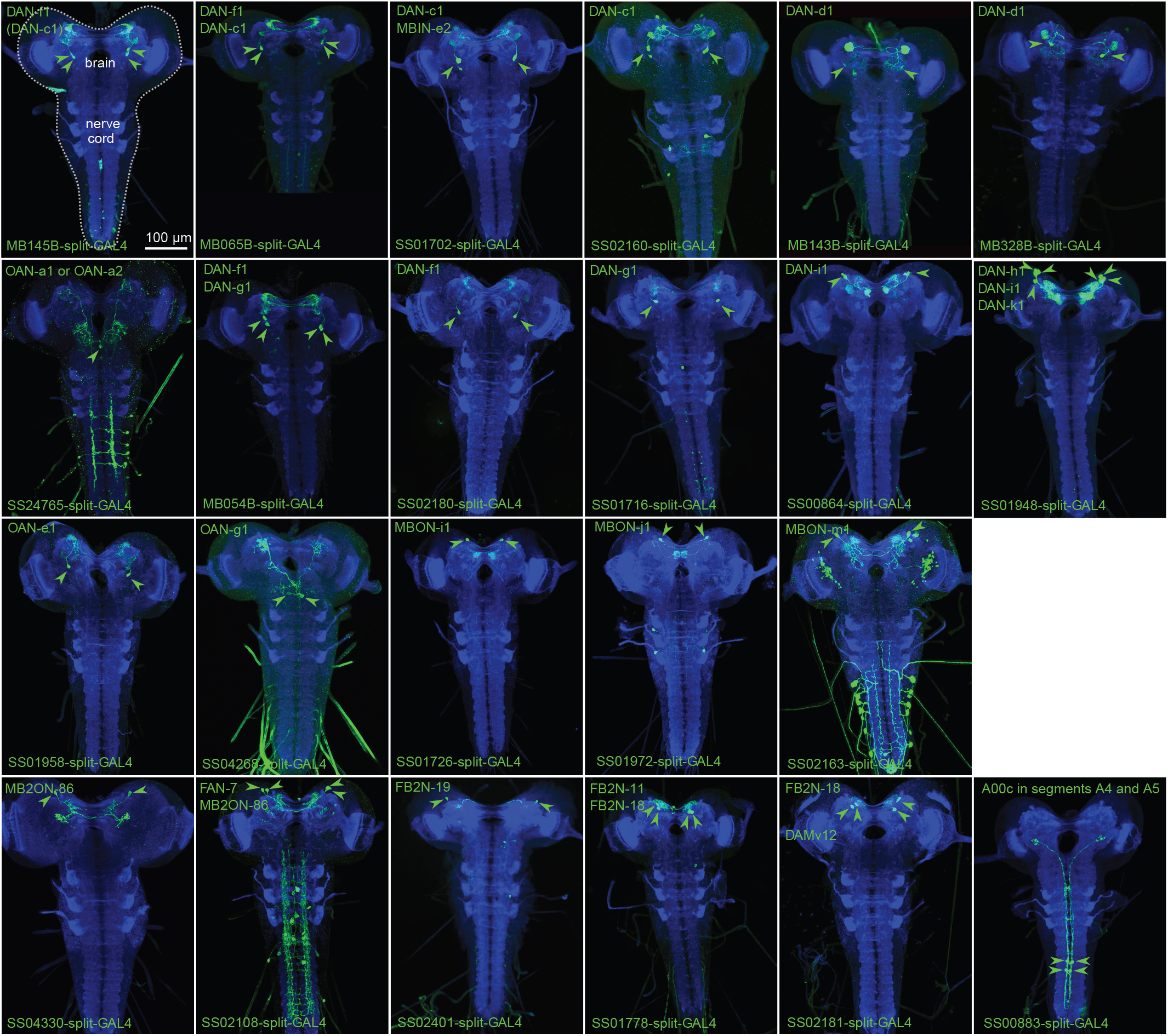
Expression patterns of Split-GAL4 lines. Each panel shows a confocal maximum intensity projection of the complete CNS of third-instar larvae (indicated by the dotted line in the first pannel), with the neuropil labeled with anti-N-Cad antibody (blue) and the Split-GAL4 line expression pattern revealed by driving UAS-myr-GFP (green). Arrowheads indicate cell bodies of identified neurons.

**Extended Data Figure 2:**
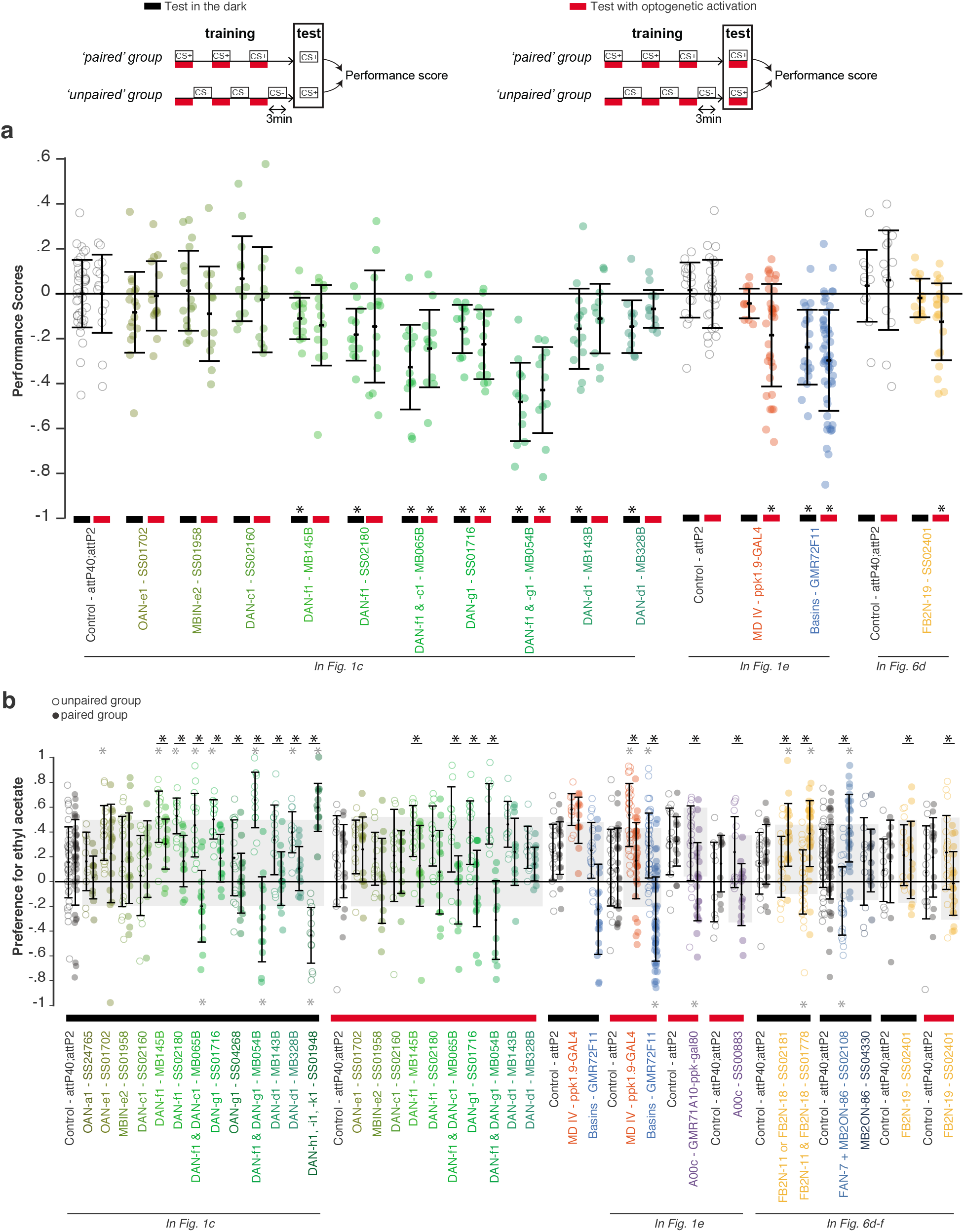
Detailed characterization of associative memories formed through different kinds of “optogenetic punishments” or “optogenetic rewards”. **a** With some natural punishments, aversive memory is behaviorally expressed by trained *Drosophila* larvae only if the punishment is present at the moment of the test (Hendel and Gerber, 2006; Eschbach *et al*. 2011; Schleyer *et al*. 2011). Here we assayed olfactory aversive memories in two ways: both with or without optogenetic punishment (*red and black bars*, respectively) during the retention test. We found that, aversive memory formed by DAN activation (*green*) was expressed to the same extent with or without the DANs activated during the retention test. Similarly memories evoked by Basins activation (*blue*) can be expressed without activation of Basins during test. However memory evoked by the activation of nociceptive MD IV neurons (*orange*) or FB2N-19 (*yellow*) was fully expressed only if these neurons were active again during the retention test. Mean and standard deviations are shown, *: p-value from a Mann-Whitney test comparison to the scores of the corresponding control group (*open circles*) compared to the value 0.05 adjusted with a Holm-Bonferroni correction for multiple comparisons. **b** Preference scores for the trained odor, ethyl acetate, when it was paired (paired group, *closed circles*) or not paired (unpaired group, *open circles*) with optogenetic punishments or rewards. Odor preference was decreased and increased, respectively, relative to genetic controls, after pairing the odor with the presence and absence of the following optogenetic punishments: coactivation of the aversive DAN-f1 and DAN-g1, co-activation of DAN-f1 and DAN-c1, or activation of Basins. On the contrary, odor preference was increased and decreased, respectively, relative to genetic controls, after pairing the odor with the presence and absence of the following optogenetic rewards: the co-activation of DAN-h1, -i1, and -k1 (*dark green);* the activation of FB2N-18 and FB2N-11 (*yellow*), or activation of FAN-7 (*blue-gray*). Thus, both absence of odor in the unpaired group of animals, as well as the presence of odor in the paired group of animals can be associated with the activation of some DANs or some of their afferent neurons. For other DANs or afferent neurons, only paired (e.g. A00c, *purple*)) or only unpaired (e.g. the modulatory DAN-f1, the nociceptive MD IV sensory neuron, or FB2N-19) contingency significantly affected odor preference with respect to the control group. Ether of these two observed types of effects can contribute to the negative or positive learning performance indexes plotted in *Fig. 1c, 1e, 5f’, 5g’* and *5h’. Black* *: p-value<0.05 from a Wilcoxon test comparison between paired and unpaired group. *Grey* *: p-value<0.05 from a Mann-Whitney U test comparison between the preference scores for a given group (paired or unpaired) and the preference scores (for paired or unpaired protocol, respectively) obtained by the control line shown on the left of each set of data. Sample sizes: N = **42**, 11, 17, 16, 12, 14, 12, 13, 12, 16, 12, 14, 12, 12, **15**, 14, 12, 11, 14, 13, 12, 14, 11, 11, 11, **18**, 11, 20, **25**, 33, 52, **14**, 21, **14**, 14, **18**, 18, 31, **52**, 27, 11, **11**, 13, 10, 20 (control groups in bold).

**Extended Data Figure 3:**
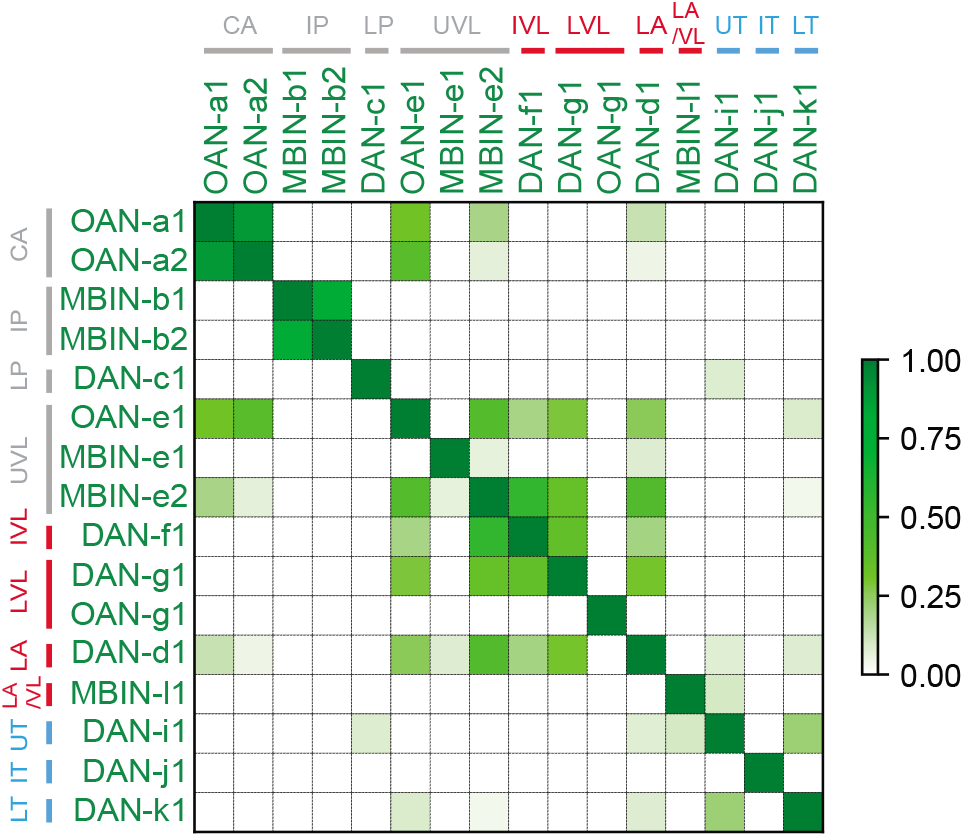
Matrix of similarity between modulator neurons based on the amount of common input. Similarity is obtained by counting the total number of inputs onto a row modulator neuron that are also inputs of the column modulator neurons, and divide by the total number of inputs onto the column modulator neurons. An input here is a connection, consisting typically of many synapses, from a specific cell type onto the modulator neuron. Inputs onto a modulator neuron type are considered if the pair of left and right neurons presynaptic to the pair of left and right modulator neurons is each above a threshold of 1% (*e.g*. the presynaptic neuron makes 3 synapses onto a neuron with 300 postsynaptic sites) and the sum of both is over 3.3% (*e.g*. the sum of both connections is above 10 synapses for receiving neurons with 300 postsynatic sites).

**Extended Data Figure 4:**
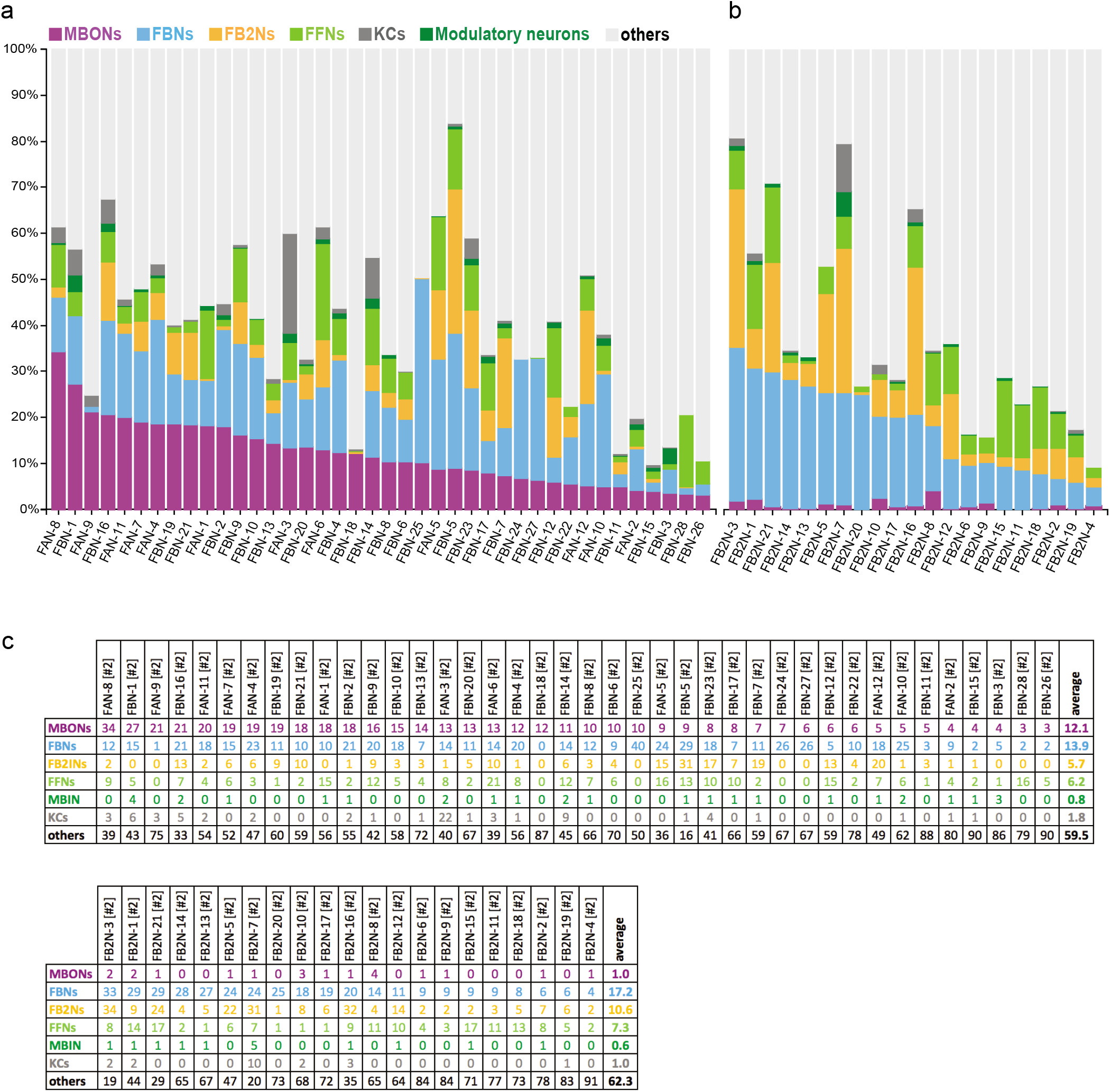
Input onto feedback neurons. Figure shows the fractions of total dendritic input each pre-modulatory neuron (FBN, FB2N or FFN) receives from KCs, modulatory neurons, MBONs, FBNs, FB2Ns, FFNs, and from other non-MB neurons (others). a FBNs receive on average 12% of their inputs directly from MBONs and most of them also receive inputs from other FBNs, with an average of 26% from MBONs and other FBNs combined (see also *Extended Data Fig. 13a*). b FB2Ns receive inputs both from FBNs (on average 17%) and from other FB2Ns (on average 28% from FBNs and FB2Ns combined). Many feedback neurons also receive a significant fraction of input from other unknown neurons from other brain areas (other than MB), suggesting that the feedback about the learnt valences of stimuli is integrated with or modulated by other information. c Tables show percent of inputs onto FBNs (*top*) and FB2Ns (*bottom*) from MBONs, FBNs, FB2Ns, FFNs, modulatory neurons, and Kenyon cells.

**Extended Data Figure 5:**
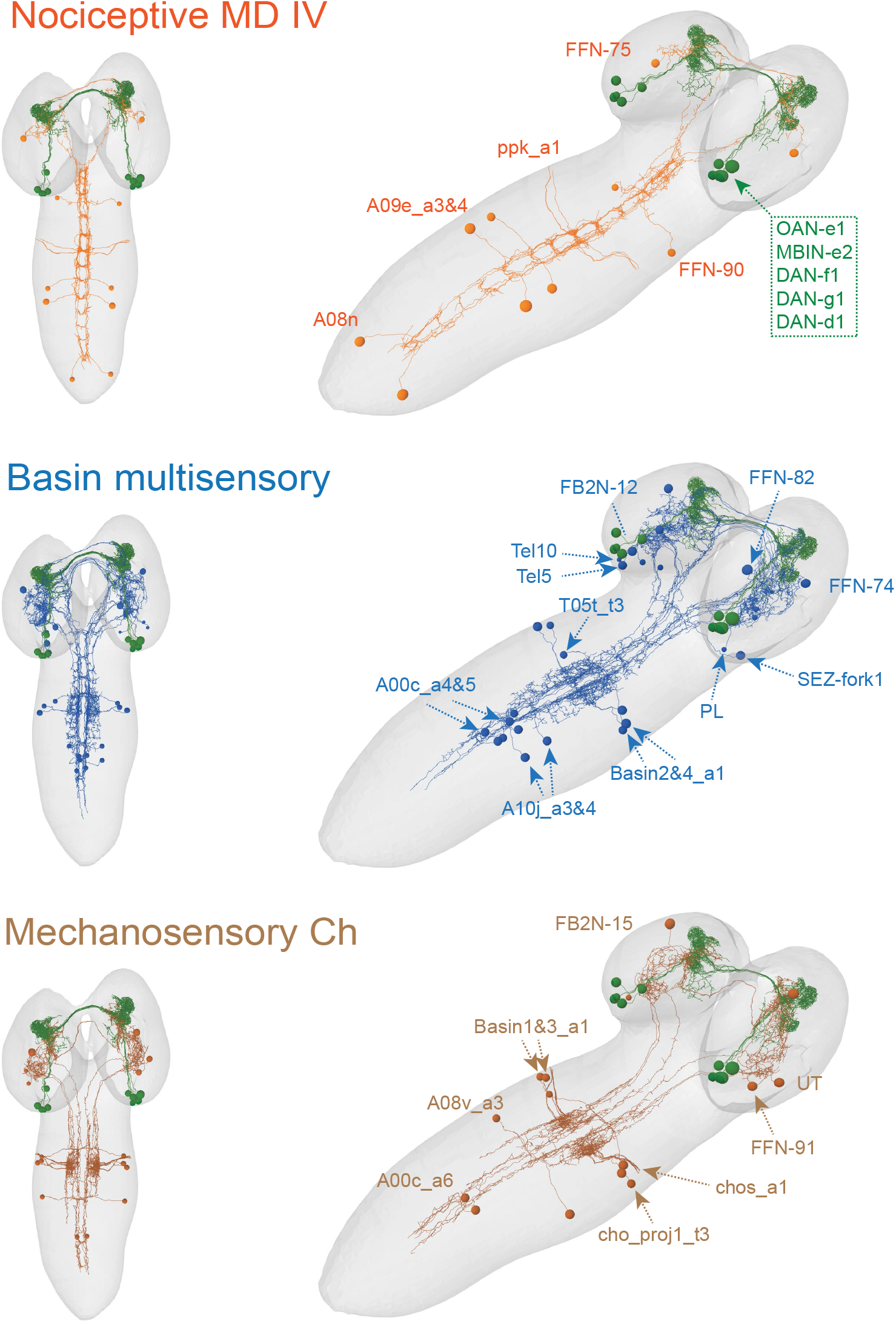
(*related to Fig. 2f*) EM reconstruction of feedforward pathways from sensory neurons to the VL modulatory neurons. Figure shows reconstructed neurons in the nociceptive (*orange*), multisensory Basin (*blue*) and mechanosensory (*brown*) pathways projecting to the VL modulatory neurons (*green*).

**Extended Data Figure 6:**
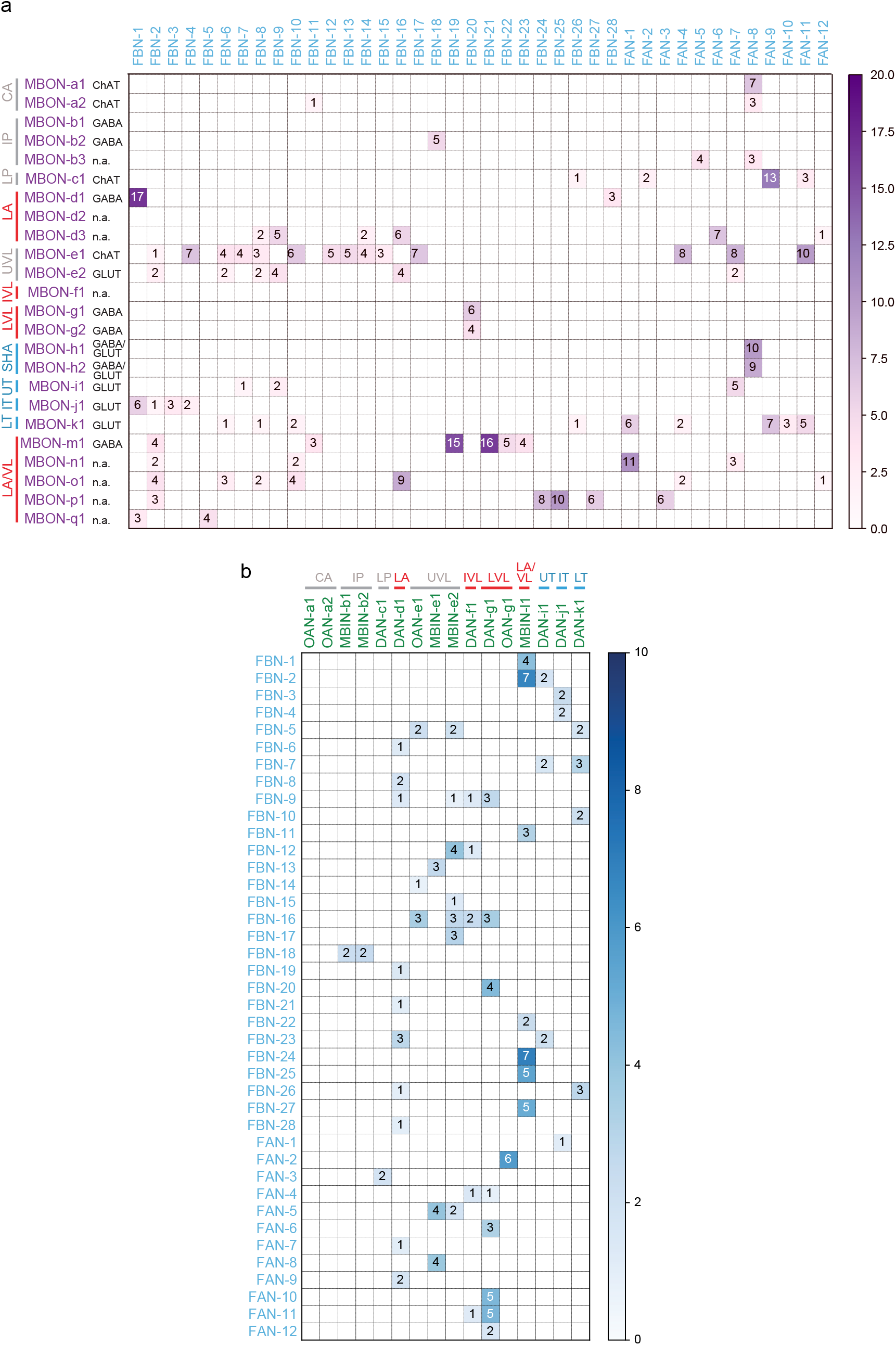
(*related to Fig. 3*) Connectivity matrices between MBONs, FBNs, and modulatory neurons. Connectivity matrix showing normalized synaptic input (expressed as % input, computed as in *Fig. 2b*) each postsynaptic neuron (*columns*) receives from each presynaptic neuron (*rows*). Only reliable connections are shown for which both the left and right homologous connections have at least 3 synapses, and their sum is at least 10, and for which the postsynaptic neuron receives at least 1% of input from the presynaptic neuron. MBONs and modulatory neurons are grouped by MB lobe. CA, Calyx; IP, Intermediate peduncle; LP, Lower peduncle; LA, Lateral appendix; UVL, Upper vertical lobe; IVL, Intermediate vertical lobe; LVL, Lower vertical lobe; SHA, Shaft; UT, Upper toe; IT, Intermediate toe; LT, Lower toe. **a** Each row and column represents MBONs (presynaptic neurons) and FBNs (postsynaptic neurons), respectively. **b** Each row and column represents FBNs (presynaptic neurons) and modulatory neurons (postsynaptic neurons), respectively.

**Extended Data Figure 7:**
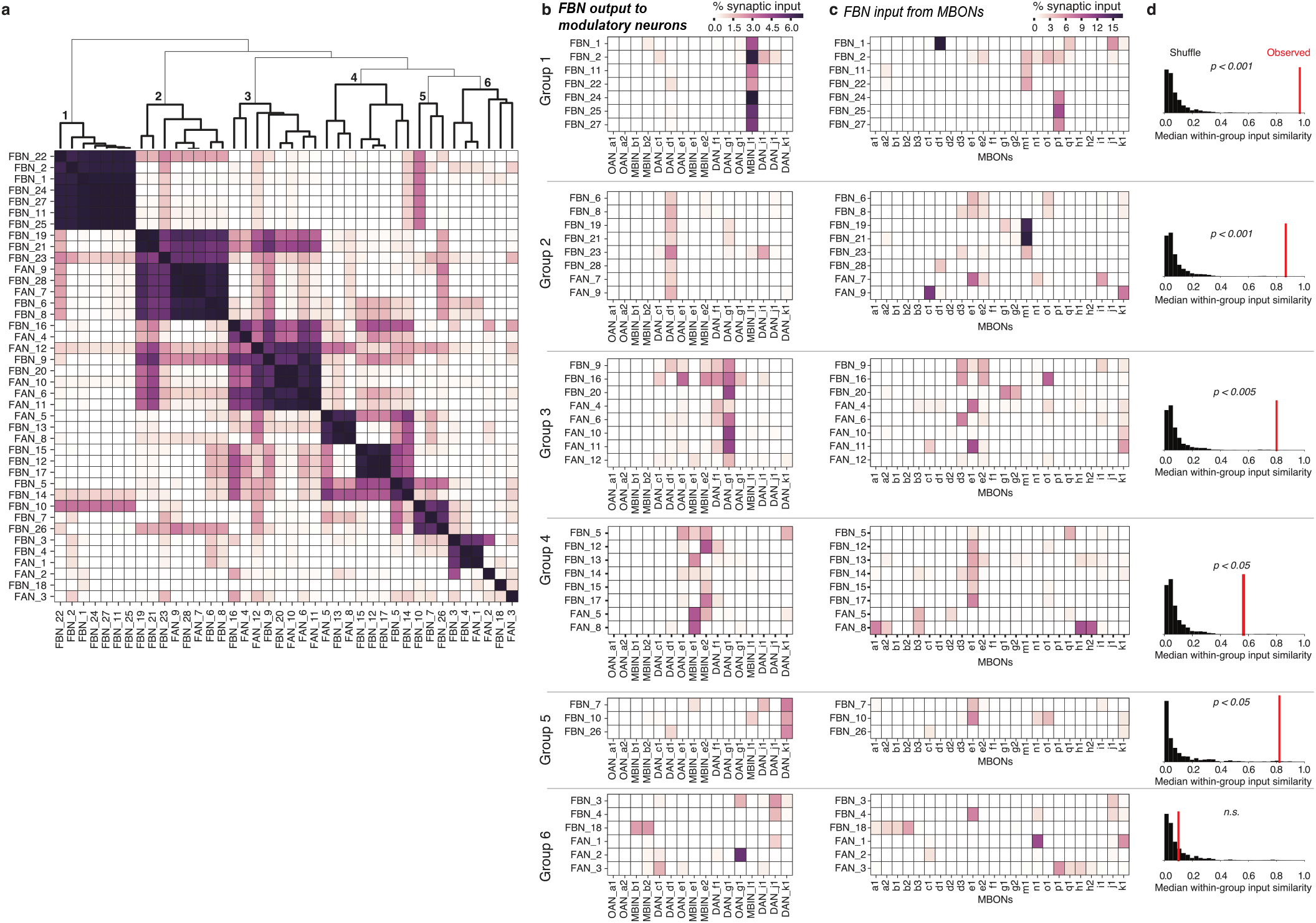
Clustering FBNs based on output onto modulatory neurons. **a** Heat map of FBN similarity based on the pattern of FBN synaptic output across all modulatory neurons. The similarity between a pair of FBNs was computed as the cosine similarity between the vectors of normalized synaptic output onto all modulatory neurons. Indices were ordered by agglomerative clustering with average linkage (dendrogram shown at top). We highlight six groups of FBNs defined by similarities in their output patterns (bold lines in dendrogram, numbered). **b** Heat maps showing patterns of synaptic output from FBNs to modulatory neurons for output groups highlighted in **a**. Each group corresponds to several FBNs strongly targeting one or a small number of modulatory neurons, suggesting that some modulatory neurons are more strongly modulated than others. **c** Heat maps showing patterns of input onto FBNs from MBONs for the output groups highlighted in **a**. **d** The observed similarity in the input patterns between FBNs within each group, compared to shuffled data. For each group (as defined by output patterns), we computed the observed median of cosine similarity of the input vectors across all pairs of neurons (red line). In Groups 1–5, the neurons clustered by outputs had more input output patterns than would be expected by chance. To determine significance, we compared the observed similarity to the distribution of the median cosine similarity for randomly permuted samples from the observed population of input vectors (black histograms, n=10000 randomized trials). A Holm-Sidak correction was applied to p-values to correct for multiple comparisons.

**Extended Data Figure 8:**
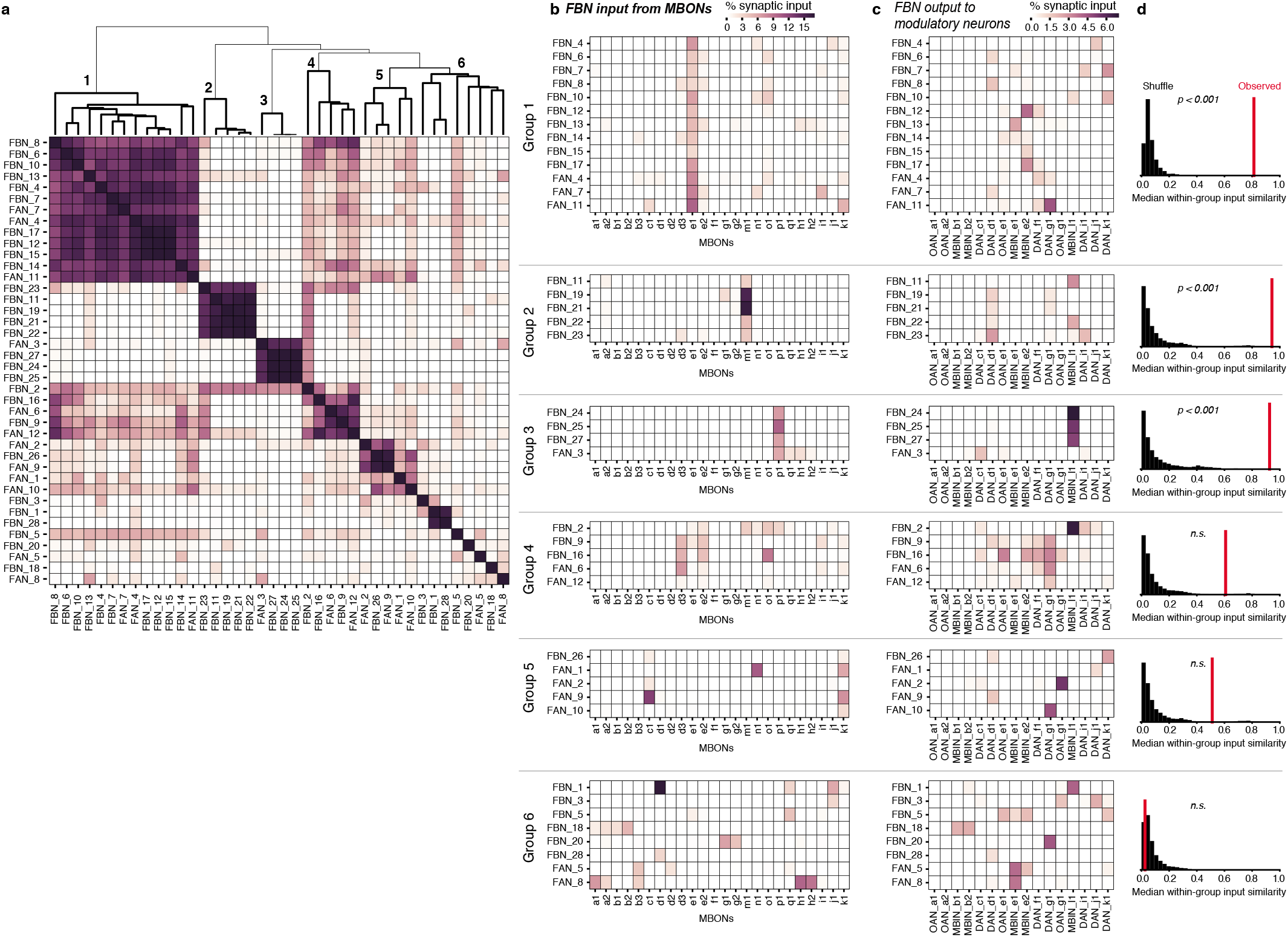
Clustering FBNs based on input from MBONs. **a** Heat map of FBN similarity based on the pattern of FBN synaptic inputs from MBONs. The similarity between a pair of FBNs was computed as the cosine similarity between the vectors of their normalized synaptic inputs from all MBONs. Indices were ordered by agglomerative clustering with average linkage (dendrogram shown at top). We highlight six groups of FBNs defined by similarities in their input patterns (bold lines in dendrogram, numbered). **b** Heat maps showing patterns of input from MBONs onto FBNs for the input groups highlighted in **a**. In all cases, connectivity is measured in normalized synaptic input on the postsynaptic neuron. Most input groups receive dominant input from a single specific MBON (Groups 1,2, 5) or small group of MBONs (Groups 3 and 4), while Group 6 is not well-clustered and contains a variety of dissimilar input patterns. **c** Heat maps showing the patterns of synaptic output from FBNs to modulatory neurons for the input groups highlighted in **a**. **d** The observed similarity in the output patterns between FBNs within each group, compared to shuffled data. For each group clustered by input pattern, we computed the observed median of cosine similarity of the output vectors across all pairs of neurons (red line). In Groups 1,2, and 3, the neurons clustered by inputs had more similar output patterns than would be expected by chance. To determine significance, we compared the observed similarity to the distribution of the median cosine similarity for randomly permuted samples from the observed population of output vectors (black histograms, n=10000 randomized trials). A Holm-Sidak correction was applied to p-values to correct for multiple comparisons.

**Extended Data Figure 9:**
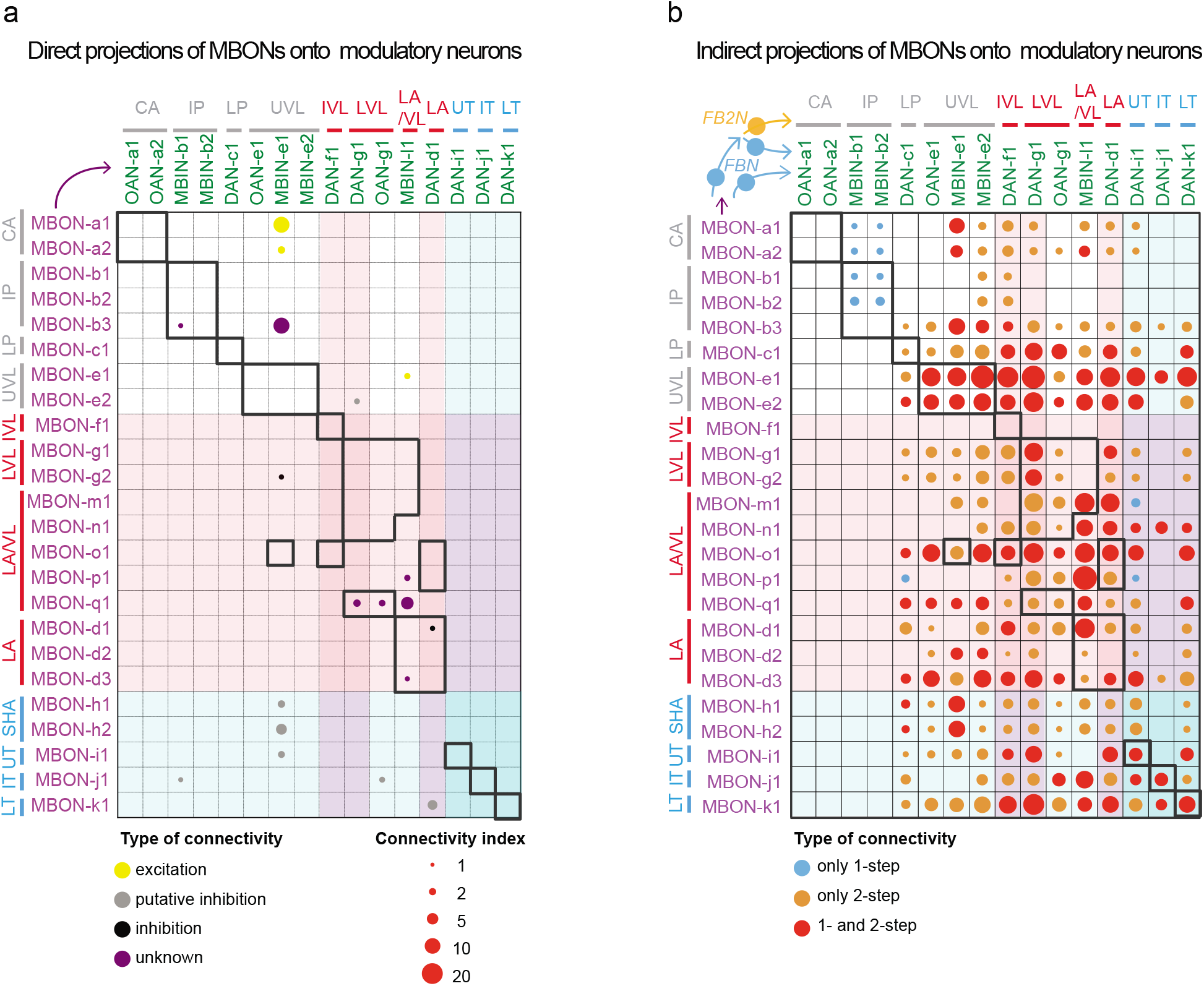
Direct MBONs to modulatory neuron connectivity is very sparse, in contrast to the very dense connectivity via one-and two-step feedback pathways. **a** Connectivity matrix showing normalized synaptic input (expressed as % input, computed as in *b*) each modulatory neuron (*columns*) receives from each MBON (*rows*). Only reliable connections for which the postsynaptic neuron receives at least 1% of input from the presynaptic neuron are shown. When the neurotransmitter of the MBON is known, the circle is color-coded to represent type of connection: excitatory (ChAT) or probably disinhibitory (GluT). Color shades represent the valence of the memory formed in a given compartment (red: aversive memory, blue: appetitive memory). True within-compartment feedback connections from an MBON that receives direct synaptic input from that modulatory neuron are boxed in bold. Very few modulatory neurons receive direct input from MBONs, in contrast to the dense connectivity between MBONs and modulatory neurons via the indirect one- and two-step feedback pathways (b). **b** Connectivity matrix showing indirect connections between MBONs and modulatory neurons via one-step and/or two-step feedback pathways. The matrix was obtained by summing the matrices from *Fig. 3b* and *Fig. 5e*. The color indicates the type of indirect connection existing between a given MBON and a given DAN. Bubble size represents a connectivity index computed as in *Fig. 3b* and *Fig. 5e*. A connectivity index of 1 or 10 means that for all connections comprising that indirect feedback pathway the presynaptic neuron accounts for 1% and 10% of input onto that postsynaptic neuron, respectively. One- and two-step feedback drastically increases the connectivity between MBONs and modulatory neurons, compared to direct connections (*a*).

**Extended Data Figure 10:**
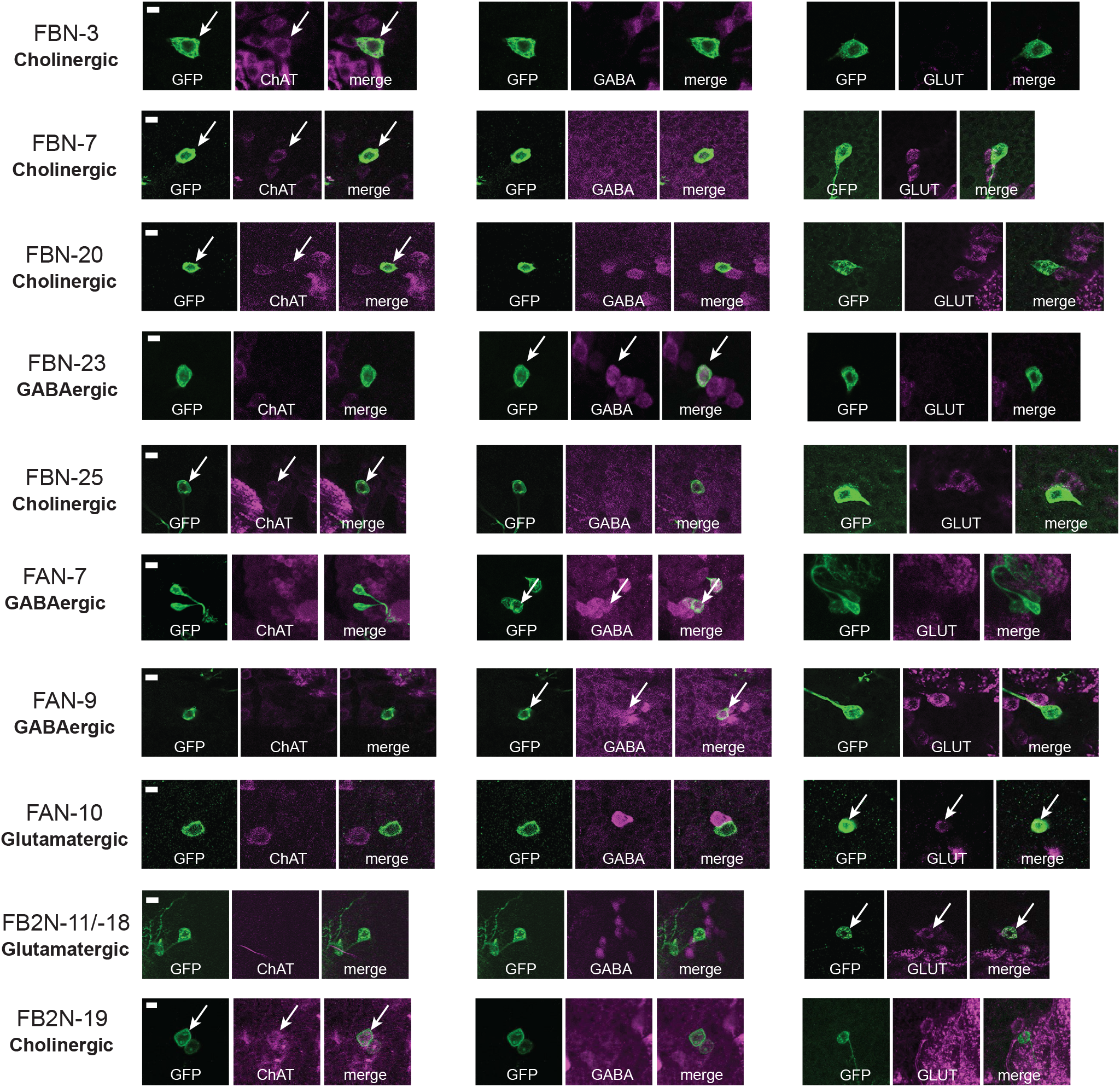
Identification of neurotransmitters expressed in some FBNs/FB2Ns. Neurotransmitter expression detected in neuron somata using antibody labelling. We identified GAL4 lines that drive gene expression in some of the FBN or FB2N neurons and used them to express GFP in these neurons. We stained central nervous systems with antibodies against GFP and either ChAT (choline acetyltransferase), GABA (gamma aminobutyric acid) or GLUT (vesicular glutamate transporter). Each row shows from left to right: the name of the individual neuron, anti-GFP (green), anti-ChAT (magenta), and both antibody stainings combined; anti-GFP (green), anti-GABA (magenta), and both antibody stainings combined; anti-GFP (green) and anti-GLUT (magenta), and both antibody stainings combined. Whether a cell is cholinergic, GABAergic or glutamatergic is listed at the beginning of each row under the neuron name. Images show confocal maximum intensity projections of specific neuronal cell bodies. Scale bars: 5 *μ*m.

**Extended Data Figure 11:**
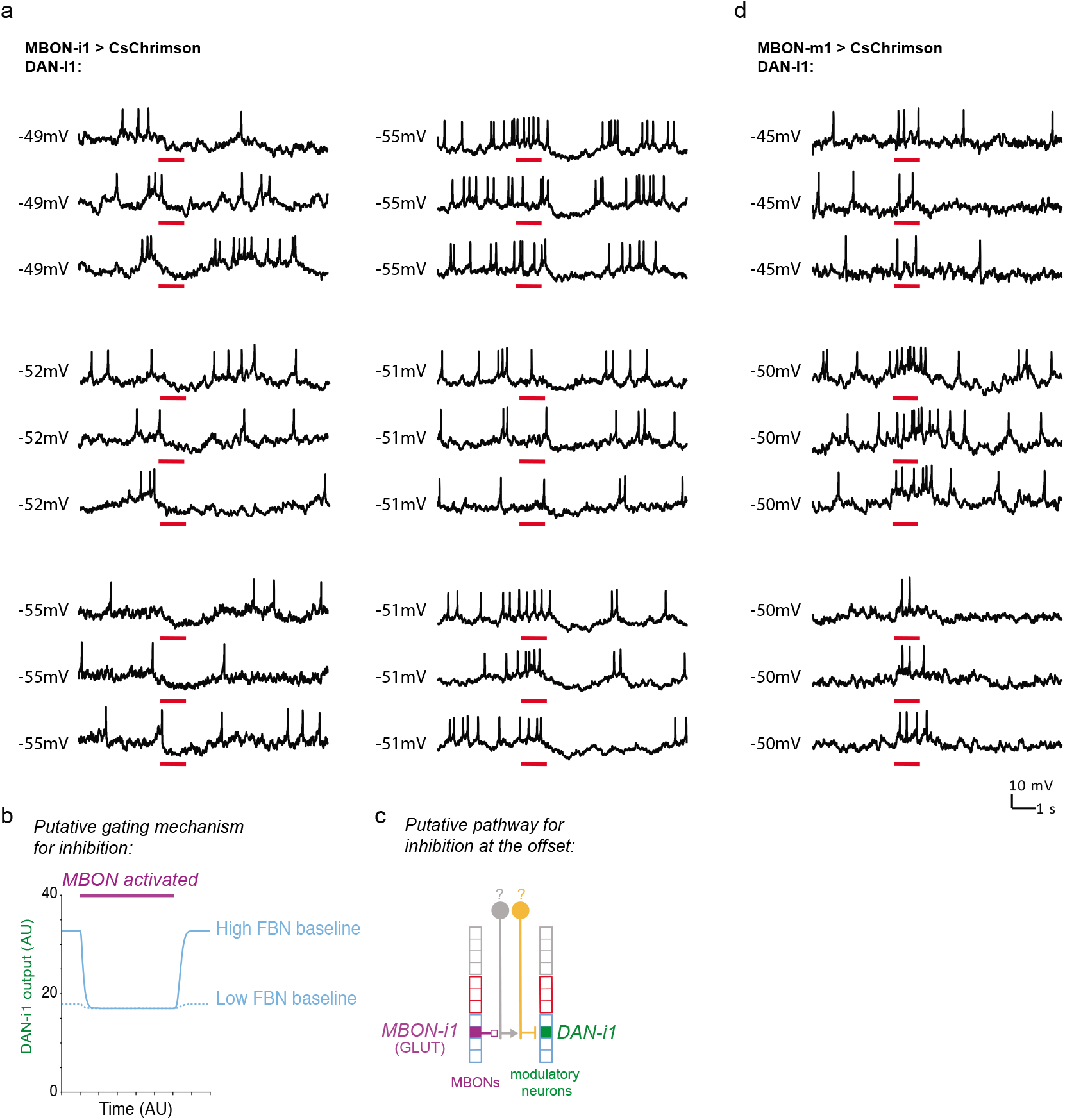
(*related to Fig. 4*) Electrophysiological recording of DAN-i1 during optogenetic stimulation of MBON-i1 or MBON-m1. **a** Raw electrophysiological traces of whole-cell patch-clamp recording of DAN-i1 (medial lobe) during optogenetic activation of the medial lobe MBON-i1 of the same compartment. Left: 3/9 animals had a long-latency inhibitory responses in DAN-i1 at the onset of MBON-i1 activation (55.3 ± 17.3 ms, n=60 traces from 3 animals). For each of them, three individual traces are shown. Inhibition at the onset in some animals but not others might result from distinct baseline states of FBN, as modeled in **b**. Right: 4/9 animals had even longer latency inhibitory responses in DAN-i1 at the offset of MBON-i1 activation (95.3 ± 43.5 ms, n=80 trials from 4 animals). Traces from 3 example animals are shown. Inhibition at the offset might be result of post-inhibitory rebound within a two-step feedback pathway like the one shown in **c**. **b** Simple rate model of the circuit comprising the MBON-i1, FBN-7 and DAN-i1, as shown in *Fig. 4b* illustrating how distinct FBN-7 baseline states could result in an inhibitory response, or no inhibitory response to the onset of MBON-1 activation in DAN-i1. Neuronal activity was modeled as a leaky integrator with logistic function response. The high and low baseline states are modeled as a stronger or weaker tonic excitatory input into the FBN-7. Purple bar indicates MBON activation. In the model, in the high-FBN-7 and low-FBN-7 baseline states, onset of MBON-1 activation evokes an inhibitory response and no response in DAN-i1, respectively. c Schematic diagram showing a putative two-step feedback pathway between MBON-i1 and the DAN-i1 that could mediate inhibition at the offset of MBON-i1 activity. This proposed mechanism would involve post-inhibitory rebound in the neuron postsynaptic to MBON-i1. An example two-step feedback pathway that could mediate this response comprises FBN-9 and FB2N-3, but the neurotransmitters of these neurons have not yet been identified due to the lack of appropriate GAL4 lines. **d** Raw electrophysiological traces of whole-cell patch-clamp recording of DAN-i1 (medial lobe) during optogenetic activation of the vertical lobe MBON-m1. Three out of three animals had a long-latency excitatory response in DAN-i1 to optogenetic activation of MBON-m1 (51.3 ± 7.7 ms, n=45 trials from 3 animals). For each of them, three individual traces are shown. Red rectangles indicate time of optogenetic stimulation. Numbers indicate absolute baseline potential.

**Extended Data Figure 12:**
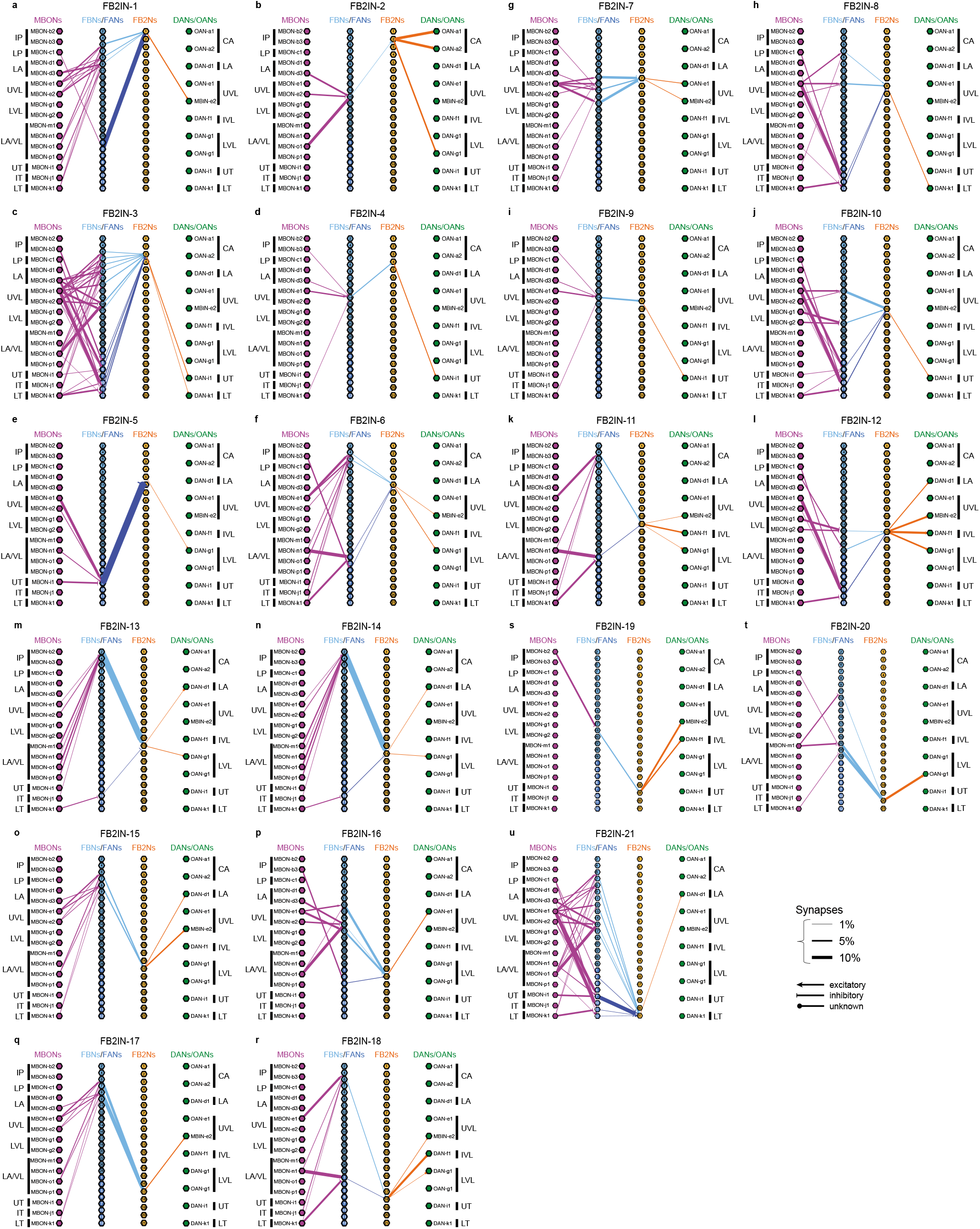
(*related to Fig. 5*) All identified within- and cross-compartment two-step feedback pathways, via FB2Ns. 21 FB2N pairs receive indirect input from MBONs and direct input from FBNs and synapsed onto modulatory neurons. Some FB2Ns synapsed onto modulatory neurons in their own compartment (true second-order feedback), as well as in other compartments, while other FB2Ns synapsed onto modulatory neurons in other compartments (pure second-order feed-across pathways). Thickness of the arrows is proportional to normalized synaptic input (as in (*Fig. 2e-f*). Arrowhead, line, square, and circle denote excitatory (ChAT), inhibitory (GABA), probably inhibitory (GLUT), and unknown neurotransmitter identity, respectively.

**Extended Data Figure 13:**
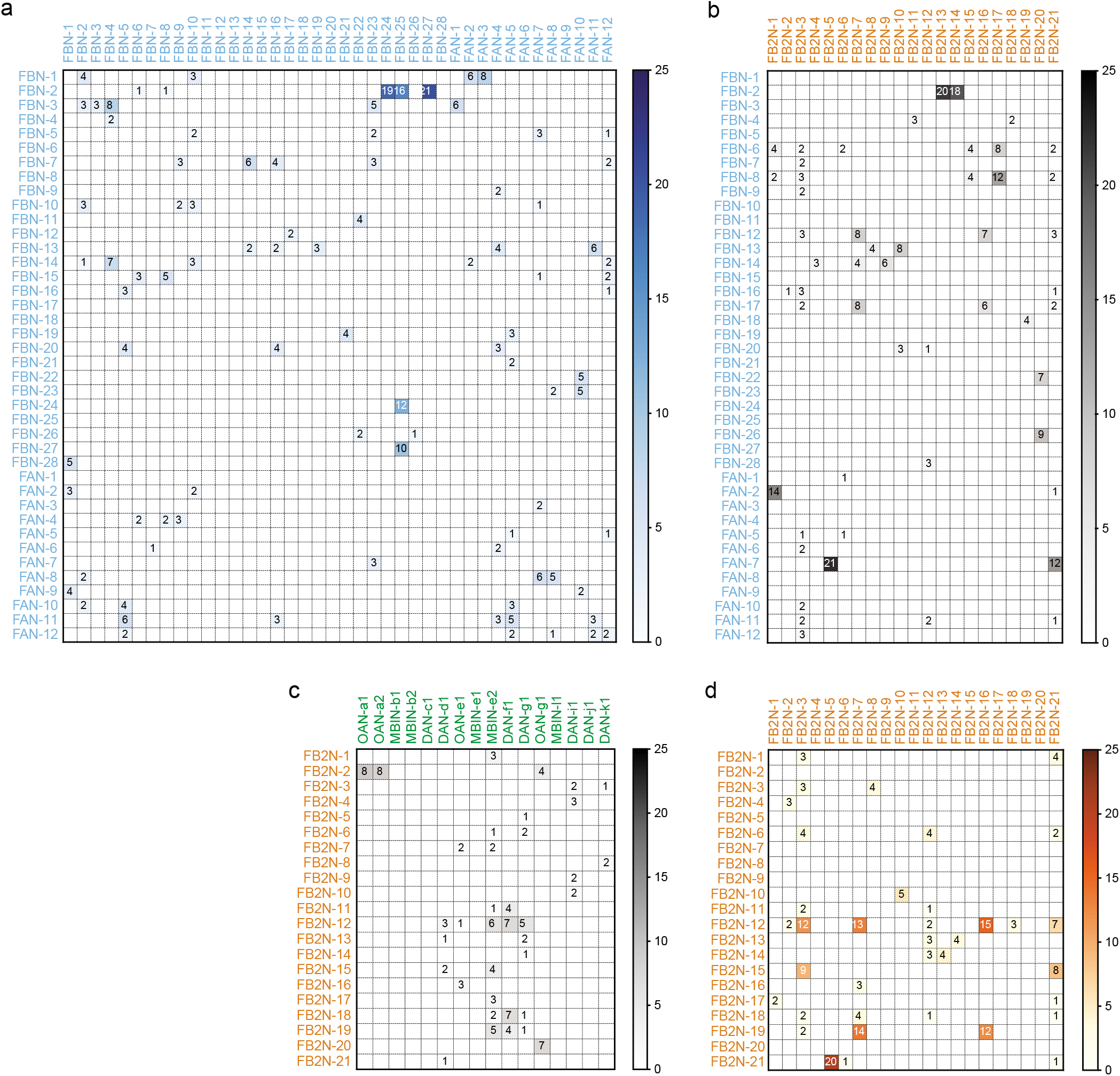
(*related to Fig. 5*) Connectivity matrices of feedback neurons with each other and with modulatory neurons. Connectivity matrix showing normalized synaptic input (expressed as % input, computed as in *Fig. 2b*) each postsynaptic neuron (*columns*) receives from each presynaptic neuron (*rows*). Only reliable connections are shown for which both the left and right homologous connections have at least 3 synapses, and their sum is at least 10, and for which the postsynaptic neuron receives at least 1% of input from the presynaptic neuron. **a** FBNs (rows) synapsing onto FBNs (*columns*). **b** FBNs (rows) synapsing onto FB2Ns (*columns*). **c** FB2Ns (rowss) synapsing onto FB2Ns (*columns*). **d** FB2Ns (*rows*) synapse onto modulatory neurons (*columns*). Modulatory neurons are grouped by their compartments and lobes. CA, Calyx; IP, Intermediate peduncle; LP, Lower peduncle; LA, Lateral appendix; UVL, Upper vertical lobe; IVL, Intermediate vertical lobe; LVL, Lower vertical lobe; SHA, Shaft; UT, Upper toe; IT, Intermediate toe; LT, Lower toe.

**Extended Data Figure 14:**
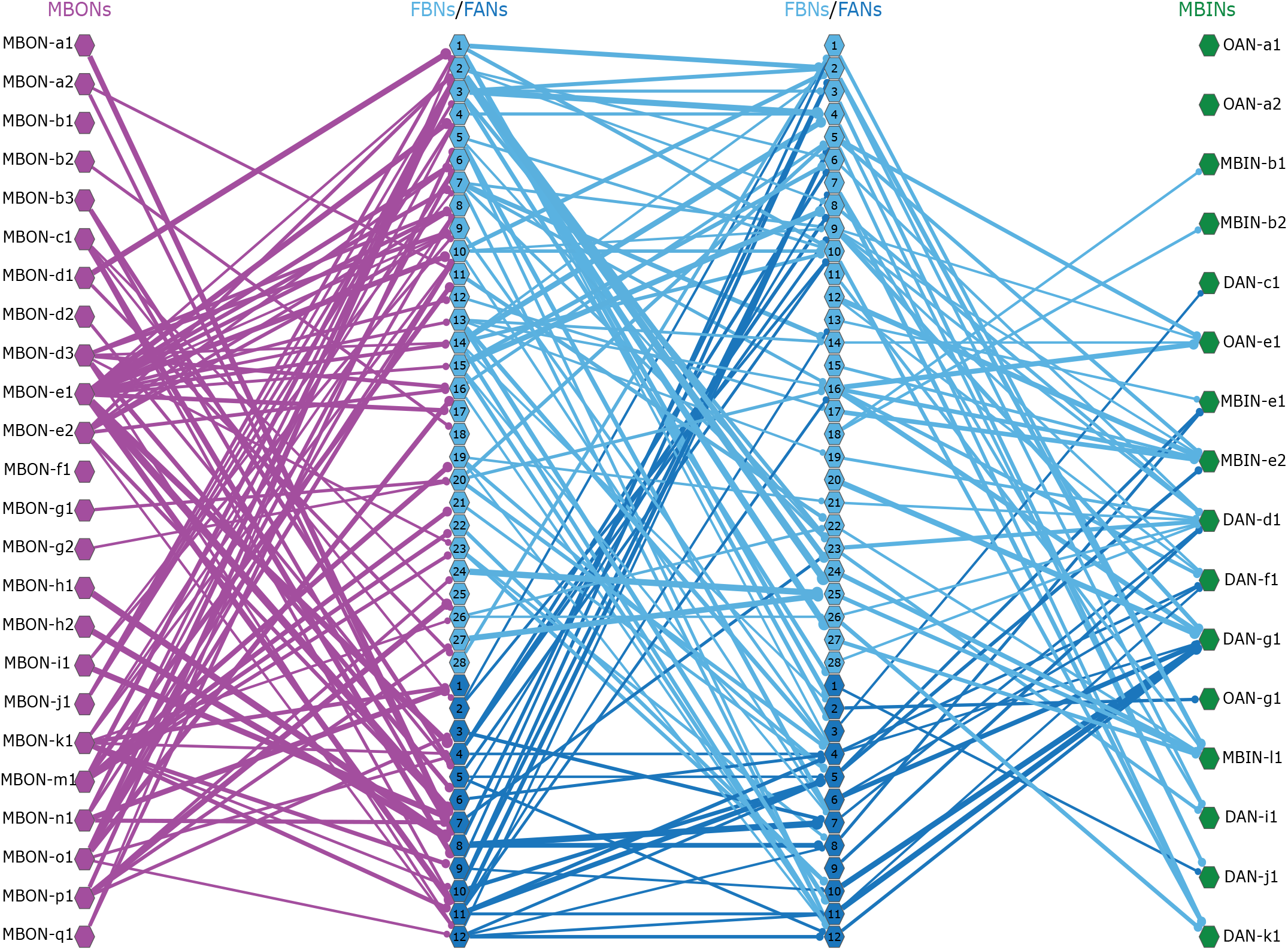
(*related to Fig. 5*) Two-step feedback via FBNs. The interconnections among FBNs enable them to also provide two-step (in addition to one-step) feedback to modulatory neurons, similarly to FB2Ns (but the latter by definition do not receive direct inputs from MBONs). Left-right homologous neurons have been grouped as a type, and only connections with 10 or more synapses are shown. With this stringent connectivity criterium, almost all identified FBNs participate in two-step feedback motifs, except for FBN-12, FBN-13, FBN-15, FBN-18, FBN-20, FBN-27, FBN-28 and FAN-9, which do not receive inputs from other FBNs. All modulatory neurons receive two-step feedback via FBNs, except for OAN-a1 and OAN-a2.

