## Supplementary Files for "Multilevel feedback architecture for adaptive regulation of learning in the insect brain"

#### Supplementary Atlas

### FBNs

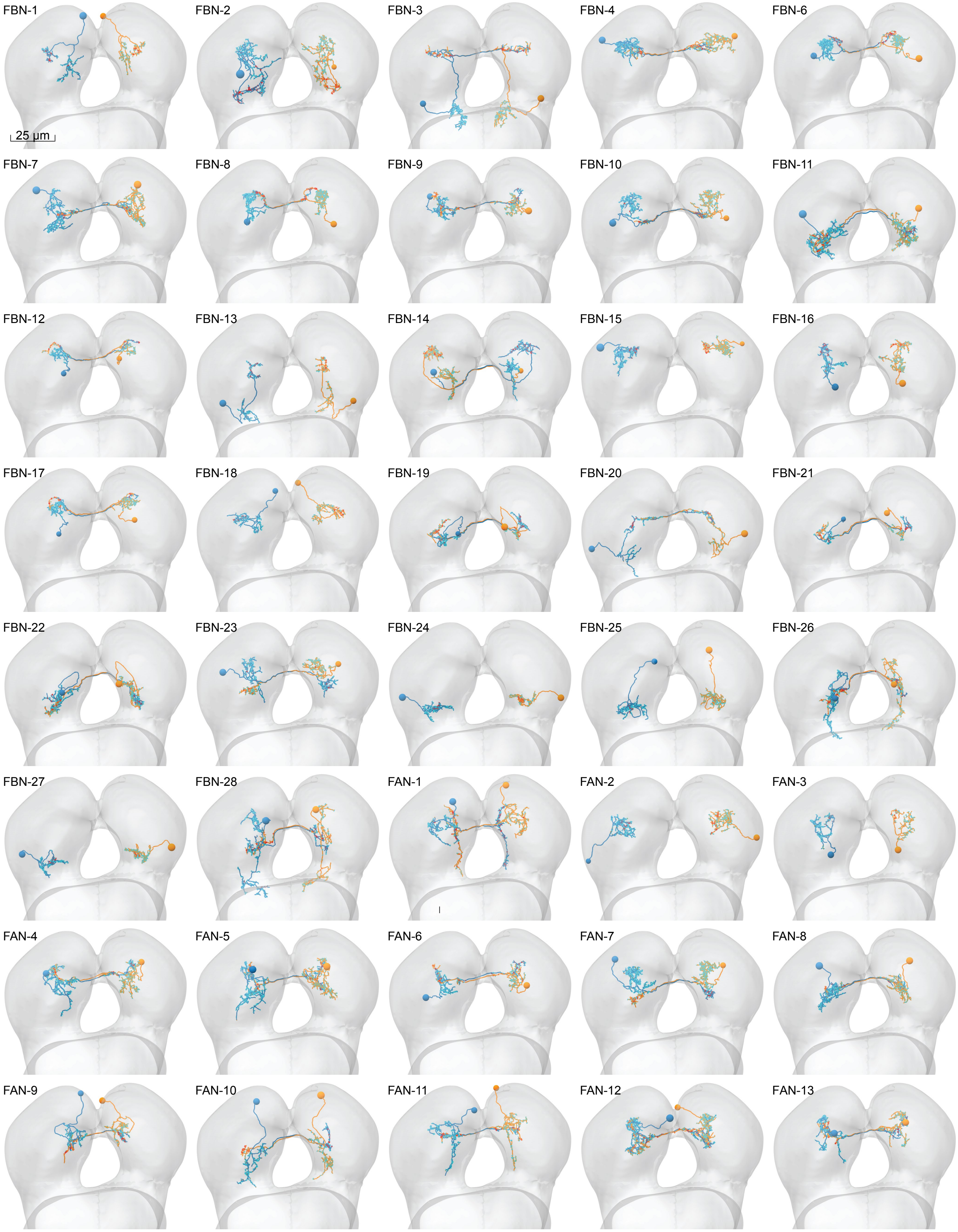

*Continues from prior page.*

## FB2Ns

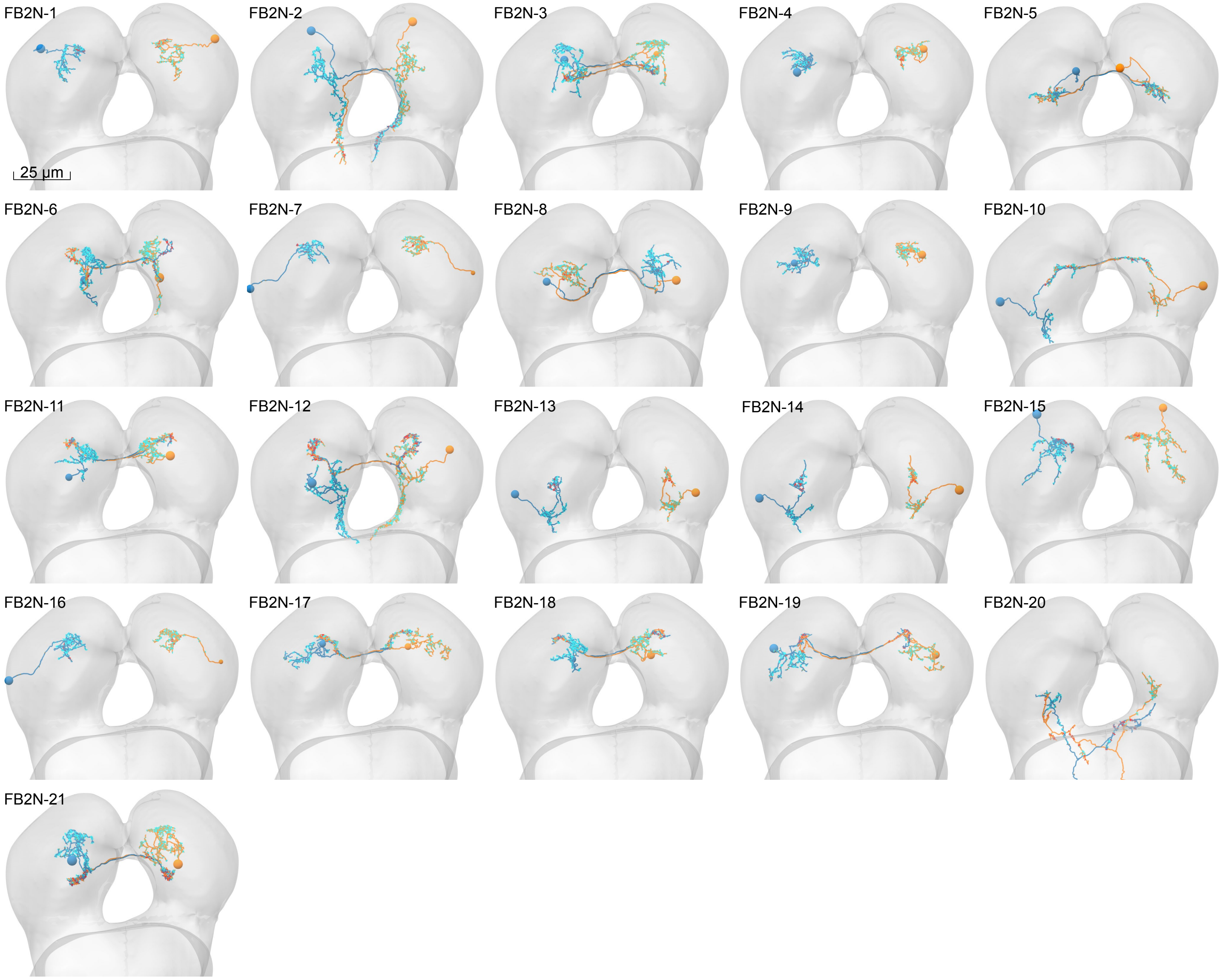


#### FFNs

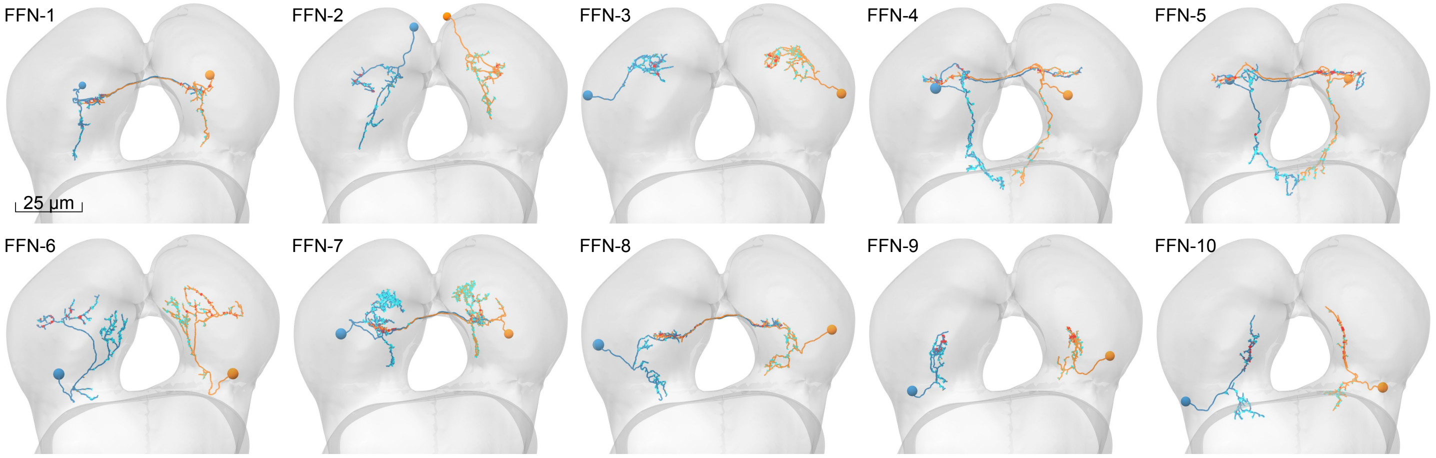

*Continues from prior page.*

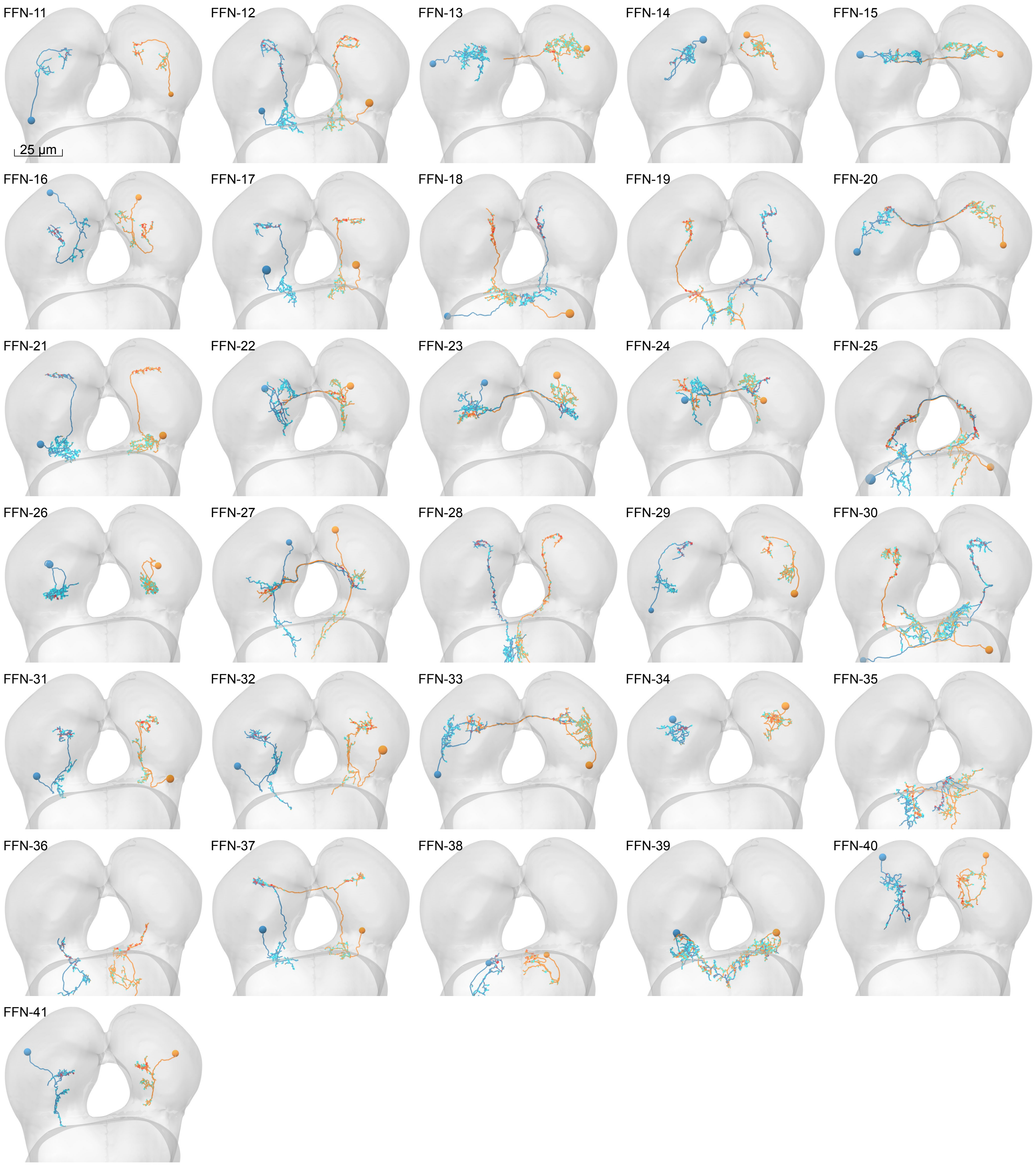


Morphology of all neurons reconstructed in this study. Posterior view of the EM-reconstituted larval central nervous system showing the brain lobes and pairs of homologous neurons.

#### Supplementary Table 1

##### DRIVERS

| | neuron targetted | driver name | AD ∩ | DBD | crossed to effector | figure | ref |
|---|---|---|---|---|---|---|---|
| **Figure 1** | OAN-a1 | *SS24765-Split-GAL4* | *105H06* | *128G03* | *UAS-CsChrimson-mV* | 1c | |
| | DAN-f1 | *SS02180-Split-GAL4* | *76F05* | *129C09* | *UAS-CsChrimson-mV, UAS-GCamp6f* | 1c + 1f | |
| | DAN-f1 | *MB145b-Split-GAL4* | *15B01* | *72B05* | *UAS-CsChrimson-mV* | 1c | |
| | DAN-f1 + DAN-c1 | *MB065b-Split-GAL4* | *14E06* | *72B05* | *UAS-CsChrimson-mV* | 1c | 6 |
| | DAN-g1 | *SS01716-Split-GAL4* | *14E06* | *27G01* | *UAS-CsChrimson-mV, UAS-GCamp6f* | 1c + 1f | 7 |
| | OAN-g1 | *SS04268-Split-GAL4* | *103G03* | *106H09* | *UAS-CsChrimson-mV* | 1c | |
| | DAN-f1 + DAN-g1 | *MB054b-Split-GAL4* | *76F05* | *15B01* | *UAS-CsChrimson-mV* | 1c | |
| | DAN-d1 | *MB143b-Split-GAL4* | *15B01* | *32F01* | *UAS-CsChrimson-mV* | 1c | |
| | DAN-d1 | *MB328b-Split-GAL4* | *82C10* | *32F01* | *UAS-CsChrimson-mV, UAS-GCamp6f* | 1c + 1f | |
| | MBIN-e2 + DAN-c1 | *SS01702-Split-GAL4* | *14E06* | *53C05* | *UAS-CsChrimson-mV, UAS-GCamp6f* | 1c + 1f | |
| | OAN-e1 | *SS01958-Split-GAL4* | *107D05* | *75F01* | *UAS-CsChrimson-mV, UAS-GCamp6f* | 1c + 1f | |
| | DAN-c1 | *SS02160-Split-GAL4* | *52C06* | *53C05* | *UAS-CsChrimson-mV* | 1c | |
| | DAN-h1 + DAN-i1 + DAN-k1 | *SS01948-Split-GAL4* | *105C04* | *107F07* | *UAS-CsChrimson-mV* | 1c | |
| | DAN-i1 | *SS00864-Split-GAL4* | *13D05* | *36B06* | *UAS-Gcamp6f* | 1f | 7 |
| | MD class IV | *ppk1.9-GAL4* | | | *UAS-CsChrimson-mV* | 1e | 8 |
| | Basins 1:4 | *GMR72F11-GAL4* | | | *UAS-CsChrimson-mV* | 1e | 9 |
| | A00c | *GMR71A10-GAL4 x repo-GAL80 ppk-GAL80* | | | *UAS-CsChrimson-mV* | 1e | 9, 11, 13 |
| | A00c | *SS00883-Split-GAL4* | *71A10* | *91E03* | *UAS-CsChrimson-mV* | 1e | |
| | chordotonal neurons | *iav-LexA in attP40* | | | *LexAop-CsChrimson-tdT* | 1f | 9 |
| | MD class IV | *ppk-1kb-hs43-lexA-GAD* in *attP2* | | | *LexAop-CsChrimson-mV* | 1f | 10 |
| | Basins 1:4 | *GMR72F11-LexA* in *JK22* | | | *LexAop-CsChrimson-tdT* | 1f | 9 |
| | Control for GAL4 | *y w;;attP2* | | | *UAS-CsChrimson-mV* | 1e | 4,5 |
| | Control for Split-GAL5 | *y w;attP40;attP2* | | | *UAS-CsChrimson-mV* | 1e | 4,5 |
| **Figure 4** | DAN-h1 + DAN-i1 + DAN-j1 | *58E02-LexAp65* in *attP40* | | | *LexAop-GCamp6f* | 4c + 4f | 13 |
| | MBON-m1 | *SS02163-Split-GAL4* | *52H01* | *40F09* | *UAS-CsChrimson-mCh* | 4c | |
| | MBON-i1 | *SS01726-Split-GAL4* | *20C05* | *14C08* | *UAS-CsChrimson-mCh* | 4f | |
| **Figure 5** | FB2IN-19 | *SS02401-Split-GAL4* | *121D03* | *125G01* | *UAS-CsChrimson-mV* | 5f' | |
| | FB2IN-11 + FB2IN-18 | *SS01778-Split-GAL4* | *78G04* | *22E12* | *UAS-CsChrimson-mV* | 5g' | |
| | FB2IN-11/-18 + MB2IN-207 | *SS02181-Split-GAL4* | *78G04* | *119C11* | *UAS-CsChrimson-mV* | 5g' | |
| | FAN-7 + MB2ON-86 | *SS02108-Split-GAL4* | *13D05* | *40F09* | *UAS-CsChrimson-mV* | 5h' | |
| | MB2ON-86 | *SS04330-Split-GAL4* | *117H07* | *40F09* | *UAS-CsChrimson-mV* | 5h' | |

*Continues from prior page.*

#### EFFECTORS

| | experiment | effector name | short name | figure | ref |
|---|---|---|---|---|---|
| **Figure 1, 5** | Optogenetic activation for behavior | *20XUAS-CsChrimson-mVenus* in *attP18* | *UAS-CsChrimson-mV* | 1c + 5f | 1 |
| | Optogenetic activation for electrophysiology | *20xUAS-CsChrimson-mCherry* in *su(Hw)attP1* | *UAS-CsChrimson-mCh* | 4 | 1 |
| **Figure 1** | Optogenetic activation of Basins or cho for imaging | *13XLexAop2-CsChrimson-tdTomato* in *VK00005* | *LexAop-CsChrimson-tdT* | 1f | 1 |
| **Figure 1** | Optogenetic activation of MD IV for imaging | *13XLexAop2-CsChrimson-mVenus in attP40* | *LexAop-CsChrimson-mV* | 1f | 1 |
| **Figure 1** | Imaging of calcium response to Basins or cho | *20xUAS-IVS-GCaMP6f 15.693*[2] in *attP2* | *UAS-GCamp6f* | 1f | 2 |
| **Figure 1** | Imaging of calcium response to MD IV | *20xUAS-IVS-GCaMP6f*[2] in *VK00005* | *UAS-GCamp6f* | 1f | 2 |
| **Figure 4** | Electrophysiology | *13xLexAop2-IVS-GCaMP6f-p10 50.693*[2] in *VK00005* | *LexAop-GCamp6f* | 4 | 2 |


1 Klapoetke, N. C. *et al.* Independent optical excitation of distinct neural populations. *Nat Methods* **11**, 338-346, doi:10.1038/nmeth.2836 (2014).

2 Chen, T. W. *et al.* Ultrasensitive fluorescent proteins for imaging neuronal activity. *Nature* **499**, 295-300, doi:10.1038/nature12354 (2013).

3 Luan, H., Peabody, N. C., Vinson, C. R. & White, B. H. Refined spatial manipulation of neuronal function by combinatorial restriction of transgene expression. *Neuron* **52**, 425-436, doi:10.1016/j.neuron.2006.08.028 (2006).

4 Pfeiffer, B. D. *et al.* Refinement of tools for targeted gene expression in Drosophila. *Genetics* **186**, 735-755, doi:10.1534/genetics.110.119917 (2010).

5 Jenett, A. *et al.* A GAL4-Driver Line Resource for Drosophila Neurobiology. *Cell Rep* **2**, 991-1001, doi:10.1016/j.celrep.2012.09.011 (2012).

6 Aso, Y. *et al.* The neuronal architecture of the mushroom body provides a logic for associative learning. *Elife (Cambridge)* **3**, e04577, doi:10.7554/eLife.04577 (2014).

7 Saumweber, T. *et al.* Functional architecture of reward learning in mushroom body extrinsic neurons of larval Drosophila. *Nature communications* **9**, 1104, doi:10.1038/s41467-018-03130-1 (2018).

8 Ainsley, J. A. *et al.* Enhanced locomotion caused by loss of the Drosophila DEG/ENaC protein Pickpocket1. *Current biology : CB* **13**, 1557-1563 (2003).

9 Ohyama, T., Schneider-Mizell, C. *et al.* A multilevel multimodal circuit enhances action selection in Drosophila. *Nature* **520**, 633-639, doi:10.1038/nature14297 (2015).

10 Vogelstein, J. T. *et al.* Discovery of brainwide neural-behavioral maps via multiscale unsupervised structure learning. *Science* **344**, 386-392, doi:10.1126/science.1250298 (2014).

11 Yang, C. H. *et al.* Control of the postmating behavioral switch in Drosophila females by internal sensory neurons. *Neuron* **61**, 519-526, doi:10.1016/j.neuron.2008.12.021 (2009).

12 Awasaki, T., Huang, Y., O'Connor, M. B. & Lee, T. Glia instruct developmental neuronal remodeling through TGF-beta signaling. *Nature neuroscience* **14**, 821-823, doi:10.1038/nn.2833 (2011).

13 Liu, C. *et al.* A subset of dopamine neurons signals reward for odour memory in Drosophila. *Nature* **488**, 512-516, doi:10.1038/nature11304 (2012).


Description of the genetic lines used in this study. The GAL4/UAS, Split-GAL4/UAS, and LexA/LexAop lines used in the corresponding figures and allowing expression of channelrhodopsins or reporters in identified neurons.

**Supplementary Adjacency Matrix**

Adjacency matrix between all neurons reconstructed in this study. See csv file.
